# A resistance-gene-directed tolerance trait and selective inhibitors proffer HMG-CoA reductase as a new herbicide mode of action

**DOI:** 10.1101/2022.04.18.488698

**Authors:** Joel Haywood, Karen J. Breese, Jingjing Zhang, Mark T. Waters, Charles S. Bond, Keith A. Stubbs, Joshua S. Mylne

## Abstract

Decades of intense herbicide use has led to resistance in weeds. Without innovative weed management practices and new herbicidal modes of action, the unabated rise of herbicide resistance will undoubtedly place further stress upon food security. HMGR (3-hydroxy-3-methylglutaryl-coenzyme A reductase) is the rate limiting enzyme of the eukaryotic mevalonate pathway successfully targeted by statins to treat hypercholesterolemia in humans. As HMGR inhibitors have been shown to be herbicidal, HMGR could represent a new mode of action target for the development of herbicides. Here we present the crystal structure of a HMGR from *Arabidopsis thaliana* (AtHMG1) which exhibits a wider active site than previously determined structures from different species. This plant conserved feature enabled the rational design of specific HMGR inhibitors, for which we engineered a tolerance trait through sequence analysis of fungal gene clusters. These results suggest HMGR to be a viable herbicide target modifiable to provide a tolerance trait.

---

As herbicide resistance continues to rise, the efficacy of herbicides has diminished^1^ such that new modes of action are desperately needed. Only one new herbicide mode of action has been brought to market in almost 40 years^2^. Weeds are yet to evolve significant resistance to clomazone and bixlozone^1^, two herbicides that disrupt isoprenoid biosynthesis by targeting the enzyme 1-deoxy-D-xylulose-5-phosphate synthase^3^. Found in all kingdoms of life, isoprenoid biosynthesis is crucial for synthesis of lipids, hormones, vitamins and defence compounds^4–6^. The biosynthetic route differs between kingdoms; most animals, fungi, protists and archaea use a mevalonate (MVA) pathway whereas most Gram-negative bacteria including cyanobacteria use a methylerythritol phosphate (MEP) pathway^7^. Through a shared evolutionary history with cyanobacteria, plants use both pathways^7–11^ compartmentalised to the cytosol (MVA) or plastids (MEP)^12–14^. None of the known modes of action for any of the commercial herbicides affect the MVA pathway^1^. An important enzyme in the MVA pathway is HMGR, which is a highly regulated^15–19^, rate-limiting enzyme of the MVA pathway and is the target of the group of hypercholesterolemia therapeutics known as statins^20,21^. Two classes (I and II) of HMGR have been defined^22^ based on the differences between the catalytic core domain structure^23,24^, the presence of an N-terminal membrane domain of between two (plants) and eight (human) membrane-spanning helices in the majority of class I enzymes^20,25^, and the varied NAD(P)H cofactor preference^26^. HMGR regulation appears to be conserved between humans and plants with the N-terminus regulated by ubiquitination whereas catalytic core activity is regulated by phosphorylation^7^. Many of the regulatory proteins differ however, and this is further complicated by plants having multiple copies or isoforms plus a wide variety of external signals modifying expression, such as light and herbivory^7, 27^.

The first potent statin inhibitor of HMGR discovered was mevastatin, isolated from *Penicillium citrinum* in 1976^28^. Lovastatin, isolated from *Aspergillus terreus* in 1978, became the first commercial statin in 1987^29^. Second-generation statins have been semi-synthetic or synthetic products^29^, but all statins competitively inhibit HMGR via a HMG-like moiety and a variable hydrophobic group that together give affinities to HMGR that are 10,000 fold higher than HMG-CoA^30^. Lovastatin and mevastatin, as well as the semi-synthetic pravastatin and synthetic atorvastatin are all known to be herbicidal^31–33^. HMGR might have been overlooked as a herbicide target due to potential off-target risks arising from its conservation in humans and the antimicrobial activity of statins^34^, but recently developed selective insecticides against HMGR illustrate the potential to develop HMGR herbicides^35^.

Here we have solved crystal structures for a plant HMGR in *apo* form and complexed with a statin. These structures reveal a wider active site conserved in plants compared to other organisms. By rational design we developed statin derivatives with over 20-fold specificity for the plant over the human enzyme and which importantly retained herbicidal activity. By comparing the AtHMG1 structure to fungal *HMGR* genes in biosynthetic clusters for natural statins, we demonstrated a single amino acid change confers statin tolerance *in vitro* and *in planta*. Together these findings suggest HMGR is a viable target for herbicide development.

## Results

### Statins range in herbicidal activity

Previous studies have shown several statins to exhibit herbicidal activity against several plant species including *Lemna gibba, Raphanus sativus*, *Scoparia dulcis* and *A. thaliana*^31–33, 36, 37^. However, there is a lack of comparative data regarding the herbicidal efficacy of statins especially for second-generation, synthetic statins. To assess herbicidal activity, we treated a model dicot and a monocot (*A. thaliana* and *Eragrostis tef*, respectively) with a dose range of eight commercially available statins on soil, pre- and post-emergence (**Fig. 1**). All statins were more herbicidal against the dicot and in general were more effective post-emergence. In line with their physicochemical properties more closely matching those of post-emergence herbicides (**Supplementary Fig. 1a)**. The synthetic statin rosuvastatin was the most herbicidal statin being lethal to *A. thaliana* at ∼15 μM without formulation beyond including a wetting agent. Given that under the same conditions formulated glyphosate (Roundup®) is lethal at ∼35 μM (**Supplementary Fig. 1b, c**) we surmised that HMGR could represent a novel herbicide target.

**Figure 1.**
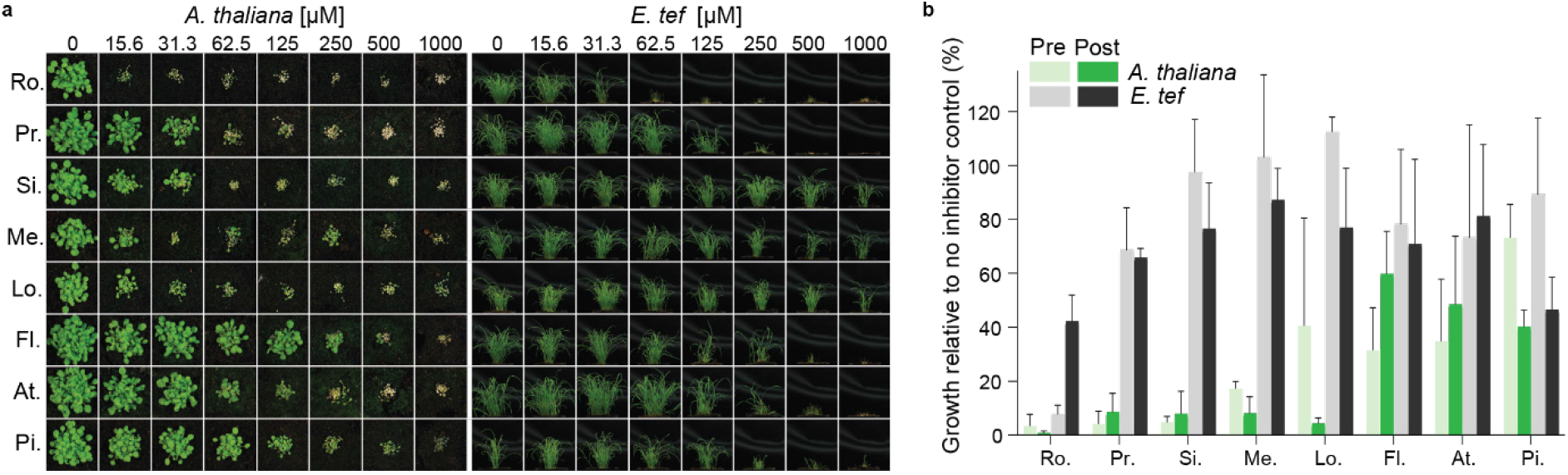
Herbicidal activity of statins varies between a model dicot and a monocot. **a**, Representative images from post-emergence treatment of a model dicot, *A. thaliana*, and pre-emergence treatment of the monocot *E. tef*, with statins: rosuvastatin (Ro.), pravastatin (Pr.), simvastatin (Si.), mevastatin (Me.), lovastatin (Lo.), fluvastatin (Fl.), atorvastatin (At.) and pitavastatin (Pi.). **b**, *A*. *thaliana* (green) and *E*. t*ef* (grey) treated with a range of statins at 62.5 μM pre- and post-emergence on soil. Inhibition was quantified using green pixel area and plotted as a percentage of no-inhibitor control. *n* = 3 replicates with the mean ± standard deviation (s.d.).

### Crystal structure of AtHMG1 reveals scope for species selective compounds

*A. thaliana* has two *HMGR* genes with different expression patterns, but *AtHMG1* (At1g76490) is the most highly epxressed^38^. The N-terminal transmembrane domains of HMGR are highly divergent between species and absent from class II HMGRs (**Supplementary Fig. 2**). By contrast, the conserved extracellular domain of AtHMG1 shares ∼54% sequence identity with HsHMGCR and strictly conserved catalytic residues (**Supplementary Fig. 2**). To develop plant-specific statins and mitigate off-target effects, we solved the crystal structure of the core domain of *apo* AtHMG1 and in complex with pitavastatin to resolutions of 1.9 and 2.1 Å, respectively, in space group *I* 4_1_ 2 2. Attempts were made to crystallise type I statins as described previously^24^, however electron density for these ligands was ambiguous. The structure of the *apo* AtHMG1 displayed a single monomer in the asymmetric unit which through crystallographic symmetry forms a homotetrameric assembly (**Fig. 2a**), consisting of two canonical class I homodimeric HMGR folds, with high structural similarity to HsHMGCR (PDB 1HWK, r.m.s.d. 1.1 Å over 371 Cα atoms, **Fig. 2b**).

**Figure 2.**
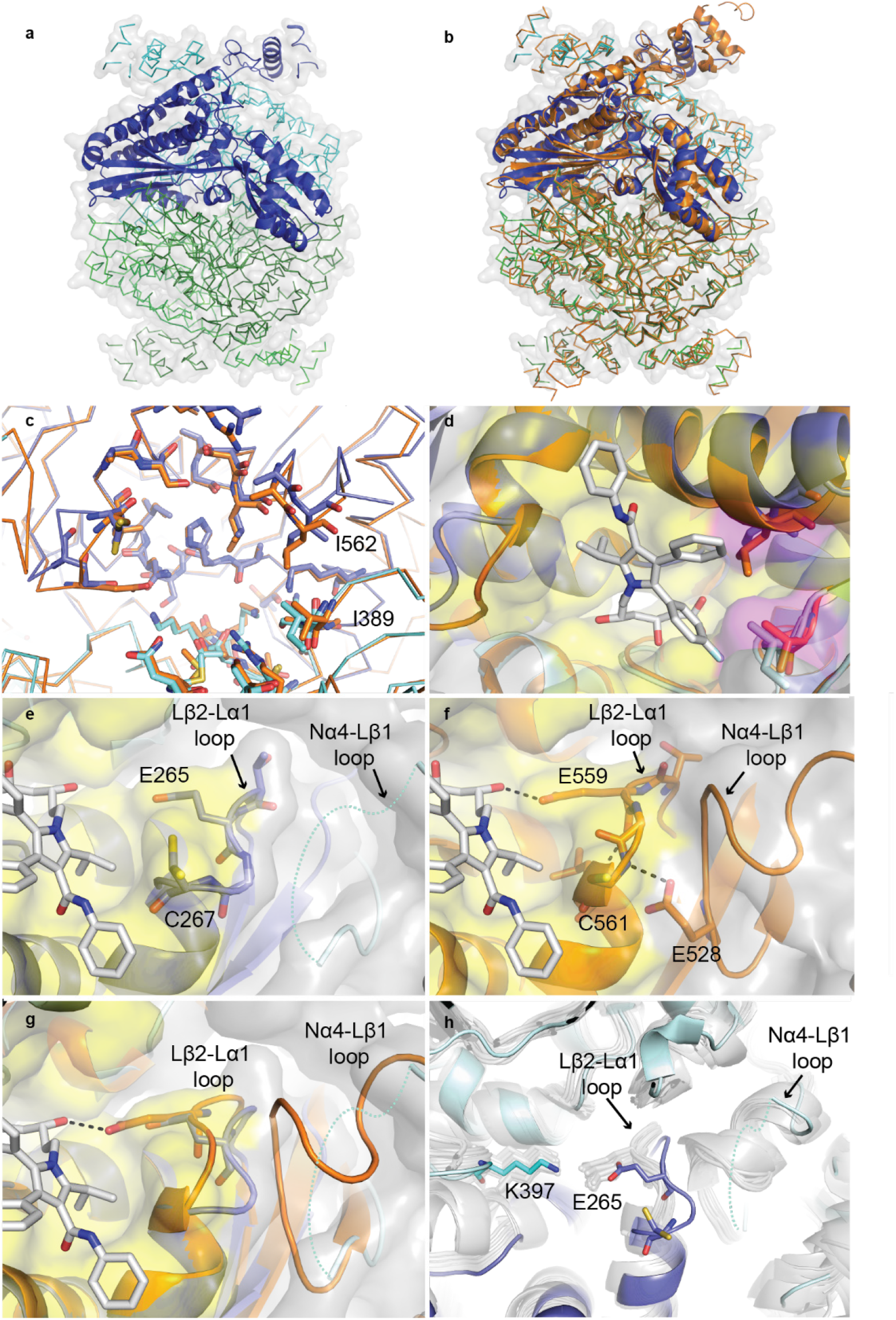
AtHMG1 active site adopts a unique conformation. **a**, *Apo* AtHMG1 displays a single monomer in the asymmetric unit (dark blue cartoon) which through crystallographic symmetry forms a homotetrameric assembly (ribbon) consisting of two canonical class I homodimeric HMGR folds (blue and green). **b**, Overlay of human HMGR (orange cartoon, PDB 1HWK) with *apo* AtHMG1 (**a**) illustrates their conserved fold. **c**, AtHMG1 (blue ribbon and sticks) has a highly conserved active site with HsHMGCR (orange ribbon and sticks). Active site delineating residues are shown as sticks. All residues are conserved except the two AtHMGR1 residues labelled. **d**, Superposition of atorvastatin in the active site of AtHMG1 illustrates the position of these substitutions relative to a bound statin. Conserved active site residues are shown with yellow surface and substitutions highlighted with magenta surface. **e**, Conformational flexibility in the Nα4-Lβ1 loop of AtHMG1 (cyan dotted line) evidenced by poor electron density is likely the result of a Pro to Val substitution. This results in the loss of a type II hydrogen bonded β-turn exhibited in HsHMGCR (**f**) that allows HsHMGCR E559 to hydrogen bond (dashed grey lines) to the open lactone ring of statins. Atorvastatin superimposed on *apo* AtHMG1 (**e**, **g**) illustrates the equivalent residue, E265, likely too far away to H-bond to statins. Conserved active site residues (**e**-**g**) are shown with yellow surface. **h**, Overlay of *apo* AtHMG1 (blue cartoon and sticks) with all published structures of HMGR (grey cartoon and sticks)^24, 26, 30, 40–50^. This conformation has not been seen in any published HMGR crystal structure to date. Topology designation from HsHMGCR^49^.

Closer inspection of the statin-binding pocket revealed two substitutions in AtHMG1 with respect to HsHMGCR located at the hydrophobic CoA binding region of the active site pocket^39^, specifically, Ile^562^/Leu^857^ and Ile^389^/Val^683^ in AtHMG1/HsHMGCR respectively (**Fig. 2c,d**). Furthermore, a plant conserved Val to Pro (Pro^236^/Val^530^ *A. thaliana/*human) substitution at the start of the Lβ1-strand is the likely cause of conformational flexibility and lack of electron density in the Nα4-Lβ1 loop adjacent to the active site-delineating Lβ2-Lα1 loop (**Fig. 2e and Supplementary Fig. 3a, b**). This flexibility results in the loss of a type II β-turn found within the HsHMGCR Lβ2-Lα1 loop that is stabilised by hydrogen bonding between a conserved Glu (Glu^234^/Glu^528^ AtHMG1/HsHMGCR) and a Cys backbone amine (Cys^267^/Cys^561^ AtHMG1/HsHMGCR) (**Fig. 2f**). This altered conformation of the Lβ2-Lα1 loop is not seen in any of the previous class I and II HMGR crystal structures^24, 26, 30, 40–50^ and allows alternative conformations of the Cys^267^ residue (**Fig. 2e,g,h**). Importantly, the arrangement of the AtHMG1 Lβ2-Lα1 loop results in Glu^265^ being unable to form a hydrogen bond with the O5-hydroxyl group of the HMG moiety of statins or the equivalent thioester oxygen of HMG-CoA, as it is shifted 2.5 Å away, creating a wider pocket (**Fig. 2g**). In this orientation it is more likely that Lys^397^ acts as a proton donor in the catalytic reduction of HMG-CoA to mevalonate as is suggested to occur in bacteria^50^ and with molecular dynamics and quantum mechanics/molecular mechanics simulations with HsHMGCR^51^. Together, these differences increase the solvent accessible area of the statin pocket from ∼314 Å^3^ in HsHMGCR to ∼357 Å^3^ in AtHMG1^52^.

The complex of AtHMG1 with pitavastatin (**Fig. 3a,b**) revealed a binding mode highly similar to fluvastatin in HsHMGCR^24^ (**Fig. 3c**). Conserved polar interactions occur with the residues local to the cis loop Arg^296^, Ser^390^, Asp^396^, Lys^397^, Lys^398^, Asn^461^ (HsHMGCR Arg^590^, Ser^684^, Asp^690^, Lys^691^, Lys^692^, Asn^755^) and a salt-bridge between the terminal carboxylate of the HMG moiety with Lys^441^ (HsHMGCR Lys^735^) (**Fig. 3a**). The fluorophenyl group of pitavastatin maintains conserved stacking interactions with Arg^296^ (HsHMGCR Arg^590^) and hydrophobic interactions between the quinoline and cyclopropyl moiety with residues Leu^268^, Ile^389^, Leu^558^, Asp^561^ (HsHMGCR Leu^562^, Val^683^, Leu^853^, Asp^856^). This complex structure however also revealed two notable differences between the binding mode of class II statins in AtHMG1 and HsHMGCR; (i) loss of hydrogen bonding to the O5-hydroxyl group of the HMG moiety of statins from Glu^265^ (HsHMGCR Glu^559^), despite a slight shift of Glu^265^ towards the bound inhibitor (**Fig. 3a, Supplementary Fig. 3c**), and (ii) the loss of hydrophobic interactions with Gly^266^, His^458^ and Ile^562^ (HsHMGCR Gly^560^, His^752^ and Leu^857^). Unique hydrophobic contacts were made between pitavastatin and residues Ser^271^ and Ser^367^ of AtHMG1 (HsHMGCR Ser^565^ and Ser^661^).

**Figure 3.**
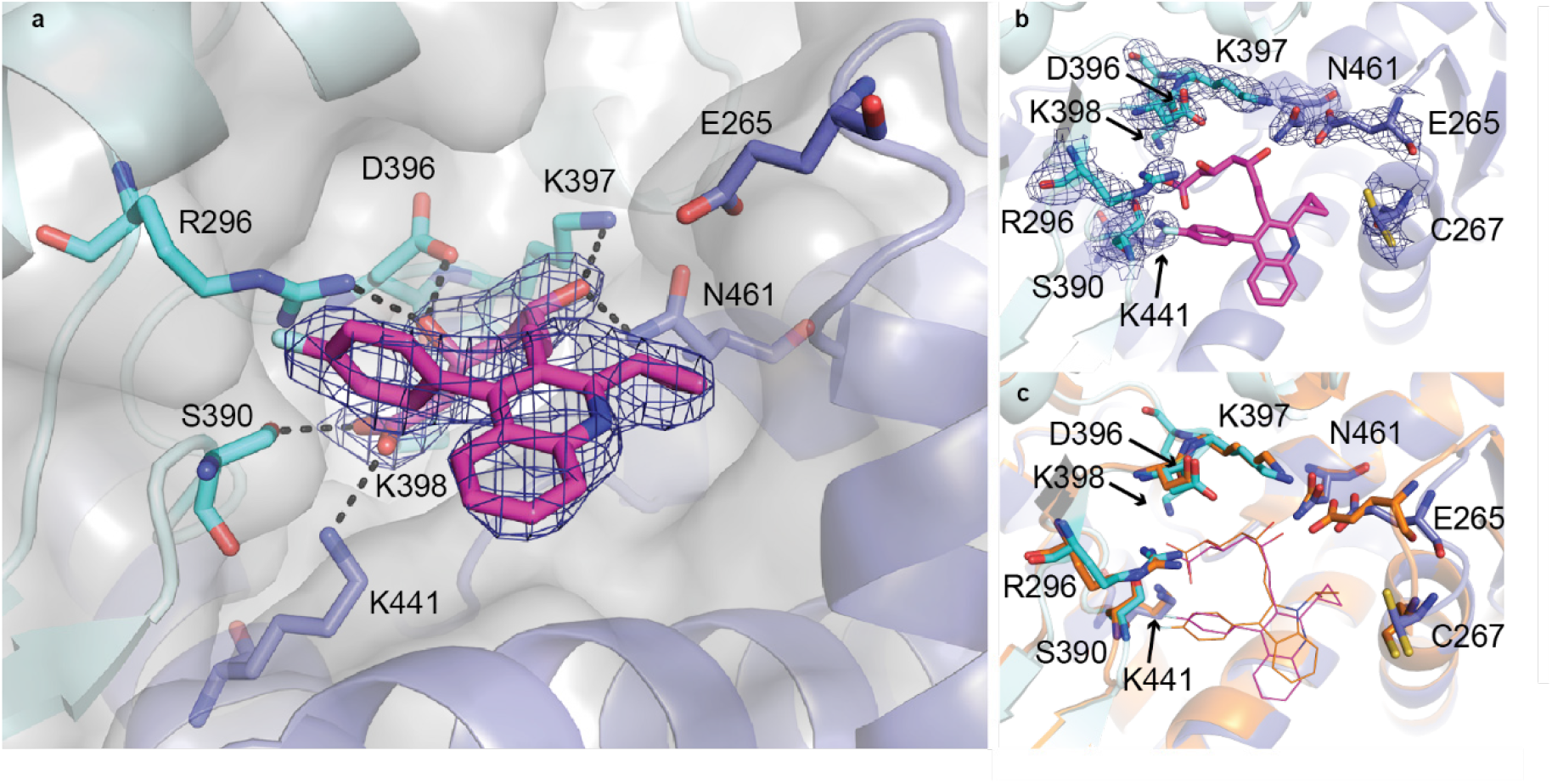
AtHMG1 E265 does not hydrogen bond with statins. **a**, AtHMG1 with pitavastatin (magenta) bound. Residues that hydrogen bond (dashed black lines) to the HMG moiety of statins in AtHMG1 are shown (blue/cyan sticks). **b**, AtHMG1 (blue cartoon) with pitavastatin (magenta line) bound, active site-delineating residues labelled and shown as sticks with electron density. **c**, AtHMG1 with pitavastatin bound superimposed on HsHMGCR bound to fluvastatin (orange cartoon and line PDB 1HWI), illustrating that binding of statins to AtHMG1 is analogous to HsHMGCR. Simulated annealing omit electron density maps (2 F_obs_ - F_calc_) contoured at 1σ level.

### Development of plant-specific analogues of statins

Our insights from the crystal structure of a model plant HMGR and the binding mode of pitavastatin provided the opportunity to rationally design plant-specific inhibitors. To this end, we sought to exploit the Lβ2-Lα1 loop region of AtHMG1 by developing analogues (**1**-**9**) of the more chemically tractable atorvastatin with modifications at the isopropyl group on the central pyrrole ring (**Supplementary Fig. 3d**). Activity of the atorvastatin scaffold against HsHMGCR was previously found to be reduced with increasing size of the alkyl substituent^53^ whereas the wider pocket of AtHMG1 might accommodate larger groups. Additionally, the loss of interactions with O5-hydroxyl group of the HMG moiety with Glu^265^ could be targeted by incorporating a hydrogen bond donor (**Fig. 2e,g**). Thus **1**-**9** were synthesised (**Supplementary Experimental**) and assessed for herbicidal activity on soil with *A. thaliana* (**Fig. 4a**) and for species-specificity against HsHMGCR and AtHMG1 *in vitro* by a fluorometric, NADPH-depletion assay (**Fig. 4b-d**).

**Figure 4.**
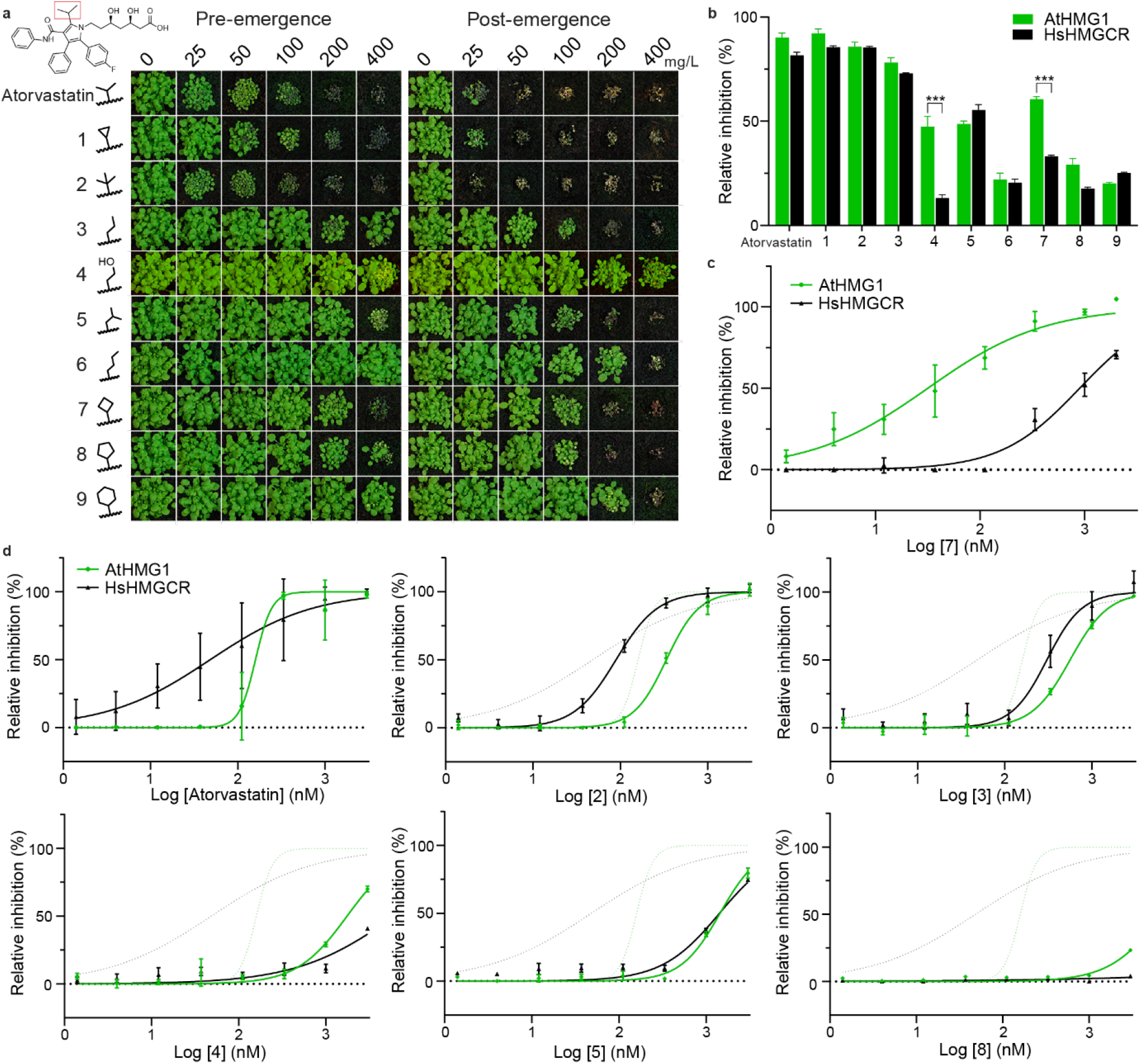
Modifying the isopropyl group of atorvastatin affects species selectivity. **a**, Herbicidal activity of atorvastatin and its analogues (**1**-**9**) against *A. thaliana* with pre- and post-emergence treatments. The isopropyl moiety of atorvastatin is boxed in red. Modifications to the isopropyl region are shown. **b**, Compounds **4** and **7** were selective *in vitro* for AtHMG1 over HsHMGCR at 500 nM. *n* = 3 independent reactions with the mean ± s.d. Significance from unpaired t-test P ≤ 0.001. **c**, *In vitro* inhibition of AtHMG1 and HsHMGCR by **7** illustrating >20-fold selectivity for AtHMG1. *n* = 3 independent reactions with the mean ± s.d. **d**, Atorvastatin and analogues **2-5** and **8** were not selective for AtHMG1 *in vitro*. Inhibition of AtHMG1 and HsHMGCR with atorvastatin inhibition profile shown as dotted lines. *n* = 3 independent reactions with the mean ± s.d. except for a single point (AtHMG1 333 µM At. *n* = 2).

Overall, atorvastatin analogues with side chains of similar length had similar herbicidal and *in vitro* inhibitory activity to the parent, whereas side chains longer than the isopropyl group had reduced activity (**Fig. 4**). Compounds **4** and **7** retained herbicidal activity and displayed a preference for AtHMG1 over HsHMGCR in an initial screen (**Fig. 4a,b**). Dose response curves confirmed compound **7** had switched preference from human to plant when compared to atorvastatin, showing >20-fold higher specificity for AtHMG1 (IC_50_ 32 nM ±12 nM) over HsHMGCR (IC_50_ 890 nM ±143 nM) *in vitro* (**Fig. 4c,d**). These molecules provide a framework for the future development of plant-specific HMGR inhibitors that might exhibit stronger herbicidal activity.

### Exploiting biosynthetic gene clusters to engineer statin tolerance

The most commercially successful herbicides are often paired with a tolerance trait in crops. Statins produced from fungal biosynthetic gene clusters usually contain a copy of *HMGR* that imparts self-resistance^54–56^, so we sought to determine the structural basis for this resistance. Sequence alignment of a *HMGR* gene (*lvrA*) from the *A. terreus* genome revealed several mutations in the cluster-associated copy that were not present in the housekeeping copy (**Supplementary Fig. 4**). The corresponding residues of the AtHMG1 crystal structure revealed a Leu (Leu^558^) to Thr mutation, whose equivalent was conserved in all *A. terreus* genomes in the NCBI database. The Leu to Thr mutation would likely disrupt the hydrophobic pocket essential for accommodating the decalin ring of natural statins (**Fig. 5a**), and so was incorporated into recombinant AtHMG1. The AtHMG1-L558T mutant was resistant to a range of statins (**Fig. 5b**) with >20-fold resistance to rosuvastatin *in vitro* (WT IC_50_ 53 nM ±20 nM, L558T IC_50_ >1000 nM) (**Fig. 5c**). Without inhibitors, AtHMG1-L558T had reduced catalytic activity (WT *K_m_* 69 µM ±16 µM and k*_cat_* 21.3 ±1.7 s^-1^, L558T *K_m_* 24 µM ±14 µM and k*_cat_* 4.8 ±0.6 s^-1^) (**Supplementary Fig. 5**), but remained within the range of previously published rates for other class I and II HMGR enzymes^57^.

**Figure 5.**
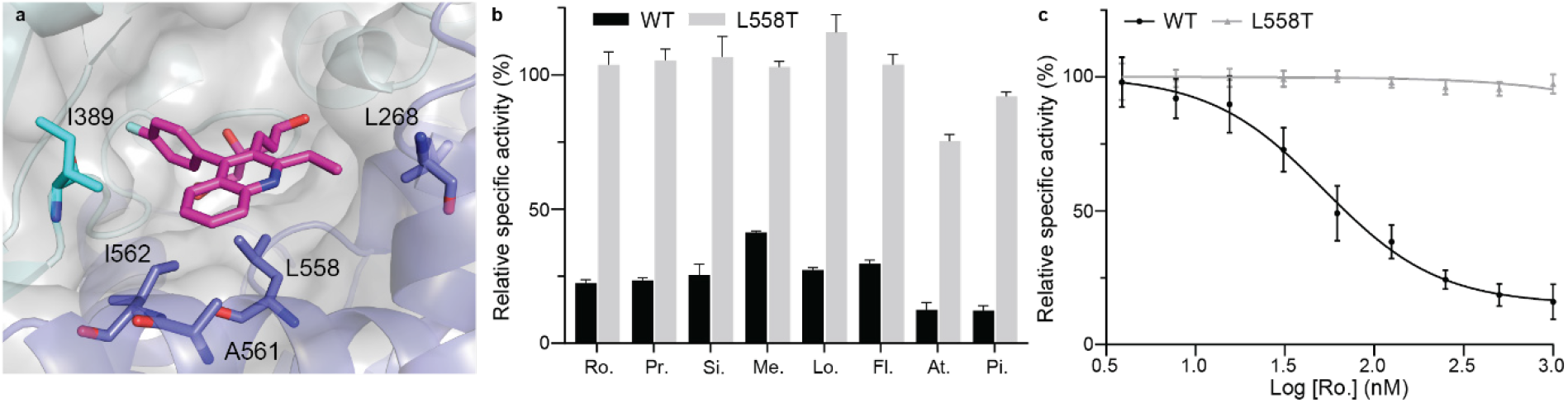
A mutation found in a statin biosynthetic cluster confers statin resistance *in vitro*. **a**, The hydrophobic pocket in AtHMG1 delineated by labelled residues (blue sticks) with pitavastatin (magenta sticks) bound, illustrating L558 proximity to the hydrophobic ring of statins. **b**, AtHMG1 with the L558T mutation (grey bar) retained activity *in vitro* in the presence of statins: rosuvastatin (Ro.), pravastatin (Pr.), simvastatin (Si.), mevastatin (Me.), lovastatin (Lo.), fluvastatin (Fl.), atorvastatin (At.) and pitavastatin (Pi.), at 500 nM. *n* = 3 independent reactions with the mean ± s.d. **c**, *In vitro* inhibition of WT and L558T AtHMG1 by rosuvastatin revealed the L558T mutation conferred >20-fold resistance. *n* = 3 independent reactions with the mean ± s.d.

To validate the potential of the L558T mutation for providing a plant tolerance trait we overexpressed full length AtHMG1 (*35S::AtHMG1*) and its equivalent with the L558T mutation (*35S::AtHMG1-L558T*) in *A. thaliana*, using a cauliflower mosaic virus (CaMV) *35S* promoter. It has previously been shown that overexpressing *AtHMG1* in *A. thaliana* can give rise to a 40-fold rise in mRNA levels and a modest rise in resistance to lovastatin compared to non-transformed WT controls^58^. Here we found with data collated from 19 independent T_2_ *35S::AtHMG1* lines and 14 independent *35S::AtHMG1-L558T* T_2_ lines that both constructs conferred similar resistance to the selectable marker hygromycin (**Fig. 6a,d**). However, the *35S::AtHMG1-L558T* lines were over six-fold more resistant to rosuvastatin (IC_50_ 300 µM vs ±18 µM) than *35S::AtHMG1* lines (IC_50_ 46 µM ±5 µM) and more than 100-fold more resistant than non-transformed WT (IC_50_ 3 µM vs ±1 µM) (**Fig. 6b,d**).

**Figure 6.**
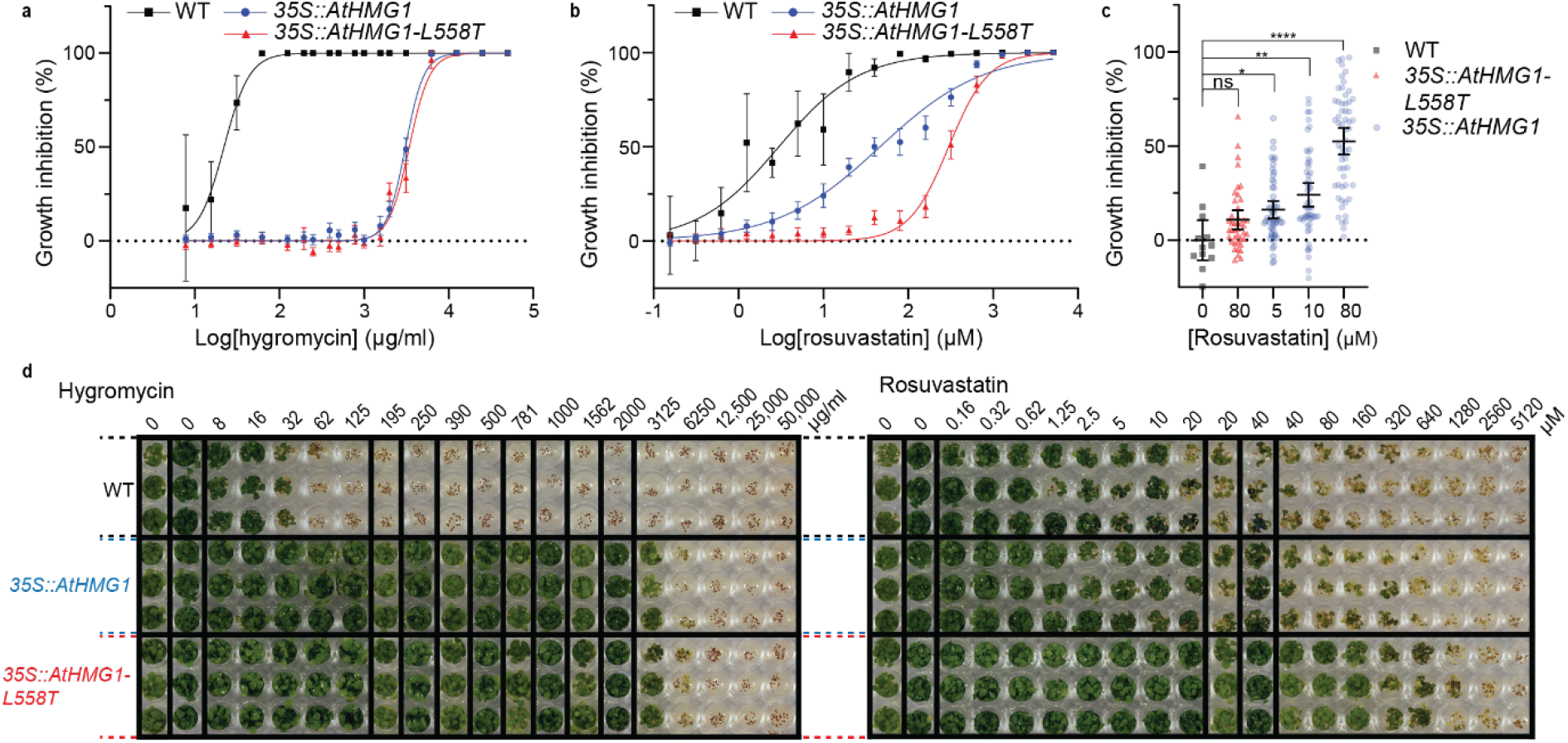
The L558T mutation gives resistance to rosuvastatin *in planta*. Resistance to hygromycin (**a**) and rosuvastatin (**b** and **c**) in 19 transgenic lines of *35S::AtHMG1* (blue) versus 14 *35S::AtHMG1-L558T* lines (red) and wild type (WT, black). Green pixels quantified and plotted as a percentage of no-inhibitor control. **a**, Both transgenic lines exhibited similar resistance to the hygromycin selectable marker, whereas WT was sensitive, mean ± 95% CI. **b**, *35S::AtHMG1-L558T* transgenic lines were six-fold more resistant to rosuvastatin than *35S::AtHMG1*, mean ± 95% CI, but ± s.d. for WT. **c**, Susceptibility of transgenics to rosuvastatin illustrated *35S::AtHMG1-L558T* (80 µM *n* = 42) was up to 16-fold less susceptible to rosuvastatin inhibition than *35S::AtHMG1* (5 µM *n* = 54, 10 µM *n* = 54) when compared to untreated WT (*n* = 12). Significance from one-way ANOVA, mean ± 95% CI. **d**, Representative image of resistance to hygromycin and rosuvastatin from a single line of *35S::AtHMG1* and *35S::AtHMG1-L558T* versus WT.

Furthermore, analysis of the effects of rosuvastatin revealed *35S::AtHMG1-L558T* lines were up to 16-fold less sensitive to treatment than *35S::AtHMG1* lines (**Fig. 6c**). These results illustrate the potential for HMGR to have a tolerance trait and further validates the *in vitro* results (**Fig 5b,c**).

## Discussion

The relentless rise in herbicide resistant weeds already poses a significant threat to global food security and as such, new herbicides with new modes of action are desperately needed. Moreover, as consumer attitudes shift, natural product ‘bioherbicides’ will rise in their appeal and currently in the USA enjoy an accelerated regulatory journey^59^.

Herein, we have validated HMGR as a potential new herbicide target. Using the HMGR crystal structure from a model plant we have demonstrated that, despite its overall sequence and structure conservation with HsHMGCR, differences in the architecture (especially the active site) can be exploited to develop plant-specific synthetic HMGR inhibitors. The progress herein provides a basis for the discovery of new natural product statins that might be suitable bioherbicides.

The differences in the architecture of AtHMG1 that allowed for species-selectivity largely arise from an unusual orientation of the Lβ2-Lα1 loop that is likely the result of increased flexibility in the neighbouring Nα4-Lβ1 loop. The atypical orientation of the Lβ2-Lα1 loop in AtHMG1 disrupts the hydrogen bonding network formed between the catalytic residues Glu^265^/Lys^397^/Asn^461^/Asp^473^ (HsHMGCR Glu^559^/Lys^691^/Asn^755^/Asp^767^), thereby retaining only those hydrogen bonds that stabilise the catalytic Lys via the adjacent Asn and Asp residues. The conserved location of the catalytic Lys between AtHMG1 and other class I and II HMGRs strongly suggests this residue is responsible for polarising the carbonyl oxygen of HMG-CoA substrate and mevaldehyde intermediate, and for performing the final protonation step. Glu^265^ is not in a favourable position to hydrogen bond to either the substrate thioester oxygen or the adjacent Asp^473^, which based on HsHMGCR *in silico* simulations (HsHMGCR Glu^559^ and Asp^767^) might be expected to hydrogen bond and stabilise the mevaldyl-CoA intermediate^51^. Further molecular dynamics studies with AtHMG1, its substrate and cofactors might determine if the role of Glu^265^ is to hydrogen bond to Asp^473^, or to directly protonate the substrate as previous modelling studies have suggested^60,61^.

Here we rationally designed a compound with >20-fold preference for plant HMGR *in vitro* with limited modification to the parent scaffold. Modelling of atorvastatin along with compounds **4** and **7** into the active site of HsHMGCR revealed a single dominant high affinity binding mode with a large drop in affinity to the next most favourable binding mode. Atorvastatin exhibited the highest affinity followed by compound **7** and **4** (**Supplementary Fig. 6**) consistent with *in vitro* results (**Fig. 4**). Modelling with AtHMG1 revealed more varied poses of the analogues, with similar affinities between the most favourable binding modes. These binding modes are possibly facilitated by a wider active site and flexibility in the Lβ2-Lα1 loop region (**Supplementary Fig. 3c**) and might account for the difficulty we had in obtaining co-crystal structures for AtHMG. The lower affinity for AtHMG1 than HsHMGCR for compounds **4** and **7** suggests further molecular dynamics simulations and crystallographic studies may be necessary to help reveal the molecular basis of *in vitro* specificity (**Fig. 4**). Notably, selectivity over HsHMGCR was also obtained by targeting the same Lβ2-Lα1 loop region in *Manduca sexta* using *gem*-difluoromethylenated HMGR inhibitors. Similar derivatives may also prove to be selective for plant HMGRs. Overall, the developed compounds provide a framework for further structure-based rational herbicide design targeting the Lβ2-Lα1 loop region of AtHMG1, which could be validated for selectivity in mammalian *in vivo* studies. Greater species selectivity might be obtainable by targeting the N-terminal domain of HMGR, which is highly divergent between humans and plants and is absent from class II HMGRs. A recent crystal structure of the regulatory elements that interact with the N-terminal domain in HsHMGCR and studies of compounds that increase HMGR degradation suggest that this could be an alternative mechanism to lower cholesterol levels^16, 18^. Future studies of the regulatory elements interacting with plant HMGR N-terminal domain and complexed crystal structures might in the same way also provide an avenue to develop more species-specific inhibitors of HMGR. The regulatory elements that control plant lipid metabolism might also provide new herbicidal targets, just as the proprotein convertase subtilisin/kexin type 9 and angiopoietin-like 3 are providing new avenues for the treatment of hypercholesterolemia^62, 63^.

By analysing the AtHMG1 crystal structure and sequences in fungal biosynthetic gene clusters, we identified a mutation conferring statin resistance without adversely affecting catalytic activity. Overexpressing this mutant protein in *A. thaliana* demonstrated its potential as a tolerance trait, but further investigations are needed. These could include (i) the efficacy of this protein mutant in different species; (ii) optimisation of expression and regulation, by modifying the N-terminal domain; (iii) its effects on sterol levels and seed set^64^; and (iv) determining what HMGR inhibitor residues remain in the treated crop or soil. Future studies might also focus on other residues that potentially impart resistance, such as the end region of the Sβ4 strand (residues 387-390), that show conservation in putative resistance genes from *Penicillium citrinum* and *Xylaria grammica* and could affect binding of the butyryl group of natural statins. We envisage that the development and discovery of new, natural product herbicides^65, 66^ might also benefit from a similar approach to engineering resistance alleles from biosynthetic gene clusters containing compounds or targets of interest.

## Methods

### Herbicidal activity assay

Approximately 30 seeds of *A. thaliana* (accession Col-0) or *E. tef* were sown in 63 x 63 x 59 mm pots of Irish peat (Bord na Móna Horticulture Ltd, Newbridge, Ireland). Seeds were incubated in the dark for 3 days at 4°C to synchronise germination. A single pre-emergence treatment (day 0) was performed when these seeds were transferred to a growth room at 22°C with a 16:8 hr light:dark photoperiod and 60% relative humidity. Two post-emergence treatments were performed following emergence of the seedlings (day 1) at days 4 and 7. Plants were watered accordingly throughout the experiment to maintain adequate moisture and photographed on day 16. Treatments were conducted with rosuvastatin, simvastatin, fluvastatin, atorvastatin (AK Scientific), pravastatin (BOC Sciences), lovastatin (Sapphire Bioscience), mevastatin, pitavastatin (Focus Bioscience), glyphosate, RoundUp^®^ and atorvastatin analogues (**1**-**9**). To treat, 0.5 mL of each compound in a final concentration of 2% dimethyl sulfoxide (DMSO) and 0.02% Brushwet (SST Dandenong, Australia) was pipetted onto seedlings. For atorvastatin analogue **1-9** synthesis and spectroscopic data see **Supplementary Information**. Growth inhibition was quantified by detecting green pixels for healthy seedlings using ImageJ and the ‘Threshold Colour’ plug-in with the following settings: hue 50-110, saturation 125-255, brightness 30-255 as described previously^67^. Data were normalised to a negative control to provide percentage inhibition.

### HMG-CoA reductase expression and purification

An *E. coli* codon-optimised DNA sequence encoding the conserved extracellular region of AtHMG1 (Uniprot P14891, At1g76490, residues 121-592) was cloned into pQE30 (Qiagen) following an N-terminal His_6_-tag and tobacco etch virus cleavage site. The protein was expressed in the T7 SHuffle Express strain of *E. coli* (New England Biolabs) transformed with pREP4 (Qiagen) with the proteins expressed and purified as previously described^68^. Briefly, cultures were grown in lysogeny broth containing 100 μg/mL ampicillin and 35 μg/mL kanamycin at 30°C to an OD_600_ of 0.8–1.0. Cells were cooled to 16°C before expression was induced with 0.1 mM isopropyl β-D-1-thiogalactopyranoside. Following an overnight culture, cells were harvested by centrifugation and lysed by ultrasonication in 100 mM HEPES (pH 7.5), 150 mM sodium chloride, 5 mM dithiothreitol, 0.1% Triton X-100. Lysed cells were then centrifuged (15,000 × g) and the supernatant was incubated in 30 mL batches with Ni-NTA resin overnight at 4°C. The resin was then washed with 50 mL of 100 mM HEPES (pH 7.5), 150 mM sodium chloride, 5 mM dithiothreitol followed by 50 mL of 100 mM HEPES (pH 7.5), 150 mM sodium chloride, 5 mM dithiothreitol, 20 mM imidazole. The protein was then eluted with 50 mL of 100 mM HEPES (pH 7.5), 150 mM sodium chloride, 5 mM dithiothreitol, 300 mM imidazole. Eluted protein was concentrated with a 30 kDa centrifugal filter unit (Millipore) and purified by size-exclusion chromatography (HiLoad 16/600 Superdex 200) in 100 mM HEPES (pH 7.5), 150 mM sodium chloride, 5 mM dithiothreitol. The protein was assessed for purity by SDS-PAGE and protein concentration determined by spectrophotometry.

### *In vitro* HMGR assay

AtHMG1 described above and human HMGR (HsHMGCR, Uniprot P04035, residues 441-888, cloned as above) were purified as above and activity was determined by spectrophotometric measurement of the decrease in absorbance at 340 nm that occurs with NADPH oxidation in the presence of substrate HMG-CoA (Sigma-Aldrich). Reactions were performed with an assay buffer consisting of 150 mM sodium chloride, 5 mM dithiothreitol, 50 mM HEPES pH 7.4 and 2% DMSO. For kinetics determinations a final concentration of 150 nM enzyme was incubated at 37°C in 300 μM NADPH and different concentrations of HMG-CoA. Nonlinear regression analysis was performed with GraphPad Prism 9 by plotting the initial reaction rates, *v*_0_, interpolated from a standard curve against the substrate concentration. The Michaelis-Menten constant, *K_m_*, was determined by fitting the data with a Michaelis-Menten equation and values for *k_cat_* were calculated by dividing *V_max_* by the molar enzyme concentration. To calculate relative specific activity, 500 nM of test compounds were pre-incubated at 37°C with enzyme and 300 μM NADPH for 15 minutes before adding HMG-CoA to 200 μM. Resultant values were background subtracted and normalised to the average of the no-inhibitor control. For IC_50_ determinations the same protocol was used, but varying inhibitor concentrations. Rosuvastatin data were plotted with a four-parameter (WT) and normalised response (L558T) nonlinear regression model and atorvastatin and analogues were plotted with a normalised response with variable slope nonlinear regression model.

### Crystallisation and data collection

The C-terminal core residues of AtHMG1 (residues 121-576) were cloned and purified as above. The core domain was concentrated to 10-15 mg/mL and used immediately for crystallisation. Crystal screening was performed with 96-well Intelli-Plates (Hampton Research) with 80 μL of reservoir solution using the sitting-drop vapour diffusion method at 16°C. Crystals were obtained with a mother-liquor of 0.2 M ammonium sulphate, 0.1 M HEPES and 35% w/v poly(acrylic acid sodium salt) 2100 from the Molecular Dimensions MIDASplus^TM^ screen. Crystals were optimised using a 96-well additive screen and well-diffracting crystals obtained in the same plates with a 1 μL droplet containing 0.6 μL of the above mother-liquor, 0.3 μL of protein and 0.1 μL of Hampton Research additive screen. Crystals used for inhibitor soaks were grown with the additives 40% v/v pentaerythritol ethoxylate (3/4 EO/OH) and 0.1 M iron(III) chloride hexahydrate. Inhibitor soaks were carried out in the same mother-liquor with 1 mg/mL of inhibitors. Single crystals were quickly soaked in mother-liquor containing 25% glycerol as a cryoprotectant before being flash frozen and stored in liquid nitrogen. Data collection was performed at 100 K on the Australian MX2 (micro-focus) beamline^69^ with 1.9 Å resolution for the *apo* form and 2.1 Å for inhibitor complexed AtHMG1.

### Crystal structure determination, refinement and model building

*Apo* and complexed AtHMG1 diffraction data were processed using XDS and scaled with AIMLESS from CCP4^70, 71^. A sequence alignment of AtHMG1 and HsHMGCR was generated using ClustalO and used to create a search model of AtHMG1 based on the last common atom of PDB 1HW8. This model was then used for molecular replacement with PHASER from CCP4^72^. Manual building and refinement was performed in iterative cycles with Coot and REFMAC5 using the CCP4 program suite^73^. Structure analysis and validation were carried out with Coot and MolProbity^74^. The refined AtHMG1 structure was then used as a search model for molecular replacement with data from inhibitor complexed crystals. Crystallographic data and refinement statistics are summarised in **Table 1** with Ramachandran plot values calculated from CCP4. Coordinates and structure factors were deposited into the PDB under accession code 7ULI and 7ULM. Figures illustrating the structures were generated using PyMol.

**Table 1:** Crystallography data collection and refinement statistics. . Numbers in parentheses refer to the highest resolution bin.

| **Data collection** | *apo* HMG1 | HMG1-pitavastatin |
| --- | --- | --- |
| Space group | *I* 4<sub>1</sub> 2 2 | *I* 4<sub>1</sub> 2 2 |
| Unit cell dimensions |  |  |
| a, b, c (Å) | 85.58, 85.58, 266.65 | 85.55, 85.55, 265.15 |
| $\alpha, \beta, \gamma$ (°) | 90.00, 90.00, 90.00 | 90.00, 90.00, 90.00 |
| Wavelength | 0.9537 | 0.9537 |
| Resolution (Å) | 1.7 | 2.1 |
| R<sub>merge</sub> (%) | 11 (434) | 25 (267) |
| *I*/σ*I* | 14.8 (1.63) | 12 (1.1) |
| Completeness (%) | 100 (99.9) | 60.9 (11.6) spherical<br>91.6 (60.7) ellipsoidal |
| Redundancy | 13.3 (11.6) | 18.2 (14) |
| CC <sub>1/2</sub> | 1.00 (0.549) | 0.996 (0.341) |
| **Refinement** |  |  |
| Resolution (Å) | 45.30-1.90 | 45.07-2.13 |
| No. reflections | 39555 | 17133 |
| R<sub>work</sub>/R<sub>free</sub> | 20.7/24.3 | 22.0/26.1 |
| No. Atoms | 5398 | 5054 |
| Protein | 5307 | 4971 |
| Water | 91 | 19 |
| Ligand |  | 64 |
| Wilson *B* (Å<sup>2</sup>) | 51.0 | 29.7 |
| Average refined B-factor (Å<sup>2</sup>) |  |  |
| Protein only (Å<sup>2</sup>) | 51.0 | 29.3 |
| Water (Å<sup>2</sup>) | 53.5 | 12.8 |
| Ligand (Å<sup>2</sup>) |  | 45.0 |
| r. m. s. deviations: |  |  |
| Bond lengths (Å) | 0.01 | 0.01 |
| Bond angles (°) | 1.36 | 1.55 |
| **Ramachandran analysis** |  |  |
| Favoured (%) | 97 | 94 |
| Allowed (%) | 3 | 6 |
| Outliers (%) | 0 | 0 |
| PDB Accession | 7ULI | 7ULM |

### *In planta* statin resistance assay

DNA encoding the full-length AtHMG1 protein (Uniprot P14891, residues 1-592) and the corresponding L558T mutant were cloned into a derivative of the pMDC43 binary vector^75^ to yield *35S::AtHMG1* and *35S::AtHMG1-L558T* transgenes, respectively. These constructs were then introduced into *Agrobacterium tumefaciens* strain LBA4404 and separately used to transform *A. thaliana* by the floral dip method^76, 77^. Seeds (T_0/1_) of transformed plants were collected and surface sterilised using 600 µL 70% ethanol, 750 µL 100% ethanol and 800 µL 50% bleach before washing with 800 µL sterile water and resuspension with 0.1% agar. Selection was performed on 30 µg/mL hygromycin growth medium (1% agar, 1% glucose, 0.45% Murashige & Skoog salts with vitamins, 0.3% 2-(*N*-morpholino)-ethanesulfonic acid (MES) (v/v), pH 5.7) in a growth room at 22°C with 16:8 hr light:dark photoperiod and 60% relative humidity. Surviving plants were transferred to 63 x 63 x 59 mm pots of Irish peat and grown to maturity in the same growth conditions. Seeds from plants with an adequate seed yield were then sterilised and selected again as described above with 30 µg/mL hygromycin growth medium. Seeds (T_2_) from 22 lines of *35S::AtHMG1* and 15 lines of *35S::AtHMG1-L558T* plants that exhibited approximately 3:1 segregation ratio of hygromycin resistant:sensitive were then sown (∼15 seeds/well, n = 3 replicates), along with wild type (WT) *A. thaliana*, on sterile 96-well microplates with 0.25 mL/well growth medium containing a low dose range serial dilution of 8-2000 µg/mL hygromycin and of 0.16-40 µM rosuvastatin (final concentration 2% DMSO) with respective media only controls, and then again on a second higher dose range of 195-50000 µg/mL hygromycin and of 20-5120 µM rosuvastatin (final concentration 2% DMSO). Plates were sealed with porous tape and grown for a minimum of 10 days with growth conditions described above. Plates were then imaged, and growth quantified using ImageJ (National Institutes of Health, 1.53 v) as described above. Total green pixels were normalised against negative controls for the respective lines (2% DMSO and water) to provide percentage inhibition. Three of 22 lines of *35S::AtHMG1* and 1 of 15 lines of *35S::AtHMG1-L558T* plants were excluded from further analysis based on poor growth of the negative control or for displaying low hygromycin resistance. One of 19 *35S::AtHMG1* lines and 2 of the 14 *35S::AtHMG1-L558T* lines had data only for the higher dose range of hygromycin and rosuvastatin. For IC_50_ determinations all data were respectively combined from 19 lines of *35S::AtHMG1,* 14 lines of *35S::AtHMG1-L558T* and WT *A. thaliana*. Growth inhibition at varying concentrations of hygromycin and rosuvastatin were plotted with a four-parameter non-linear regression model using GraphPad Prism 9.

## Data availability

The refined structural protein models are deposited under PDB accession codes 7ULI (*apo* HMG1) and 7ULM (HMG1-pitavastatin). All structures cited in this publication are available under their respective PDB accession codes. All other raw data are available on request.

## Contributions

J.H. and J.S.M. designed and coordinated research. J.H., J.Z. and K.J.B. performed plant assays. K.J.B. and K.A.S. designed and synthesised atorvastatin analogues. M.T.W. made binary constructs used by J.H. and J.S.M. for plant transgenesis. J.H. analysed transgenic lines, made recombinant proteins, performed assays, and acquired crystals. J.H. and C.S.B. solved crystal structures. J.H. and J.S.M. wrote the manuscript with input from all authors.

## Acknowledgements

The authors thank Grishma Vadlamani and Yit-Heng Chooi for helpful comments. This research was undertaken in part using the MX2 beamline at the Australian Synchrotron, part of The Australian Nuclear Science and Technology Organisation, and made use of the Australian Cancer Research Foundation detector. J.H. was supported by an Australian Research Council Discovery Early Career Researcher Award (DE180101445) and funded in part by Nexgen Plants. This work was supported by an Australian Research Council Discovery Project DP190101048 to J.S.M., K.A.S. and J.H. and an ARC Linkage Infrastructure Equipment and Facilities Grant (LE190100123) to K.A.S.

## Supplementary Information

**Supplementary Table 1:** HMGR sequences used for sequence conservation analysis.

| HMGR in species | Group | % identity to HMGR | Accession code |
| --- | --- | --- | --- |
| *Arabidopsis thaliana* | Eudicot | 100 | P14891.1 |
| *Arabidopsis suecica* | Eudicot | 98 | KAG7659775.1 |
| *Arabidopsis lyrata subsp lyrata* | Eudicot | 98 | XP_002887642.2 |
| *Arabidopsis arenosa* | Eudicot | 98 | CAE5964346.1 |
| *Capsella rubella* | Eudicot | 95 | XP_006302046.1 |
| *Camelina sativa* | Eudicot | 94 | XP_010428712.1 |
| *Eutrema salsugineum* | Eudicot | 92 | XP_006390188.1 |
| *Brassica rapa* | Eudicot | 89 | XP_009128179.1 |
| *Raphanus sativus* | Eudicot | 87 | XP_018445978.1 |
| *Tarenaya hassleriana* | Eudicot | 84 | XP_010537243.1 |
| *Eucalyptus grandis* | Eudicot | 80 | KCW68146.1 |
| *Populus trichocarpa* | Eudicot | 78 | XP_002301898.2 |
| *Ricinus communis* | Eudicot | 78 | XP_002510732.1 |
| *Juglans regia* | Eudicot | 78 | XP_018843042.2 |
| *Lupinus angustifolius* | Eudicot | 78 | XP_019461234.1 |
| *Theobroma cacao* | Eudicot | 78 | EOY15882.1 |
| *Daucus carota* | Eudicot | 78 | XP_017253170.1 |
| *Jatropha curcas* | Eudicot | 77 | XP_012073564.1 |
| *Cannabis sativa* | Eudicot | 77 | XP_030495961.1 |
| *Vitis vinifera* | Eudicot | 77 | XP_002275827.1 |
| *Citrus sinensis* | Eudicot | 76 | XP_006473861.1 |
| *Glycine max* | Eudicot | 75 | XP_003519474.1 |
| *Solanum tuberosum* | Eudicot | 75 | XP_006342182.1 |
| *Rosa chinensis* | Eudicot | 75 | XP_024163983.1 |
| *Medicago truncatula* | Eudicot | 75 | XP_003617066.1 |
| *Gossypium barbadense* | Eudicot | 75 | KAB2042521.1 |
| *Chenopodium quinoa* | Eudicot | 73 | XP_021714065.1 |
| *Brachypodium distachyon* | Monocot | 73 | XP_003572378.1 |
| *Spinacia oleracea* | Eudicot | 72 | XP_021846881.1 |
| *Trifolium subterraneum* | Eudicot | 72 | GAU28089.1 |
| *Ipomoea triloba* | Eudicot | 71 | XP_031109081.1 |
| *Malus domestica* | Eudicot | 71 | XP_008348952.1 |
| *Solanum lycopersicum* | Eudicot | 71 | XP_010317674.1 |
| *Sorghum bicolor* | Monocot | 71 | XP_002445887.1 |
| *Triticum turgidum* | Monocot | 70 | VAI86078.1 |
| *Triticum aestivum* | Monocot | 70 | XP_044433217.1 |
| *Zea mays* | Monocot | 70 | NP_001130818.1 |
| *Oryza sativa* | Monocot | 70 | AAD38873.1 |
| *Setaria viridis* | Monocot | 69 | XP_034581600.1 |
| *Cocos nucifera* | Monocot | 68 | KAG1358991.1 |
| *Pisum sativum* | Eudicot | 68 | AAL37041.1 |
| *Digitaria exilis* | Monocot | 68 | KAF8780957.1 |

**Supplementary Figure 1.**
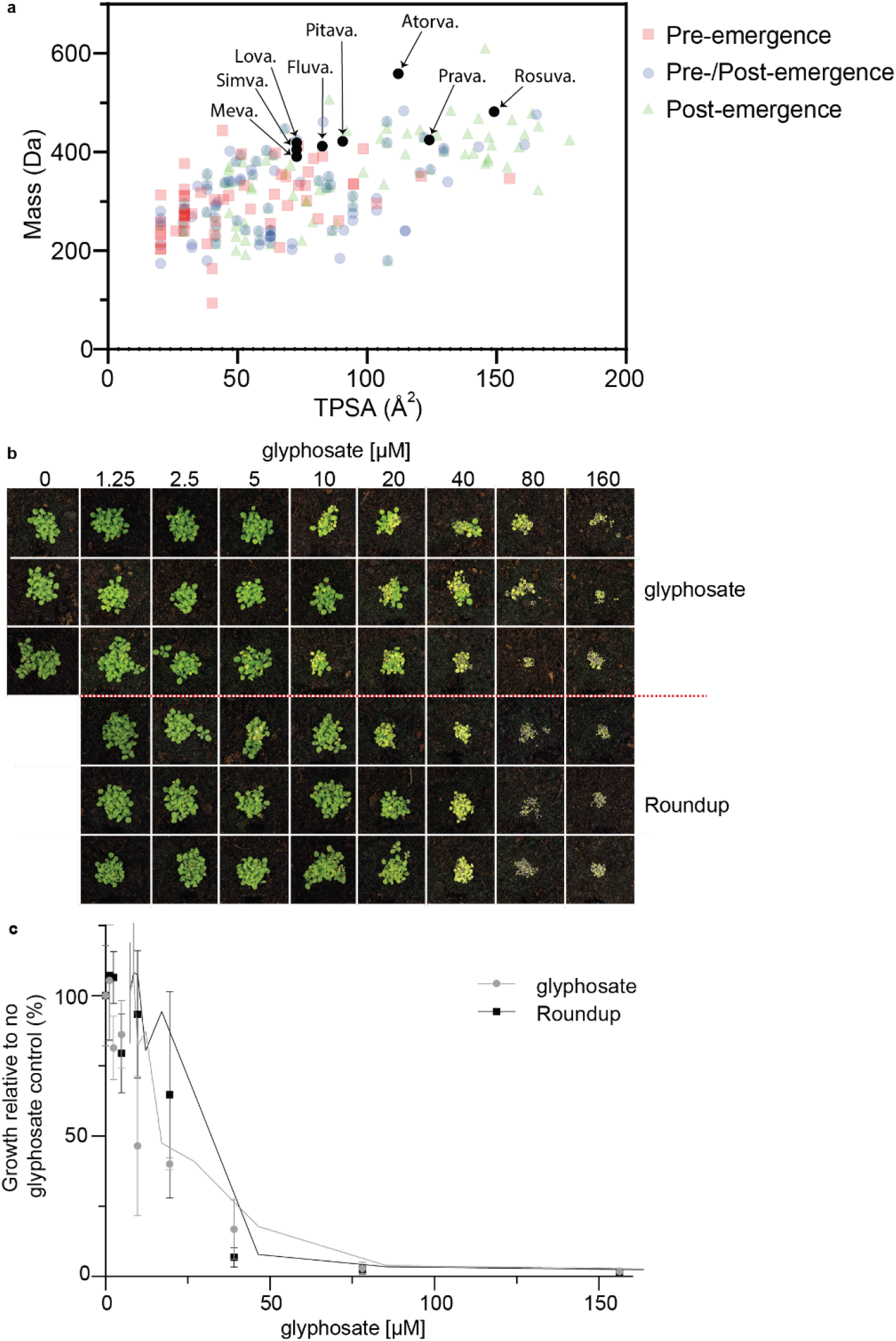
Statins have similar physicochemical properties to post-emergence herbicides and an activity akin to glyphosate. Through analysis of 360 commercial herbicides^78^, we were able to classify 56 as pre-emergence herbicides, 87 as both pre/post emergence herbicides and 104 as post-emergence herbicides. The physicochemical properties of these 247 herbicides were plotted and compared to the physicochemical properties of the commercially available statins (**a**). Pre-emergence herbicides tend to have a smaller mass and smaller topological polar surface area (TPSA) than post-emergence herbicides. (**b, c**) Post-emergence dose range of glyphosate formulated and diluted in 0.02% Brushwet or as Roundup^®^ (360 g/L glyphosate) diluted in water, applied on *A*. *thaliana*. Images taken 12 days post-emergence (**b**) and quantified using ImageJ software (**c**) *n* = 3 replicates with the mean ± standard deviation (s.d.).

**Supplementary Figure 2.**
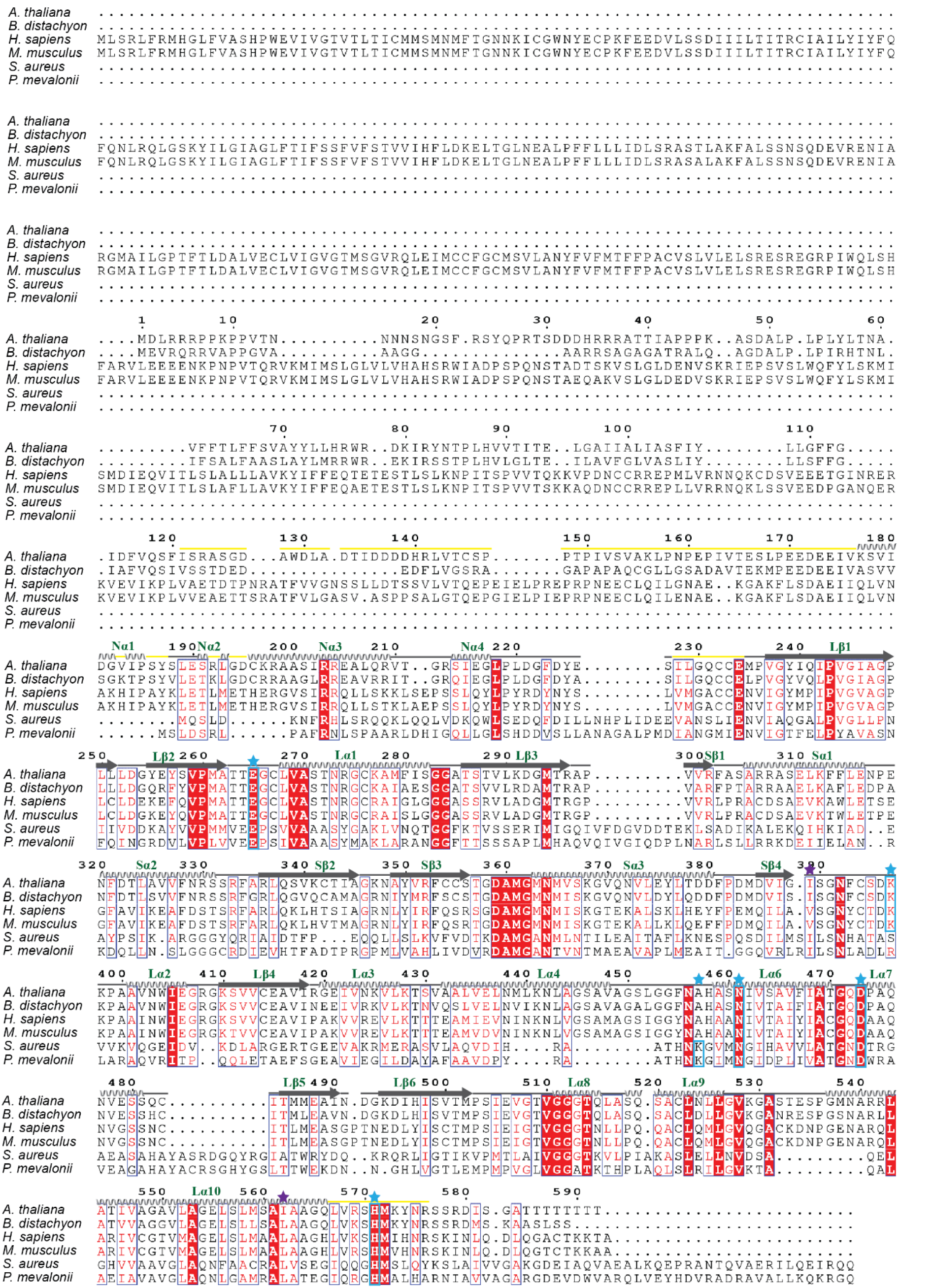
Sequence alignment of AtHMG1 with class I and II HMGRs. Secondary structure elements of AtHMG1 extracellular core domain with labels (green), based on topology designation from human HMGR^49^, are shown above sequence. Solid grey arrows indicate β-strands, grey helices indicate α-helices, grey lines indicate loop regions, yellow lines indicate regions lacking electron density in the *apo* structure. Conserved regions (red box), highly similar residues (red text)^79^. Residues implicated in catalysis from previous studies are highlighted with a cyan star and box. *A. thaliana* active site residues that are divergent from mammalian class I HMGRs are highlighted with a purple star.

**Supplementary Figure 3.**
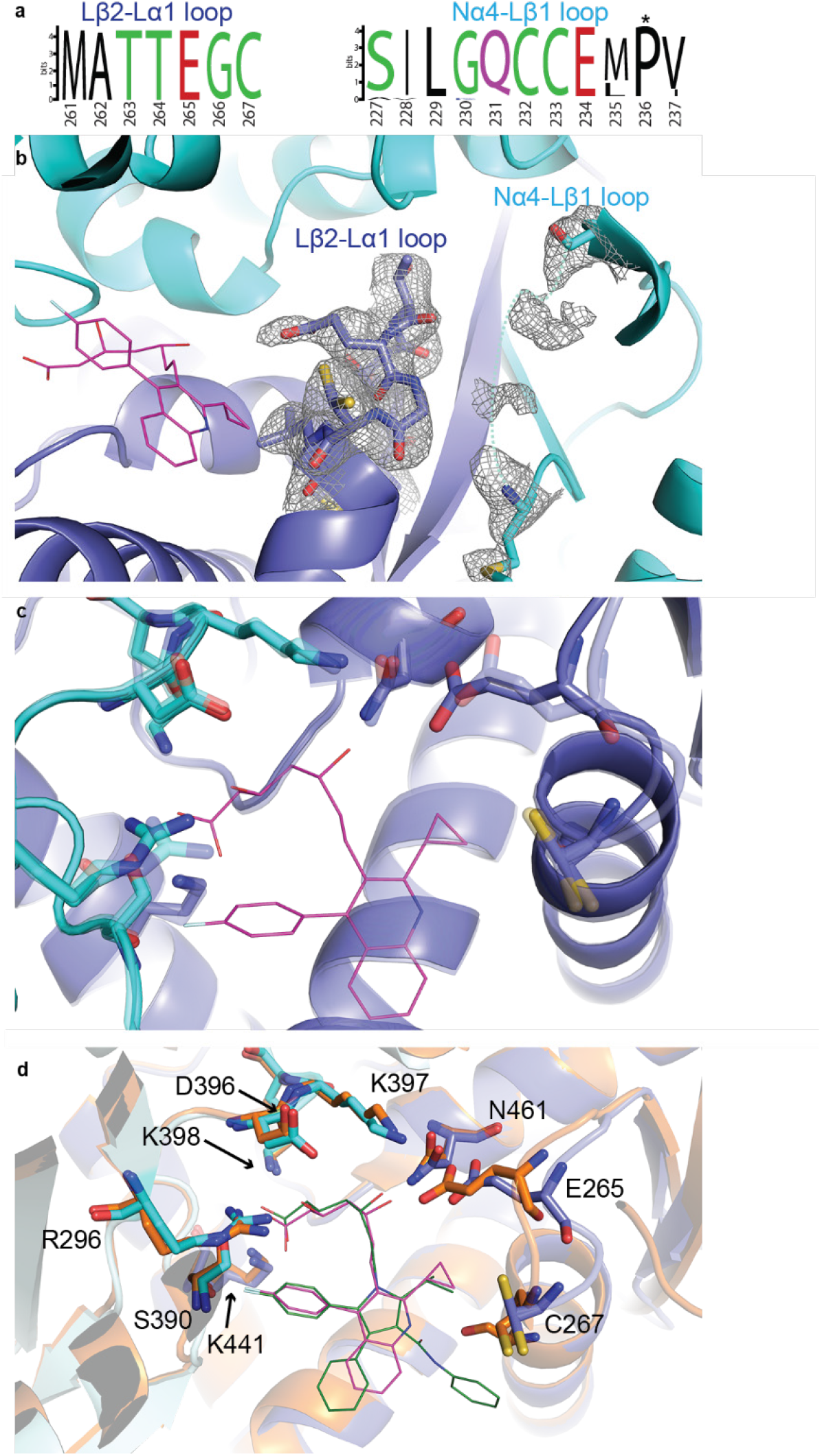
A unique architecture of AtHMG1 could be targeted for the rational design of plant specific inhibitors. (**a**) Relative abundance of residues in the Lβ2-Lα1 and Nα4-Lβ1 loops from 40 plant species (**Supplementary Table 1**), illustrated using the WebLogo server^80^. Pro^236^ (highlighted with an asterisk) is conserved in diverse plant species. (**b**) Simulated annealing omit electron density maps (2 Fobs - Fcalc) contoured at 1σ level illustrate defined density for the apo AtHMG1 Lβ2-Lα1 loop and poorly defined density for the adjacent Nα4-Lβ1 loop. Pitavastatin (magenta line) is superimposed for reference. (**c**) Overlay of Apo (transparent, blue cartoon) and pitavastatin bound (blue cartoon) AtHMG1 active site delineating residues reveals a highly similar overall architecture and a slight shift of Glu^265^ towards the bound inhibitor. (**d**) HsHMGCR1 (PDB 1HWK, orange cartoon) superimposed onto AtHMG1 pitavastatin complex shows the isopropyl group on the central pyrrole ring of atorvastatin (green line) could be modified to target the unique architecture of AtHMG1 Lβ2-Lα1 loop region.

**Supplementary Figure 4.**
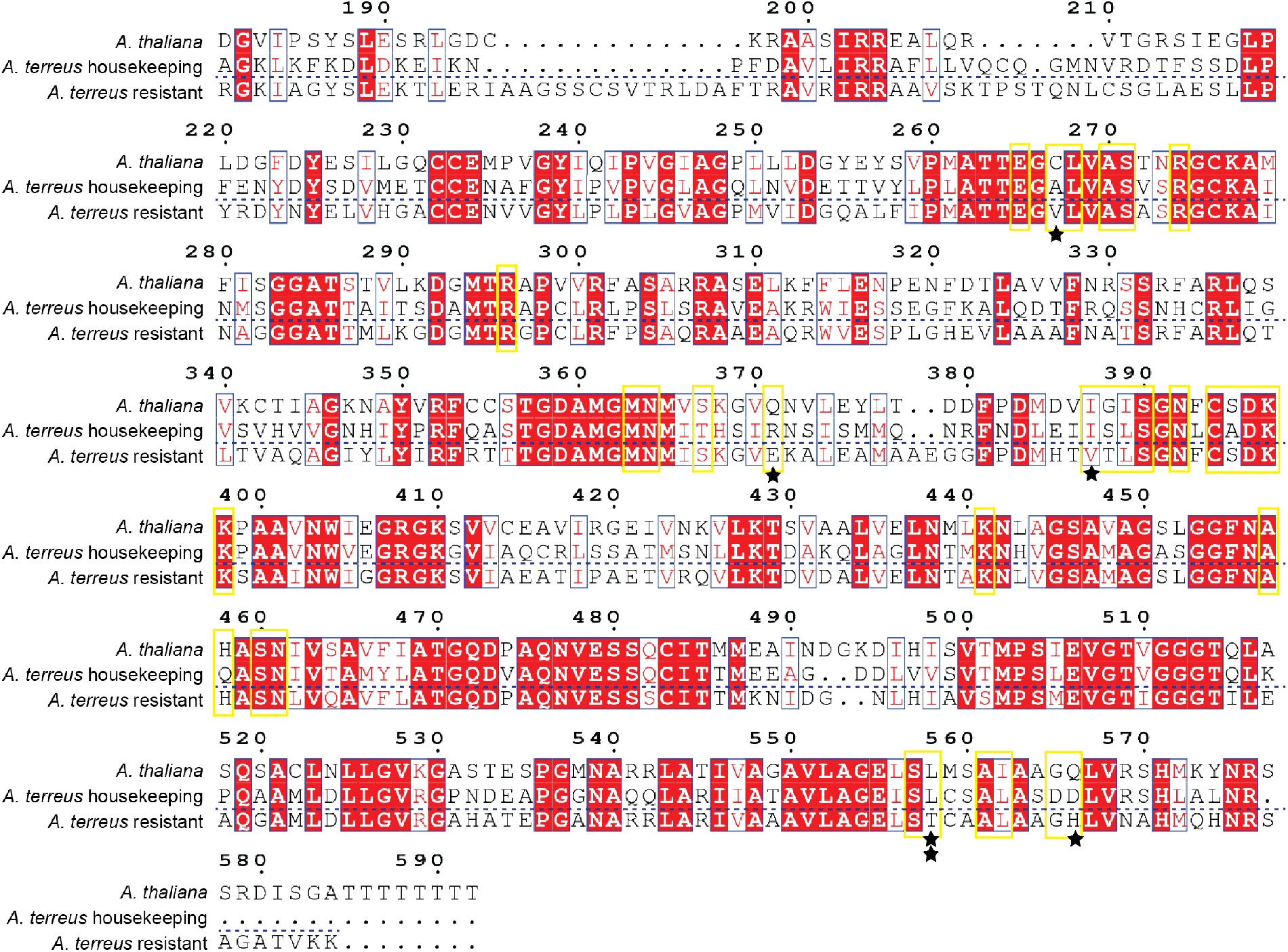
Sequence alignment of AtHMG1 with *Aspergillus terreus* HMGR. Comparison of the sequences of AtHMG1 and *A*. *terreus* NIH2624 putative *HMGR* housekeeping gene (ATEG_02145) with *A*. *terreus* NIH2624 reported HMGR self-resistant gene^54-56^(ATEG_09965) from the lovastatin biosynthetic gene cluster reveals several residues potentially conferring HMGR with statin resistance HMGR (black stars). AtHMG1 active site-delineating residues shown with yellow boxes. Sequence numbers shown for AtHMG1 residue L558 is highlighted with two stars. Conserved regions (red box), highly similar residues (red text)^79^.

**Supplementary Figure 5.**
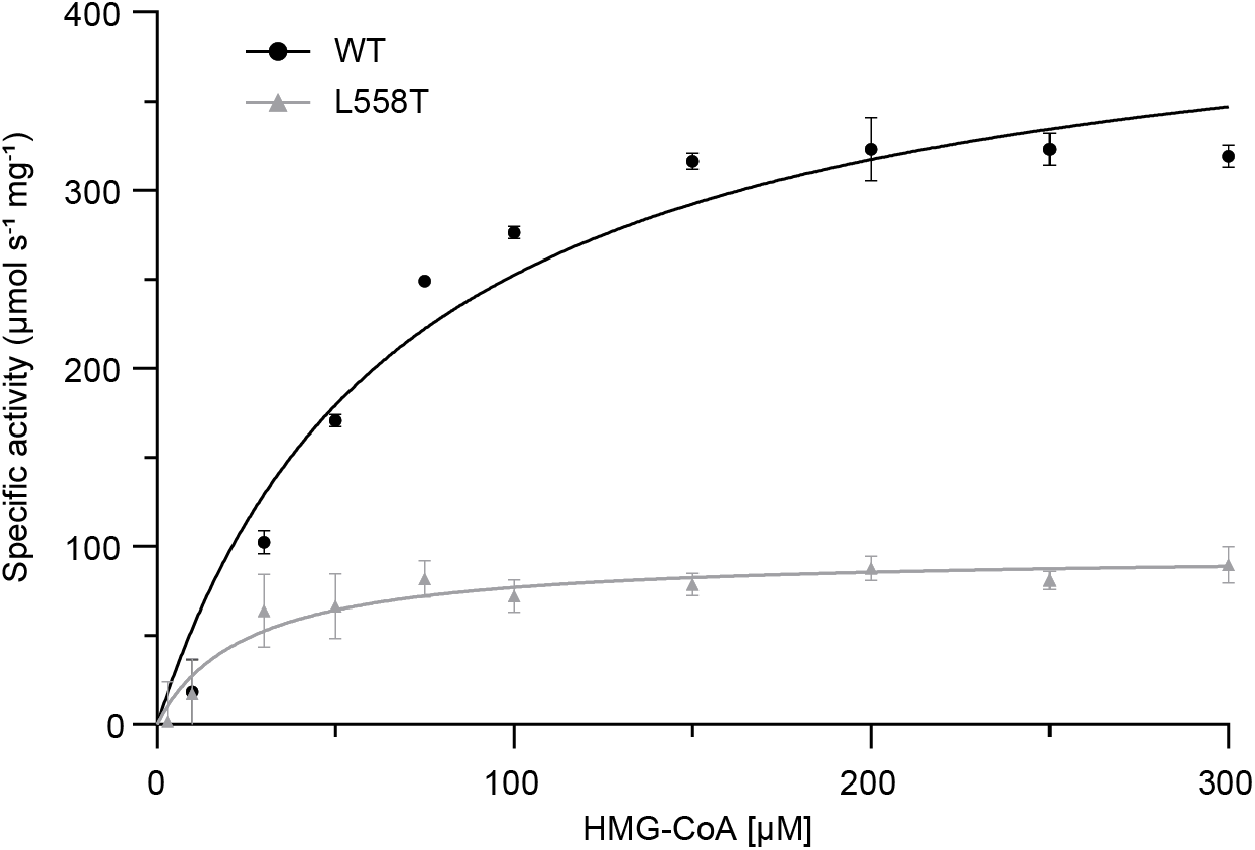
Steady state kinetic data for AtHMG1 and L558T mutant. Substrate HMG-CoA saturation curve for WT (black circles) and L558T mutant (grey triangles) with Michaelis-Menten fit. *n* = 3 independent reactions with the mean ± s.d.

**Supplementary Figure 6.**
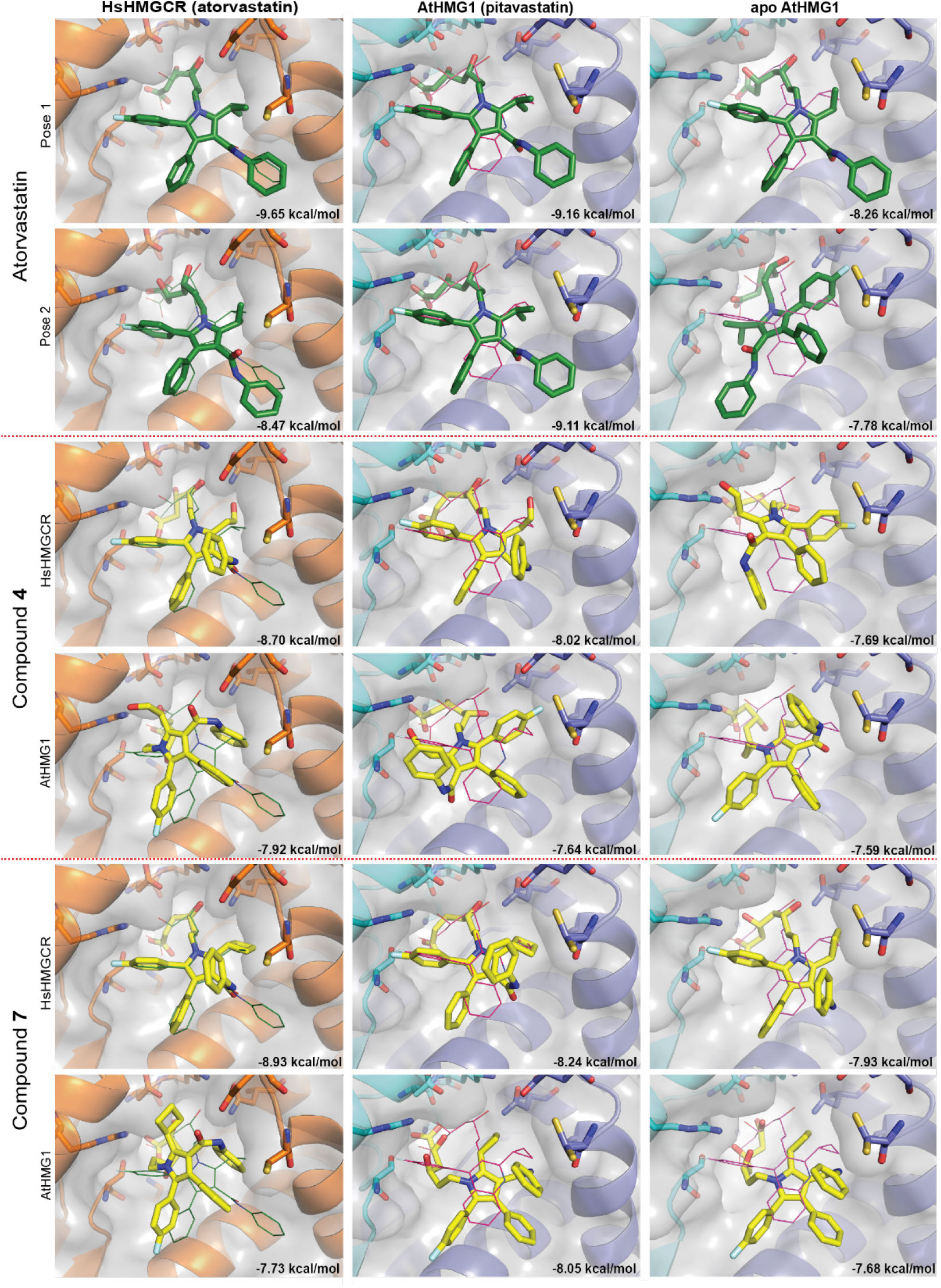
Modelling of atorvastatin and AtHMG1-specific analogs binding to HsHMGCR and AtHMG1. Compounds were docked into HsHMGCR with atorvastatin ligand removed (1HWK, left column, orange cartoon), AtHMG1 with pitavastatin ligand removed (middle column, blue cartoon) and apo AtHMG1 (right column, blue cartoon) using GNINA software^81^. Active site-delineating residues are shown as sticks (orange and blue/cyan). Binding modes of atorvastatin (green line HsHMGCR) and pitavastatin (magenta line AtHMG1) from crystal structures are superimposed for reference. Top two binding poses by affinity are shown. Compounds **4** and **7** (yellow sticks) are predicted to bind HsHMGCR in a manner analogous to atorvastatin (green sticks). Modelling predicts more varied binding modes for AtHMG1 with lower affinity.

## Supplementary Experimental

### General Experimental

All reagents and materials were purchased from commercial suppliers. Thin layer chromatography (TLC) was affected on Merck silica gel 60 F254 aluminium-backed plates and spots stained by heating with vanillin dip (6 g vanillin, 1 mL conc. H_2_SO_4_, 100 mL ethanol), unless stated otherwise. Flash column chromatography was performed on Merck silica gel using the specified solvents. NMR spectra were obtained on a Bruker Avance IIIHD 400, 500 or 600 spectrometers. The solvents used were CDCl_3_ or DMSO-*d*_6_ with CHCl_3_ (^1^H, δ 7.26 ppm), CDCl_3_ (^13^C, δ 77.16 ppm), CD_3_S(O)CD_2_H (^1^H, δ 2.50 ppm) or (CD_3_)_2_SO (^13^C, δ 39.52 ppm) used as an internal standard. Infrared spectra were obtained with neat samples on a PerkinElmer spectrum one FT-IR spectrometer fitted with a PerkinElmer Universal Attenuated Total Reflectance (ATR) sampling accessory. High resolution mass spectra (HR-MS) were obtained on a Waters LCT Premier XE TOF spectrometer, run in W-mode, using the ESI equipped ion source, in positive or negative mode.

### Synthesis of 1-9

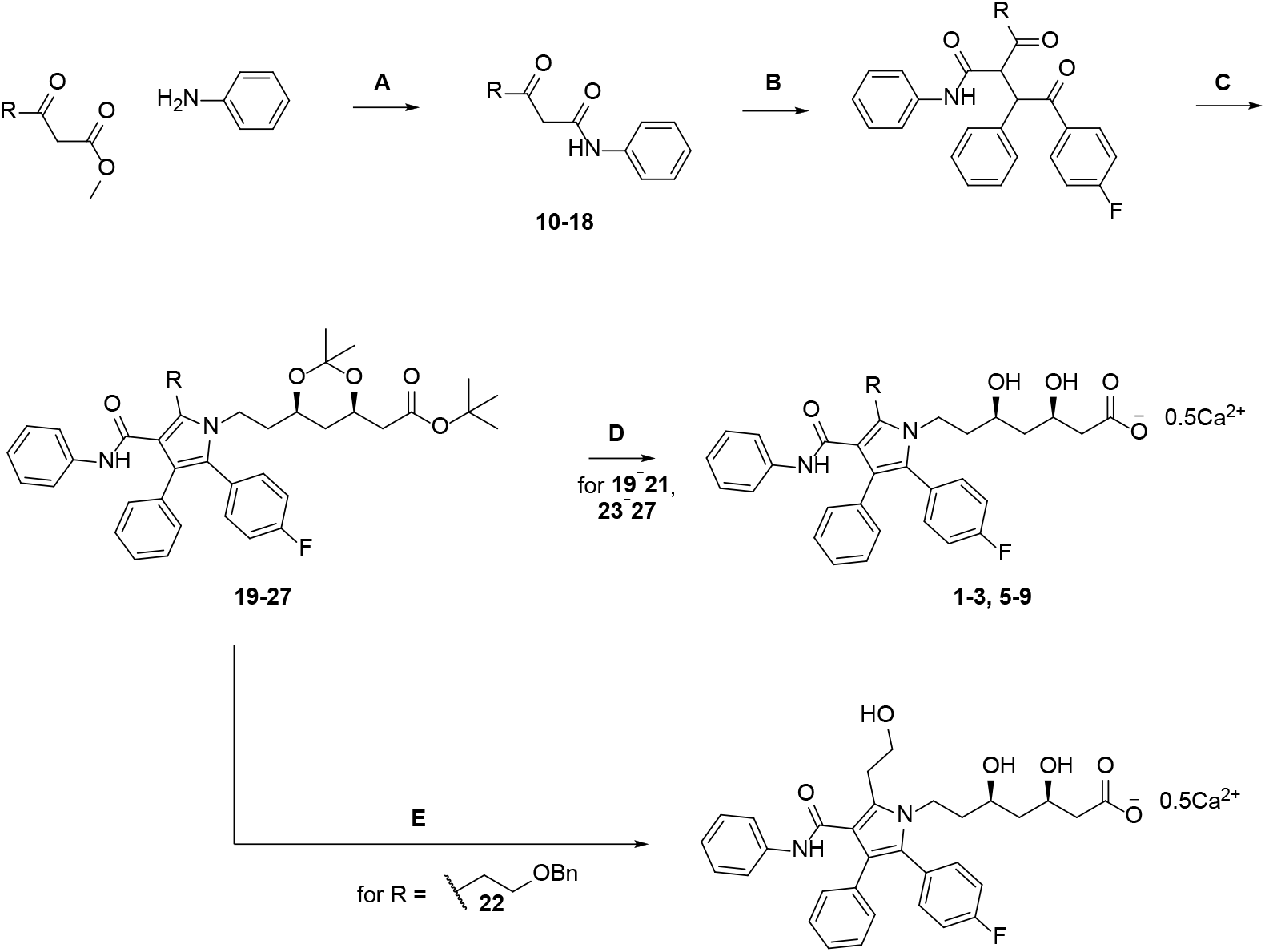

A) TEA, toluene, Δ; B) 2-bromo-1-(3-fluorophenyl)-2-phenylethanone, K_2_CO_3_, acetone; C) (4*R*,6*R*)-*tert*-butyl-6-(2-aminoethyl)-2,2-dimethyl-1,3-dioxane-4-acetate, pivalic acid, 4:1:1 heptane/toluene/THF, Δ; D) i) HCl, MeOH; ii) NaOH, MeOH; iii) Ca(OAc)_2_.H_2_O; E) i) HCl, MeOH; ii) H_2_, Pd(OH)_2_/C, ethanol; iii) NaOH, MeOH; iv) Ca(OAc)_2_.H_2_O.

### General Procedure A^1^

A solution of the methyl 3-oxo-alkanoate (9.50 mmol, 1.0 equiv), aniline (11.4 mmol, 1.2 equiv) and triethylamine (2.37 mmol, 0.25 equiv) in toluene (10 mL) were heated to reflux for 18 h. The solution was allowed to cool to r.t., and the resulting crystalline solid was filtered, washed with toluene (2 x 3 mL) and air dried to yield the compound of interest.

### β-Oxo-*N*-phenylcyclopropanepropanamide (10)

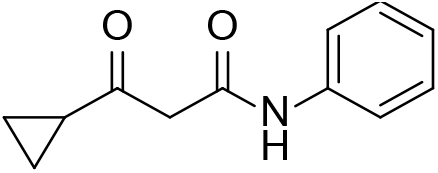

Prepared using General Procedure A (white solid, 1.45 g, 61%). ^1^H NMR (600 MHz, CDCl_3_): δ 9.37 (bs, 1H), 7.55-7.54 (m, 2H), 7.34-7.30 (m, 2H), 7.12-7.09 (m, 1H), 3.72 (s, 2H), 2.07-2.03 (m, 1H), 1.21-1.28 (m, 2H), 1.08-1.05 (m, 2H); ^13^C NMR (151 MHz, CDCl_3_): δ 207.9, 163.8, 137.7, 129.1, 124.6, 120.2, 49.1, 22.1, 12.6; HR-MS (ESI+): *m/z* calculated for C_12_H_14_NO_2_ [M+H]^+^: 204.1025, found: 204.1019.

### 4,4-Dimethyl-3-oxo-N-phenylpentanamide (11)

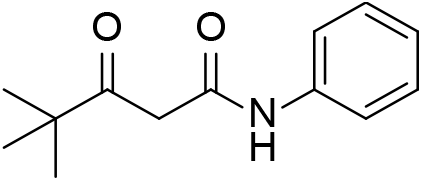

A solution of ethyl 4,4-dimethyl-3-oxo-pentanoate (2.07 mL, 11.6 mmol, 1.0 equiv), aniline (1.27 mL, 13.9 mmol, 1.2 equiv) and triethylamine (0.41 mL, 2.9 mmol, 0.25 equiv) in toluene (10 mL) were heated at 70 °C for 0.5 h, then to reflux for 4 h. The solution was allowed to cool to r.t., washed with 1M HCl (20 mL) and water (2 x 20 mL), dried over MgSO_4_, filtered and concentrated, then purified by silica gel chromatography (10-15% EtOAc/hexanes) to yield the title compound as a pale yellow solid (1.61g, 63%). Spectral data matched those previously reported.^2^

### 3-Oxo-N-phenylhexanamide (12)

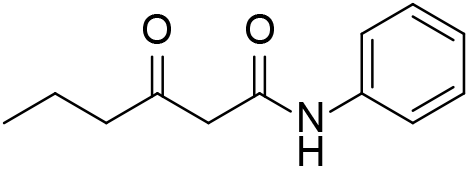

Prepared using General Procedure A (off-white solid, 2.32 g, 82%). ^1^H NMR (400 MHz, CDCl_3_): δ 9.15 (bs, 1H), 7.56-7.53 (m, 2H), 7.35-7.30 (m, 2H), 7.14-7.10 (m, 1H), 3.56 (s, 2H), 2.57 (t, *J* = 7.2 Hz, 2H), 1.67 (tt, *J* = 7.4, 7.2 Hz, 2H), 0.96 (t, *J* = 7.4 Hz, 3H); ^13^C NMR (101 MHz, CDCl_3_): δ 207.7, 163.7, 137.7, 129.1, 124.6, 120.2, 49.2, 46.1, 16.9, 13.6; FTIR (ATR): ʋ 3290, 1711, 1657, 1597, 1547 cm^-1^; HR-MS (ESI+): *m/z* calculated for C_12_H_16_NO_2_ [M+H]^+^: 206.1181, found: 206.1183.

### 3-Oxo-N-phenyl-5-(benzyloxy)pentanamide (13)

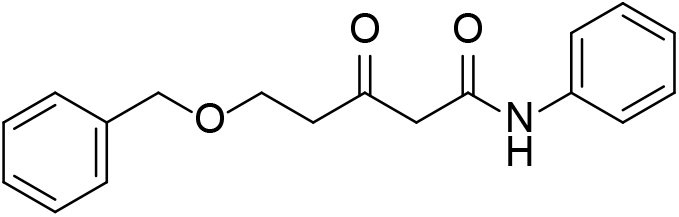

A solution of methyl 3-oxo-5-(benzyloxy)pentanoate^3^ (2.98 g, 12.6 mmol, 1.0 equiv), aniline (2.30 mL, 25.2 mmol, 2.0 equiv) and DMAP (308 mg, 2.52 mmol, 0.20 equiv) in toluene (70 mL) were to reflux for 8 h. The solution was allowed to cool to r.t., then purified by silica gel chromatography (20-100% EtOAc/hexanes) to yield the title compound as a yellow oil (1.23 g, 31%). R_f_ 0.36 (40% EtOAc/hexanes); ^1^H NMR (400 MHz, CDCl_3_): δ 9.05 (bs, 1H), 7.53-7.51 (m, 2H), 7.35-7.27 (m, 7H), 9.13-7.09 (m, 1H), 4.52 (s, 2H), 3.78 (t, *J* = 5.9 Hz, 2H), 3.61 (s, 2H), 2.83 (t, *J* = 5.9 Hz, 2H); ^13^C NMR (126 MHz, CDCl_3_): δ 206.0, 163.6, 137.8, 137.6, 129.1, 128.6, 128.0, 127.9, 124.6, 120.3, 73.5, 64.9, 49.7, 44.2; FTIR (ATR): ʋ 3300, 1716, 1660, 1598, 1543 cm^-1^; HR-MS (ESI+): *m/z* calculated for C_18_H_19_NO_3_Na [M+Na]^+^: 320.1263, found: 320.1263.

### 5-Methyl-3-oxo-N-phenylhexanamide (14)

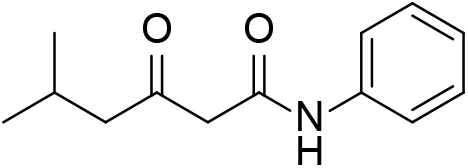

Prepared using General Procedure A (white solid, 843 mg, 61%). The filtrate was concentrated, redissolved in hot toluene (2 mL) and allowed to cool to r.t. The resulting solid was filtered, washed with toluene (2 x 1 mL) and air dried to yield further off-white crystalline solid (168 mg, 12%). Spectral data matched those previously reported.^4^

### 3-Oxo-N-phenylheptanamide (15)

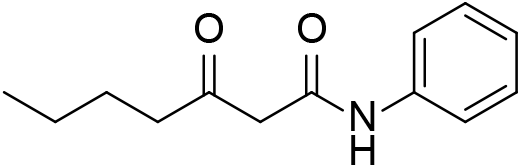

A solution of methyl 3-oxo-heptanoate (1.50 g, 9.48 mmol, 1.0 equiv), aniline (1.04 mL, 11.4 mmol, 1.2 equiv) and triethylamine (0.33 mL, 2.4 mmol, 0.25 equiv) in toluene (10 mL) were heated to reflux for 18 h. The solution was allowed to cool to r.t., washed with 1M HCl (2 x 10 mL) and water (10 mL), dried over MgSO_4_, filtered, concentrated to 5 mL and cooled in ice. The resultant solid was filtered, washed with cold toluene (2 x 3 mL) and air dried to yield the title compound as a cream solid (1.38 g, 66%). ^1^H NMR (400 MHz, CDCl_3_): δ 9.17 (bs, 1H), 7.55-7.53 (m, 2H), 7.34-7.29 (m, 2H), 7.13-7.09 (m, 1H), 3.56 (s, 2H), 2.58 (t, *J* = 7.4 Hz, 2H), 1.60 (tt, *J* = 7.5, 7.4 Hz, 2H), 1.34 (tt, *J* = 7.5, 7.3 Hz, 2H), 0.92 (t, *J* = 7.3 Hz, 3H); ^13^C NMR (101 MHz, CDCl_3_): δ 208.0, 163.6, 137.7, 129.1, 124.6, 120.2, 49.1, 44.0, 25.6, 22.2, 13.9; FTIR (ATR): ʋ 3254, 1713, 1657, 1598, 1548 cm^-1^; HR-MS (ESI+): *m/z* calculated for C_13_H_17_NO_2_Na [M+Na]^+^: 242.1157, found: 242.1153.

### β-Oxo-*N*-phenylcyclobutanepropanamide (16)

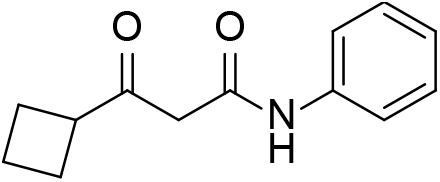

A solution of ethyl 3-cyclobutyl-3-oxopropanoate (1.52 g, 8.93 mmol, 1.0 equiv), aniline (0.98 mL, 11 mmol, 1.2 equiv) and triethylamine (0.31 mL, 2.2 mmol, 0.25 equiv) in xylene (9 mL) were to reflux for 18 h. The solution was allowed to cool to r.t., washed with 1M HCl (10 mL) and water (2 x 10 mL), dried over MgSO_4_, filtered and concentrated, then purified by silica gel chromatography (0-20% EtOAc/hexanes) to yield the title compound as a brown oil (1.32 g, 68%). R_f_ 0.24 (20% EtOAc/hexanes); ^1^H NMR (400 MHz, CDCl_3_): δ 9.27 (bs, 1H), 7.56-7.53 (m, 2H), 7.34-7.29 (m, 2H), 7.13-7.09 (m, 1H), 3.49 (s, 2H), 3.43-3.34 (m, 1H), 2.33-2.17 (m, 4H), 2.07-1.95 (m, 2H), 1.90-1.80 (m, 2H); ^13^C NMR (101 MHz, CDCl_3_): δ 208.5, 163.8, 137.7, 129.1, 124.6, 120.3, 46.5, 46.5, 24.3, 17.6; FTIR (ATR): ʋ 3289, 1699, 1654, 1599, 1532 cm^-1^; HR-MS (ESI+): *m/z* calculated for C_13_H_15_NO_2_Na [M+Na]^+^: 240.1000, found: 240.1000.

### β-Oxo-*N*-phenylcyclopentanepropanamide (17)

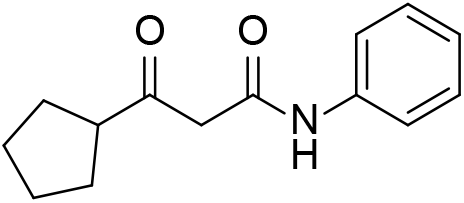

A solution of ethyl 3-cyclopentyl-3-oxopropanoate (1.53 g, 8.31 mmol, 1.0 equiv), aniline (0.91 mL, 10 mmol, 1.2 equiv) and triethylamine (0.29 mL, 2.1 mmol, 0.25 equiv) in xylene (8 mL) were to reflux for 18 h. The solution was allowed to cool to r.t., washed with 1M HCl (10 mL) and water (2 x 10 mL), dried over MgSO_4_, filtered and concentrated, then purified by silica gel chromatography (0-20% EtOAc/hexanes) to yield the title compound as a brown oil (1.40 g, 73%). R_f_ 0.30 (20% EtOAc/hexanes); ^1^H NMR (400 MHz, CDCl_3_): δ 9.27 (bs, 1H), 7.56-7.53 (m, 2H), 7.34-7.29 (m, 2H), 7.13-7.08 (m, 1H), 3.60 (s, 2H), 3.03-2.95 (m, 1H), 1.92-1.74 (m, 4H), 1.72-1.58 (m, 4H); ^13^C NMR (101 MHz, CDCl_3_): δ 210.1, 163.9, 137.7, 129.1, 124.6, 120.2, 52.8, 48.2, 28.7, 26.1; FTIR (ATR): ʋ 3299, 1710, 1658, 1598, 1542 cm^-1^; HR-MS (ESI+): *m/z* calculated for C_14_H_17_NO_2_Na [M+Na]^+^: 254.1157, found: 254.1157.

### β-Oxo-*N*-phenylcyclohexanepropanamide (18)

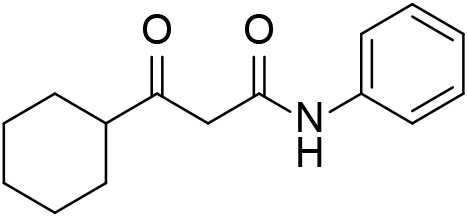

A solution of ethyl 3-cyclohexyl-3-oxopropanoate (1.20 g, 6.05 mmol, 1.0 equiv), aniline (0.66 mL, 7.3 mmol, 1.2 equiv) and triethylamine (0.21 mL, 1.5 mmol, 0.25 equiv) in toluene (6 mL) were heated to reflux for 24 h. The solution was allowed to cool to r.t., washed with 1M HCl (10 mL) and water (2 x 10 mL), dried over MgSO_4_, filtered and concentrated, then purified by silica gel chromatography (0-20% EtOAc/hexanes) to yield the title compound as a cream solid (795 mg, 54%). R_f_ 0.37 (20% EtOAc/hexanes); ^1^H NMR (400 MHz, CDCl_3_): δ 9.22 (bs, 1H), 7.55-7.53 (m, 2H), 7.34-7.29 (m, 2H), 7.13-7.09 (m, 1H), 3.59 (s, 2H), 2.50-2.43 (m, 1H), 1.93-1.89 (m, 2H), 1.83-1.78 (m, 2H), 1.72-1.67 (m, 2H), 1.42-1.16 (m, 4H); ^13^C NMR (101 MHz, CDCl_3_): δ 211.0, 163.8, 137.7, 129.1, 124.6, 120.2, 52.1, 47.3, 28.1, 25.8, 25.5; FTIR (ATR): ʋ 3257, 1711, 1659, 1600, 1557 cm^-1^; HR-MS (ESI+): *m/z* calculated for C_15_H_19_NO_2_Na [M+Na]^+^: 268.1313, found: 268.1312.

### General Procedure B^5^

The 3-oxo-*N*-phenylalkanamide **10**-**18** (5.31 mmol, 1.0 equiv) and 2-bromo-1-(3-fluorophenyl)-2-phenylethanone^6^ (5.31 mmol, 1.0 equiv) and potassium carbonate (7.97 mmol, 1.5 equiv) were stirred in acetone (8 mL) at r.t. while protected from light for 18 h. The mixture was then filtered, and the filtrate was purified by silica gel chromatography (10-20% EtOAc/hexanes) to yield the intermediate 4-fluoro-α-(1-oxoalkyl)-γ-oxo-*N*,β-diphenylbenzene butyramide as a mixture of diastereomers, which were then used in Procedure C.

### General Procedure C^7^

The 4-fluoro-α-(1-oxoalkyl)-γ-oxo-*N*,β-diphenylbenzene butyramide from General Procedure B (2.32 mmol, 1.0 equiv), (4*R*,6*R*)-*tert*-butyl-6-(2-aminoethyl)-2,2-dimethyl-1,3-dioxane-4-acetate (2.39 mmol, 1.03 equiv) and pivalic acid (1.55 mmol, 0.67 equiv) in 4:1:1 heptane/toluene/THF (15 mL) was heated to reflux for 18 h, then cooled to r.t., washed with 0.5 M NaOH (15 mL), 0.5 M HCl (15 mL) and water (5 mL), then dried over MgSO_4_, filtered and concentrated. The residue was purified by silica gel chromatography (5-40% EtOAc/hexanes) to yield the compound of interest.

### 1,1-Dimethylethyl (4R,6*R*)-6-[2-[5-cyclopropyl-2-(4-fluorophenyl)-3-phenyl-4-[(phenylamino)carbonyl]-1*H*-pyrrol-1-yl]ethyl]-2,2-dimethyl-1,3-dioxane-4-acetate (19)

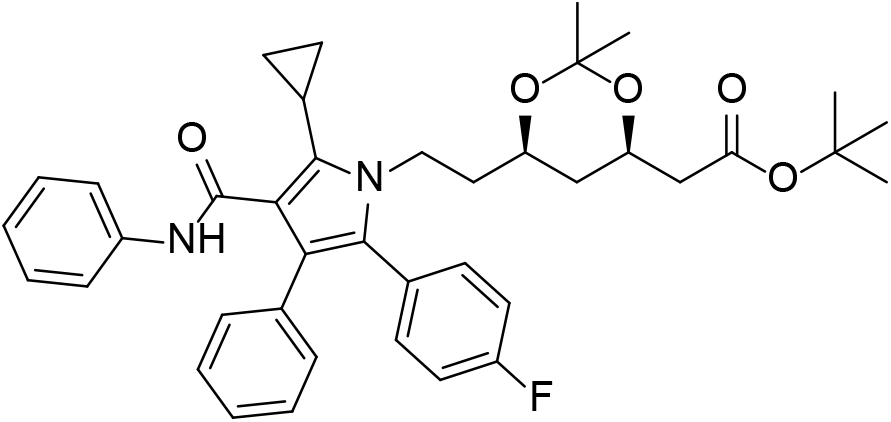

Prepared using General Procedure B (pale yellow resin, 706 mg, 24%; R_f_ 0.19 (20% EtOAc/hexanes)), followed by General Procedure C (pale yellow resin, 800 mg, 75%). R_f_ 0.16 (20% EtOAc/hexanes); ^1^H NMR (500 MHz, CDCl_3_): δ 7.22-7.13 (m, 11H), 7.02-6.97 (m, 3H), 6.89 (s, 1H), 4.25-4.19 (m, 1H), 4.16-4.10 (m, 2H), 3.66-3.61 (m, 1H), 2.36 (dd, *J* = 15.2, 7.0 Hz, 1H), 2.22 (dd, *J* = 15.2, 6.2 Hz, 1H), 1.95-1.89 (m, 1H), 1.66-1.54 (m, 2H), 1.43 (s, 9H), 1.36 (s, 3H), 1.32-1.29 (m, 1H), 1.28 (s, 3H), 1.13-1.09 (m, 2H), 1.02 (ddd, *J* = 11.9, 11.9, 11.9 Hz, 1H), 0.79-0.76 (m, 2H); ^13^C NMR (126 MHz, CDCl_3_): δ 170.3, 163.9, 162.4 (d, *J* = 248 Hz), 138.6, 136.7, 134.5, 133.0 (d, *J* = 8 Hz), 130.6, 129.3, 128.9, 128.4, 128.2 (d, *J* = 3 Hz), 126.7, 123.7, 121.2, 119.6, 118.0, 115.6 (d, *J* = 22 Hz), 98.8, 80.8, 66.3, 66.0, 42.6, 40.5, 37.3, 36.1, 30.1, 28.2, 19.7, 7.8, 7.7, 7.0; FTIR (ATR): ʋ 1727, 1667, 1595, 1509 cm^-1^; HR-MS (ESI+): *m/z* calculated for C_40_H_45_N_2_O_5_FNa [M+Na]^+^: 675.3210, found: 675.3208.

### 1,1-Dimethylethyl (4R,6R)-6-[2-[2-(4-fluorophenyl)-5-(1,1-dimethylethyl)-3-phenyl-4-[(phenylamino)carbonyl]-1*H*-pyrrol-1-yl]ethyl]-2,2-dimethyl-1,3-dioxane-4-acetate (20)

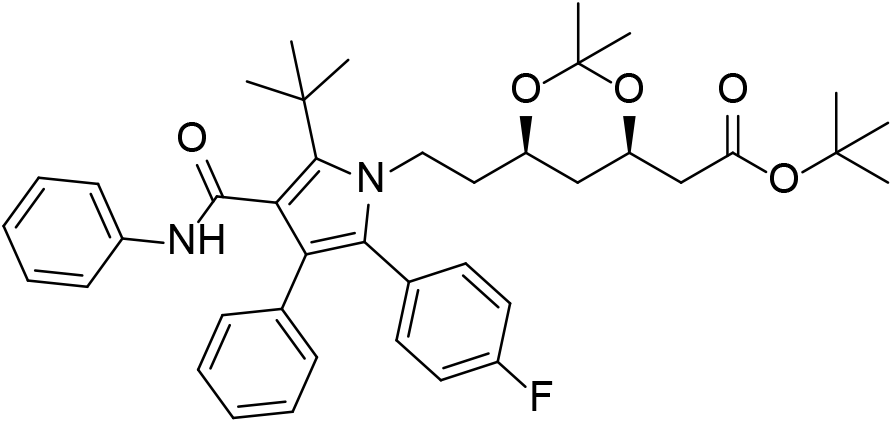

Prepared using General Procedure B (white solid, 594 mg, 19%; R_f_ 0.30 (20% EtOAc/hexanes)), followed by General Procedure C (light yellow resin, 102 mg, 11%). R_f_ 0.28 (20% EtOAc/hexanes); ^1^H NMR (600 MHz, CDCl_3_): δ 7.23-7.21 (m, 2H), 7.18-7.16 (m, 4H), 7.08-7.06 (m, 4H), 7.05-6.99 (m, 5H), 4.38-4.33 (m, 1H), 4.12-4.08 (m, 1H), 4.06-4.01 (m, 1H), 3.55-3.51 (m, 1H), 2.34 (dd, *J* = 15.3, 7.0 Hz, 1H), 2.20 (dd, *J* = 15.3, 6.2 Hz, 1H), 1.58 (s, 9H), 1.53-1.44 (m, 2H), 1.42 (s, 9H), 1.34 (s, 3H), 1.27 (s, 3H), 1.19 (ddd, *J* = 12.7, 2.4, 2.4 Hz, 1H), 0.93 (ddd, *J* = 11.9, 11.9, 11.9 Hz, 1H); ^13^C NMR (151 MHz, CDCl_3_): δ 170.3, 168.0, 162.3 (d, *J* = 248 Hz), 138.1, 134.5, 133.2 (d, *J* = 8 Hz), 130.4, 129.6, 128.9, 128.8 (d, *J* = 3 Hz), 128.1, 125.9, 124.3, 122.0, 120.5, 118.8, 115.6 (d, *J* = 22 Hz), 98.7, 80.8, 66.6, 66.0, 42.6, 37.4, 35.9, 34.2, 31.5, 30.0, 28.2, 19.8; FTIR (ATR): ʋ 3296, 1741, 1650, 1533 cm^-1^; HR-MS (ESI+): *m/z* calculated for C_41_H_49_N_2_O_5_FNa [M+Na]^+^: 691.3523, found: 691.3524.

### 1,1-Dimethylethyl (4R,6R)-6-[2-[2-(4-fluorophenyl)-3-phenyl-4-[(phenylamino)carbonyl-5-propyl]-1*H*-pyrrol-1-yl]ethyl]-2,2-dimethyl-1,3-dioxane-4-acetate (21)

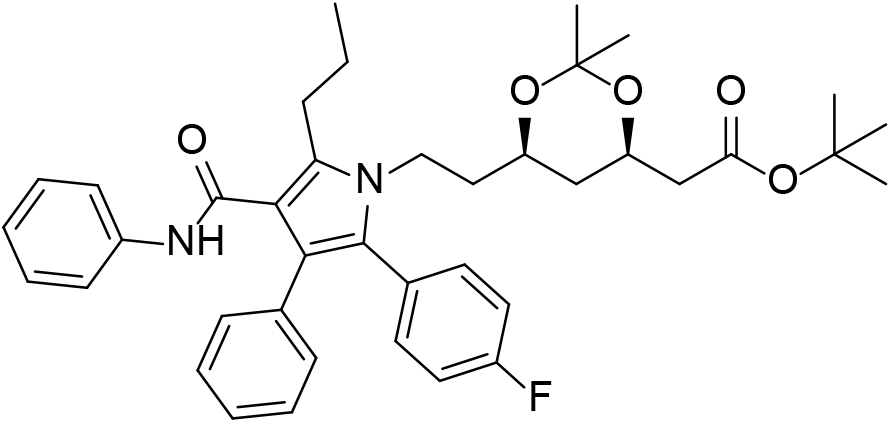

Prepared using General Procedure B, followed by trituration with DCM/hexane (1:5, 4 mL) (white solid, 1.13 g, 56%; R_f_ 0.30 (20% EtOAc/hexanes)), followed by General Procedure C (light yellow resin, 1.21 g, 71%). R_f_ 0.44 (20% EtOAc/hexanes); ^1^H NMR (400 MHz, CDCl_3_): δ 7.28-7.25 (m, 3H), 7.24-7.21 (m, 2H), 7.19-7.13 (m, 4H), 7.06-7.03 (m, 2H), 7.00-6.93 (m, 4H), 4.19-4.13 (m, 1H), 4.06-3.98 (m, 1H), 3.90-3.83 (m, 1H), 3.68-3.61 (m, 1H), 3.20 (dt, *J* = 15.8 7.1 Hz, 1H), 3.04 (dt, *J* = 15.8 7.1 Hz, 1H), 2.38 (dd, *J* = 15.3, 7.0 Hz, 1H), 2.23 (dd, *J* = 15.3, 6.2 Hz, 1H), 1.79-1.70 (m, 2H), 1.65-1.54 (m, 2H), 1.43 (s, 9H), 1.38 (s, 3H), 1.36-1.32 (m, 1H), 1.32 (s, 3H), 1.08-1.00 (m, 1H), 1.07 (t, *J* = 7.3 Hz, 3H); ^13^C NMR (101 MHz, CDCl_3_): δ 170.3, 163.9, 162.4 (d, *J* = 248 Hz), 139.7, 138.7, 134.9, 133.0 (d, *J* = 8 Hz), 131.3, 129.4, 128.8, 128.7, 128.2 (d, *J* = 3 Hz), 127.3, 123.3, 121.6, 119.4, 115.5 (d, *J* = 21 Hz), 114.1, 98.9, 80.8, 66.0, 42.6, 40.2, 37.8, 36.2, 30.1, 28.2, 27.5, 23.9, 19.8, 14.5; FTIR (ATR): ʋ 3401, 1727, 1661, 1595, 1529 cm^-1^; HR-MS (ESI+): *m/z* calculated for C_40_H_47_N_2_O_5_FNa [M+Na]^+^: 677.3367, found: 677.3369.

### 1,1-Dimethylethyl (4R,6R)-6-[5-(2-benzyloxyethyl)-2-[2-(4-fluorophenyl)-3-phenyl-4-[(phenylamino)carbonyl]-1*H*-pyrrol-1-yl]ethyl]-2,2-dimethyl-1,3-dioxane-4-acetate (22)

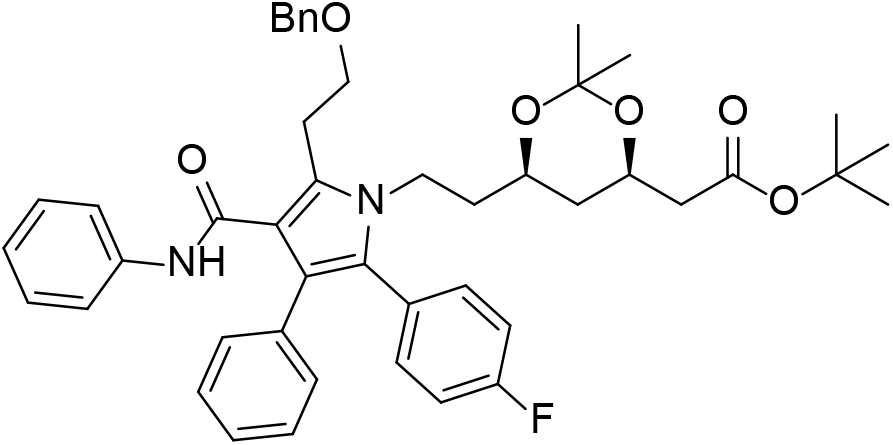

Prepared using General Procedure B with 2-iodo-1-(3-fluorophenyl)-2-phenylethanone^8^ (yellow resin, 936 mg, 46%; R_f_ 0.22 (20% EtOAc/hexanes)), followed by General Procedure C (light yellow resin, 204 mg, 15%). *Note* - unstable in CHCl_3_. ^1^H NMR (500 MHz, CD_3_CN): δ 8.18 (s, 1H), 7.33-7.18 (m, 16H), 2.09-2.05 (m, 2H), 7.00-6.97 (m, 1H), 4.62-4.57 (m, 2H), 4.16-4.11 (m, 1H), 3.99-3.93 (m, 1H), 3.89-3.82 (m, 3H), 3.67-3.62 (m, 1H), 3.38-3.32 (m, 1H), 3.30-3.24 (m, 1H), 2.23 (dd, *J* = 15.0, 4.9 Hz, 1H), 2.18-2.13 (m, 1H), 1.53-1.40 (m, 11H), 1.32 (s, 3H), 1.27-1.23 (m, 1H), 1.20 (m, 3H), 0.92-0.85 (ddd, *J* = 11.9, 11.9, 11.9 Hz, 1H); ^13^C NMR (126 MHz, CD_3_CN): δ 170.9, 164.7, 163.3 (d, *J* = 246 Hz), 140.1, 139.5, 136.3, 134.5 (d, *J* = 8 Hz), 134.0, 131.7, 131.0, 129.6, 129.5 (d, *J* = 3 Hz), 129.4, 129.0, 128.7, 128.6, 127.5, 124.1, 123.2, 120.1, 117.9, 116.1 (d. *J* = 22 Hz), 99.4, 81.0, 73.6, 70.7, 67.0, 43.4, 41.2, 38.1, 36.6, 30.4, 28.3, 26.8, 20.0. FTIR (ATR): ʋ 3401, 1727, 1660, 1530 cm^-1^; HR-MS (ESI+): *m/z* calculated for C_46_H_52_N_2_O_6_F [M+H]^+^: 747.3809, found: 747.3804.

### 1,1-Dimethylethyl (4R,6*R*)-6-[2-[2-(4-fluorophenyl)-5-(2-methylpropyl)-3-phenyl-4-[(phenylamino)carbonyl]-1*H*-pyrrol-1-yl]ethyl]-2,2-dimethyl-1,3-dioxane-4-acetate (23)

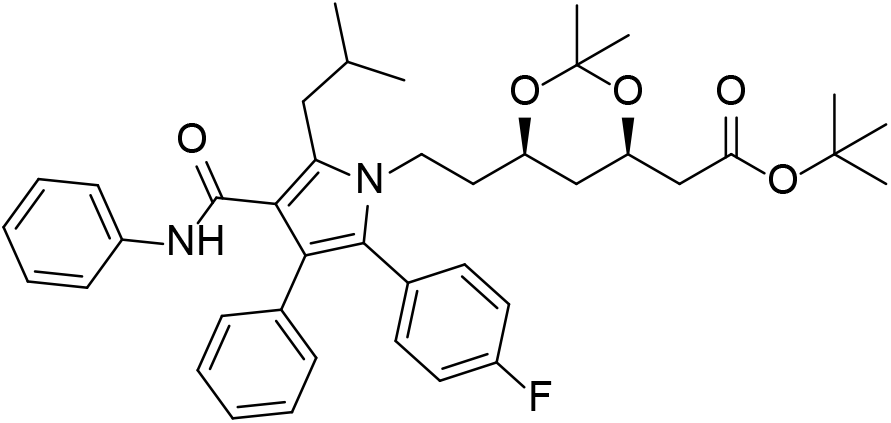

Prepared using General Procedure B (off-white solid, 977 mg, 49%; R_f_ 0.34 (20% EtOAc/hexanes)), followed by General Procedure C (off-white resin, 793 mg, 53%). R_f_ 0.40 (20% EtOAc/hexanes); ^1^H NMR (500 MHz, CDCl_3_): δ 7.29-7.23 (m, 3H), 7.22-7.20 (m, 2H), 7.18-7.13 (m, 4H), 7.04-7.02 (m, 2H), 7.00-6.94 (m, 3H), 6.91 (s, 1H), 4.14-4.12 (m, 1H), 4.06-4.00 (m, 1H), 3.94-3.88 (m, 1H), 3.62-3.57 (m, 1H), 3.19 (dd, *J* = 14.2, 7.6 Hz, 1H), 3.01 (dd, *J* = 14.2, 7.1 Hz, 1H), 2.37 (dd, *J* = 15.3, 7.0 Hz, 1H), 2.22 (dd, *J* = 15.3, 6.2 Hz, 1H), 2.05-1.97 (m, 1H), 1.56-1.44 (m, 2H), 1.43 (s, 9H), 1.38 (s, 3H), 1.33 (s, 3H), 1.30 (ddd, *J* = 12.8, 2.4, 2.4 Hz, 1H), 1.05-0.98 (m, 7H); ^13^C NMR (126 MHz, CDCl_3_): δ 170.3, 164.2, 162.3 (d, *J* = 248 Hz), 138.6, 138.6, 134.9, 133.0 (d, *J* = 8 Hz), 131.2, 129.5, 128.8, 128.7, 128.3 (d, *J* = 3 Hz), 127.2, 123.3, 121.6, 119.5, 115.5 (d, *J* = 22 Hz), 115.1, 98.9, 80.8, 66.0, 65.8, 42.6, 40.3, 37.4, 36.2, 30.1, 30.1, 28.2, 22.7, 22.5, 19.8; FTIR (ATR): ʋ 3405, 1727, 1663, 1595, 1528 cm^-1^; HR-MS (ESI+): *m/z* calculated for C_41_H_49_N_2_O_5_FNa [M+Na]^+^: 691.3523, found: 691.3528.

### 1,1-Dimethylethyl (4R,6R)-6-[2-[5-butyl-2-(4-fluorophenyl)-3-phenyl-4-[(phenylamino)carbonyl]-1*H*-pyrrol-1-yl]ethyl]-2,2-dimethyl-1,3-dioxane-4-acetate (24)

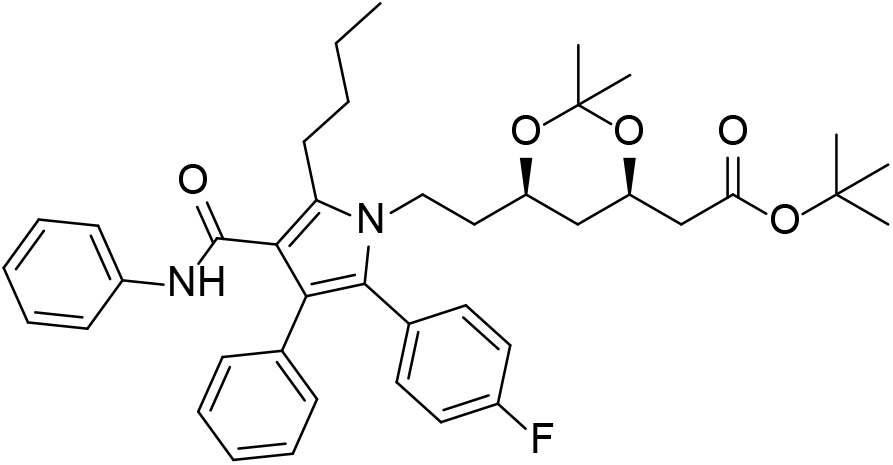

Prepared using General Procedure B (off-white resin, 1.29 g, 56%; R_f_ 0.35 (20% EtOAc/hexanes)), followed by General Procedure C (light yellow resin, 914 mg, 59%). R_f_ 0.50 (20% EtOAc/hexanes); ^1^H NMR (500 MHz, CDCl_3_): δ 7.30-7.25 (m, 3H), 7.24-7.21 (m, 2H), 7.18-7.13 (m, 4H), 7.06-7.04 (m, 2H), 7.00-6.94 (m, 4H), 4.19-4.13 (m, 1H), 4.02 (ddd, *J* = 14.4, 9.7, 4.7 Hz, 1H), 3.87 (ddd, *J* = 14.4, 9.4, 7.0 Hz, 1H), 3.67-3.62 (m, 1H), 3.23 (ddd, *J* = 14.1, 9.0, 7.0 Hz, 1H), 3.07 (ddd, *J* = 14.1, 8.9, 6.9 Hz, 1H), 2.38 (dd, *J* = 15.3, 7.0 Hz, 1H), 2.23 (dd, *J* = 15.3, 6.2 Hz, 1H), 1.72-1.46 (m, 6H), 1.43 (s, 9H), 1.38 (s, 3H), 1.35-1.32 (m, 1H), 1.32 (s, 3H), 1.05 (ddd, *J* = 11.9, 11.9, 11.9 Hz, 1H), 0.99 (t, *J* = 7.4 Hz, 3H); ^13^C NMR (126 MHz, CDCl_3_): δ 170.3, 163.9, 162.3 (d, *J* = 248 Hz), 139.9, 138.7, 134.9, 133.0 (d, *J* = 8 Hz), 131.3, 129.4, 128.8, 128.7, 128.2 (d, *J* = 3 Hz), 127.3, 123.3, 121.6, 119.4, 115.5 (d, *J* = 21 Hz), 114.0, 98.9, 80.8, 66.0, 42.6, 40.2, 37.7, 36.2, 32.8, 30.1, 28.2, 25.2, 23.1, 19.8, 14.2; FTIR (ATR): ʋ 3404, 1727, 1662, 1594, 1529 cm^-1^; HR-MS (ESI+): *m/z* calculated for C_41_H_49_N_2_O_5_FNa [M+Na]^+^: 691.3523, found: 691.3521.

### 1,1-Dimethylethyl (4R,6R)-6-[2-[5-cyclobutyl-2-(4-fluorophenyl)-3-phenyl-4-[(phenylamino)carbonyl]-1*H*-pyrrol-1-yl]ethyl]-2,2-dimethyl-1,3-dioxane-4-acetate (25)

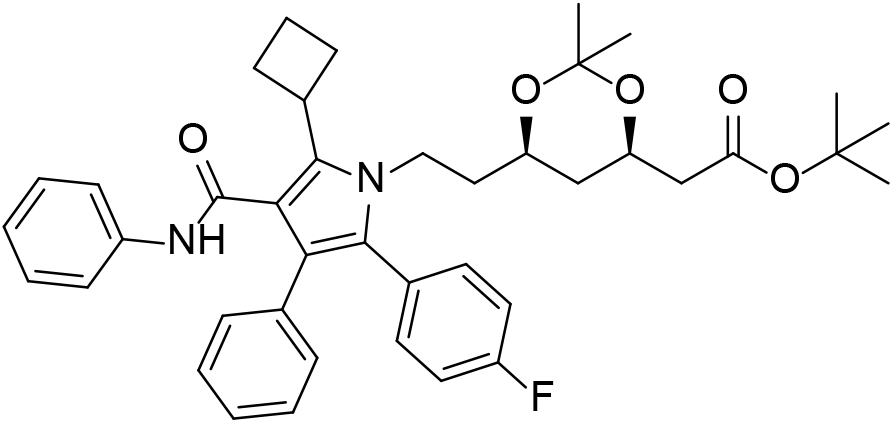

Prepared using General Procedure B (cream solid, 1.08 g, 55%; R_f_ 0.28 (20% EtOAc/hexanes)), followed by General Procedure C (light yellow resin, 1.30 g, 88%). R_f_ 0.32 (20% EtOAc/hexanes); ^1^H NMR (600 MHz, CDCl_3_): δ 7.22-7.11 (m, 11H), 7.02-6.98 (m, 3H), 6.88 (bs, 1H), 4.16-4.11 (m, 1H), 4.04 (ddd, *J* = 14.5, 9.9, 4.8 Hz, 1H), 3.97-3.91 (m, 1H), 3.85 (ddd, *J* = 14.5, 9.8, 6.3 Hz, 1H), 3.64-3.60 (m, 1H), 2.64-2.55 (m, 2H), 2.47-2.42 (m, 2H), 2.37 (dd, *J* = 15.3, 7.0 Hz, 1H), 2.22 (dd, *J* = 15.3, 6.2 Hz, 1H), 2.06-1.98 (m, 1H), 1.92-1.87 (m, 1H), 1.61-1.58 (m, 2H), 1.43 (s, 9H), 1.36 (s, 3H), 1.32-1.29 (m, 1H), 1.31 (s, 3H), 1.02 (ddd, *J* = 11.9, 11.9, 11.9 Hz, 1H); ^13^C NMR (151 MHz, CDCl_3_): δ 170.3, 165.1, 162.4 (d, *J* = 248 Hz), 138.4, 138.1, 134.6, 133.1 (d, *J* = 8 Hz), 130.3, 129.1, 128.9, 128.4, 128.3 (d, *J* = 3 Hz), 126.5, 123.8, 121.6, 119.9, 117.1, 115.6 (d, *J* = 21 Hz), 98.8, 80.8, 66.3, 66.0, 42.6, 40.8, 37.7, 36.1, 33.0, 30.1, 29.6, 29.3, 28.2, 19.8, 18.9; FTIR (ATR): ʋ 3409, 1725, 1667, 1595, 1526, 1509 cm^-1^; HR-MS (ESI+): *m/z* calculated for C_41_H_47_N_2_O_5_FNa [M+Na]^+^: 689.3367, found: 689.3372.

### 1,1-Dimethylethyl (4R,6*R*)-6-[2-[5-cyclopentyl-2-(4-fluorophenyl)-3-phenyl-4-[(phenylamino)carbonyl]-1*H*-pyrrol-1-yl]ethyl]-2,2-dimethyl-1,3-dioxane-4-acetate (26)

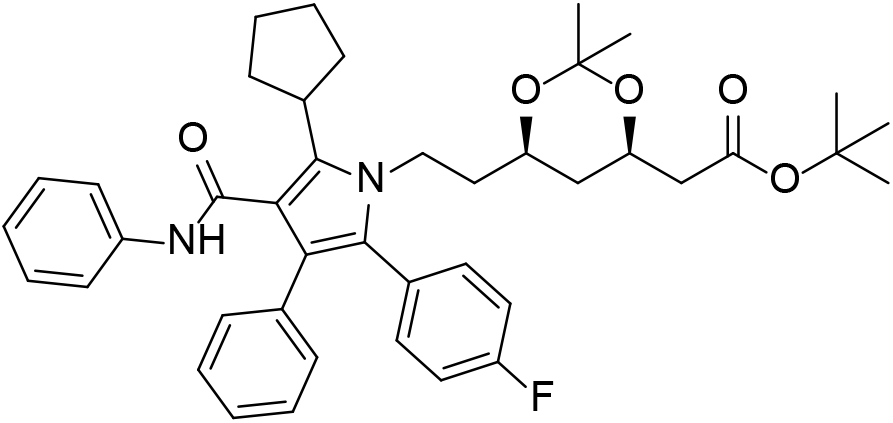

Prepared using General Procedure B (white solid, 1.20 g, 62%; R_f_ 0.32 (20% EtOAc/hexanes)), followed by General Procedure C (pale yellow resin, 910 mg, 58%). R_f_ 0.44 (20% EtOAc/hexanes); ^1^H NMR (600 MHz, CDCl_3_): δ 7.21-7.15 (m, 9H), 7.06-7.04 (m, 2H), 7.00-6.96 (m, 3H), 6.85 (bs, 1H), 4.17-4.13 (m, 1H), 4.11-4.06 (m, 1H), 3.83 (ddd, *J* = 14.5, 10.5, 5.9 Hz, 1H), 3.69-3.64 (m, 1H), 3.64-3.58 (m, 1H), 2.38 (dd, *J* = 15.3, 7.0 Hz, 1H), 2.23 (dd, *J* = 15.3, 6.2 Hz, 1H), 2.16-2.11 (m, 2H), 2.09-2.01 (m, 2H), 1.99-1.92 (m, 2H), 1.73-1.58 (m, 4H), 1.43 (s, 9H), 1.36 (s, 3H), 1.33 (ddd, *J* = 12.7, 2.4, 2.4 Hz, 1H), 1.30 (s, 3H), 1.04 (ddd, *J* = 11.9, 11.9, 11.9 Hz, 1H); ^13^C NMR (151 MHz, CDCl_3_): δ 170.3, 164.8, 162.4 (d, *J* = 248 Hz), 139.6, 138.6, 134.8, 133.2 (d, *J* = 8 Hz), 130.7, 129.3, 128.8, 128.5, 128.4 (d, *J* = 3 Hz), 126.7, 123.6, 122.0, 119.6, 115.8, 115.5 (d, *J* = 21 Hz), 98.8, 80.8, 66.5, 66.0, 42.6, 41.0, 38.1, 37.1, 36.1, 32.7, 32.5, 30.0, 28.2, 26.6, 26.5, 19.8; FTIR (ATR): ʋ 3409, 1726, 1666, 1595, 1525, 1509 cm^-1^; HR-MS (ESI+): *m/z* calculated for C_42_H_49_N_2_O_5_FNa [M+Na]^+^: 703.3523, found: 703.3519.

### 1,1-Dimethylethyl (4*R*,6*R*)-6-[2-[5-cyclohexyl-2-(4-fluorophenyl)-3-phenyl-4-[(phenylamino)carbonyl]-1*H*-pyrrol-1-yl]ethyl]-2,2-dimethyl-1,3-dioxane-4-acetate (27)

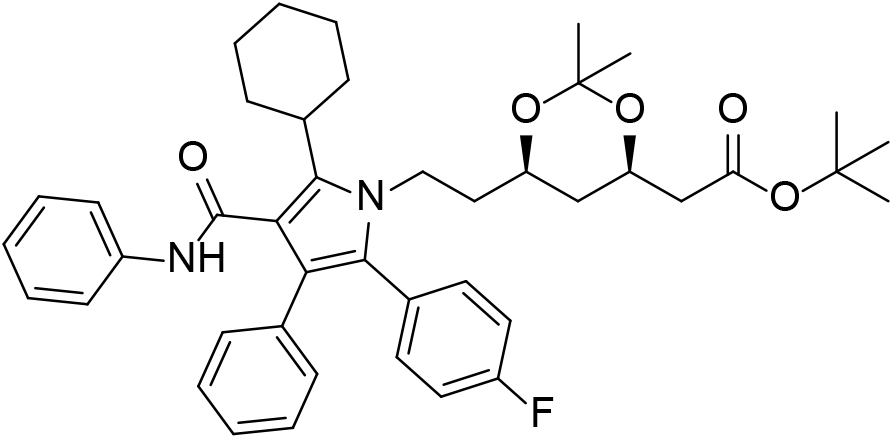

Prepared using General Procedure B (cream solid, 600 mg, 43%; R_f_ 0.36 (20% EtOAc/hexanes)), followed by General Procedure C (off-white resin, 295 mg, 33%). R_f_ 0.40 (20% EtOAc/hexanes); ^1^H NMR (400 MHz, CDCl_3_): δ 7.21-7.13 (m, 9H), 7.10-7.08 (m, 2H), 7.01-6.97 (m, 3H), 6.85 (bs, 1H), 4.20-4.13 (m, 1H), 4.12-4.05 (m, 1H), 3.84-3.76 (m, 1H), 3.72-3.66 (m, 1H), 3.11-3.05 (m, 1H), 2.39 (dd, *J* = 15.3, 6.9 Hz, 1H), 2.25 (dd, *J* = 15.3, 6.2 Hz, 1H), 2.24-2.13 (m, 2H), 1.88-1.85 (m, 4H), 1.68-1.59 (m, 2H), 1.44 (s, 9H), 1.40-1.34 (m, 6H), 1.33 (s, 3H), 1.08 (ddd, *J* = 11.9, 11.9, 11.9 Hz, 1H); ^13^C NMR (101 MHz, CDCl_3_): δ 170.3, 165.2, 162.4 (d, *J* = 248 Hz), 140.3, 138.6, 134.8, 133.3 (d, *J* = 8 Hz), 130.5, 128.8, 128.8, 128.5 (d, *J* = 3 Hz), 128.4, 126.5, 123.7, 121.9, 119.8, 116.0, 115.5 (d, *J* = 21 Hz), 98.8, 80.8, 66.5, 66.1, 42.6, 40.9, 38.4, 37.2, 36.2, 31.7, 31.6, 30.2, 28.2, 27.5, 25.9, 19.8; FTIR (ATR): ʋ 3409, 1727, 1667, 1595, 1525, 1509 cm^-1^; HR-MS (ESI+): *m/z* calculated for C_43_H_51_N_2_O_5_FNa [M+Na]^+^: 717.3680, found: 717.3683.

### General Procedure D^9^

To a stirred solution of the 1,1-dimethylethyl (4*R*,6*R*)-6-[2-[5-(alkyl)-2-(4-fluorophenyl)-3-phenyl-4-[(phenylamino)carbonyl]-1*H*-pyrrol-1-yl]ethyl]-2,2-dimethyl-1,3-dioxane-4-acetate **19**-**21**, **23**-**27** (0.75 mmol, 1.0 equiv) in methanol (12 mL) was added 1M HCl (1.1 mL, 1.1 mmol, 1.5 equiv). After 1.5 h at r.t., the solution was purified by silica gel chromatography (20-50% EtOAc/hexanes), then dissolved in methanol (5 mL) and 1M NaOH (1.5 mL, 1.5 mmol, 2.0 equiv) was added. The solution was stirred at r.t. for 2 h, then a solution of calcium acetate hydrate (72 mg, 0.41 mmol, 0.55 equiv) in water (1 mL) was added dropwise. After stirring for an additional 15 min, the precipitate was filtered, washed with water (3 x 5 mL) and dried to yield the compound of interest.

### (β*R*,δ*R*)-5-Cyclopropyl-2-(4-fluorophenyl)-β,δ-dihydroxy-3-phenyl-4-[(phenylamino)carbonyl]-1*H*-pyrrole-1-heptanoic acid hemicalcium salt (1)

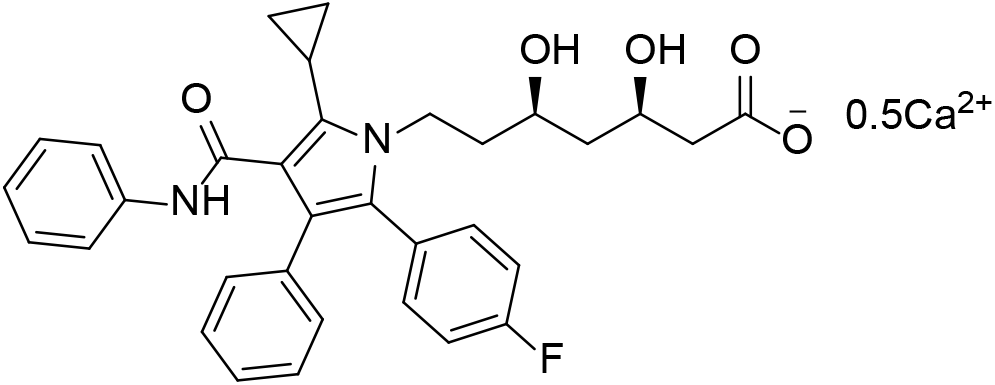

Prepared using General Procedure D (white powder, 272 mg, 41%). ^1^H NMR (600 MHz, DMSO-d_6_): δ 10.02 (s, 1H), 7.62-7.61 (m, 2H), 7.27-7.22 (m, 4H), 7.20-7.17 (m, 2H), 7.10-7.07 (m, 2H), 7.07-6.99 (m, 4H), 5.95 (bs, 1H), 4.68 (bs, 1H), 4.13-4.08 (m, 1H), 4.00-3.95 (m, 1H), 3.79-3.76 (m, 1H), 3.57-3.54 (m, 1H), 2.07 (dd, *J* = 15.3, 4.1 Hz, 1H), 1.97-1.92 (m, 2H), 1.66-1.60 (m, 1H), 1.57-1.51 (m, 1H), 1.42-1.37 (m, 1H), 1.26-1.22 (m, 1H), 0.87-0.85 (m, 2H), 0.67-0.65 (m, 2H); ^13^C NMR (151 MHz, DMSO-d_6_): δ 178.0, 165.1, 161.6 (d, *J* = 245 Hz), 139.6, 134.9, 133.2 (d, *J* = 8 Hz), 132.0, 129.3, 128.5, 128.3, 127.6, 125.3, 122.9, 120.2, 119.3, 119.0, 115.4 (d, *J* = 21 Hz), 66.3, 66.3, 44.0, 43.8, 41.1, 38.1, 6.6, 5.9, 5.9; FTIR (ATR): ʋ 3399, 1650, 1594, 1558, 1509 cm^-1^; HR-MS (ESI-): *m/z* calculated for C_33_H_32_N_2_O_5_F [M-0.5Ca]^-^: 555.2295, found: 555.2299.

### (β*R*,δ*R*)-2-(4-Fluorophenyl)-β,δ-dihydroxy-5-(1,1-dimethylethyl)-3-phenyl-4-[(phenylamino)carbonyl]-1*H*-pyrrole-1-heptanoic acid hemicalcium salt (2)

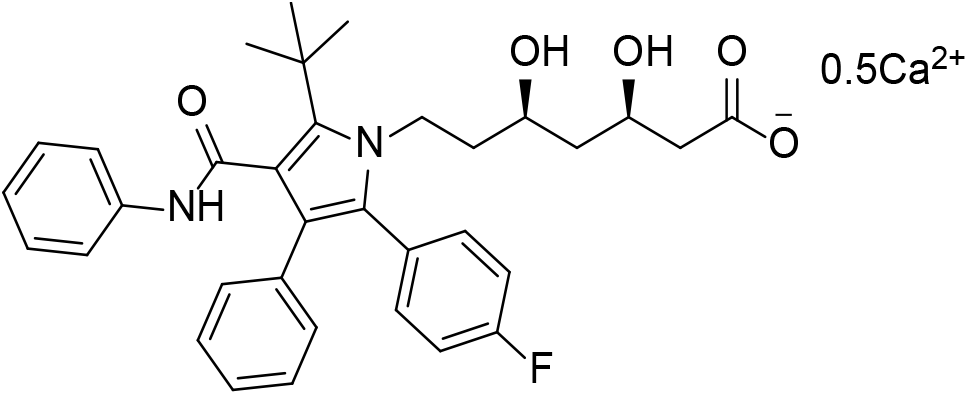

Prepared using General Procedure D (white powder, 27 mg, 34%). ^1^H NMR (600 MHz, DMSO-d_6_): δ 10.16 (s, 1H), 7.52-7.50 (m, 2H), 7.22-7.15 (m, 6H), 7.07-7.03 (m, 4H), 6.99-6.97 (m, 2H), 6.13 (bs, 1H), 4.72 (bs, 1H), 4.29-4.25 (m, 1H), 4.07-4.03 (m, 1H), 3.60-3.57 (m, 1H), 3.42-3.38 (m, 1H), 1.99 (dd, *J* = 14.9, 2.9 Hz, 1H), 1.85 (dd, *J* = 14.9, 8.0 Hz, 1H), 1.50 (s, 9H), 1.41-1.32 (m, 2H), 1.30-1.23 (m, 1H), 1.05-1.03 (m, 1H); ^13^C NMR (151 MHz, DMSO-d_6_): δ 177.7, 167.5, 161.5 (d, *J* = 245 Hz), 139.4, 135.2, 134.9, 133.2 (d, *J* = 8 Hz), 129.6, 129.2, 128.9 (d, *J* = 3 Hz), 128.4, 127.5, 125.4, 123.0, 120.8, 119.5, 118.9, 115.4 (d, *J* = 21 Hz), 66.4, 66.2, 43.8, 42.8, 38.2, 33.4, 30.8; FTIR (ATR): ʋ 3317, 1661, 1594, 1563, 1508 cm^-1^; HR-MS (ESI-): *m/z* calculated for C_34_H_36_N_2_O_5_F [M-0.5Ca]^-^: 571.2613, found: 571.2611.

### (β*R*,δ*R*)-2-(4-Fluorophenyl)-β,δ-dihydroxy-3-phenyl-4-[(phenylamino)carbonyl-5-propyl]-1*H*-pyrrole-1-heptanoic acid hemicalcium salt (3)

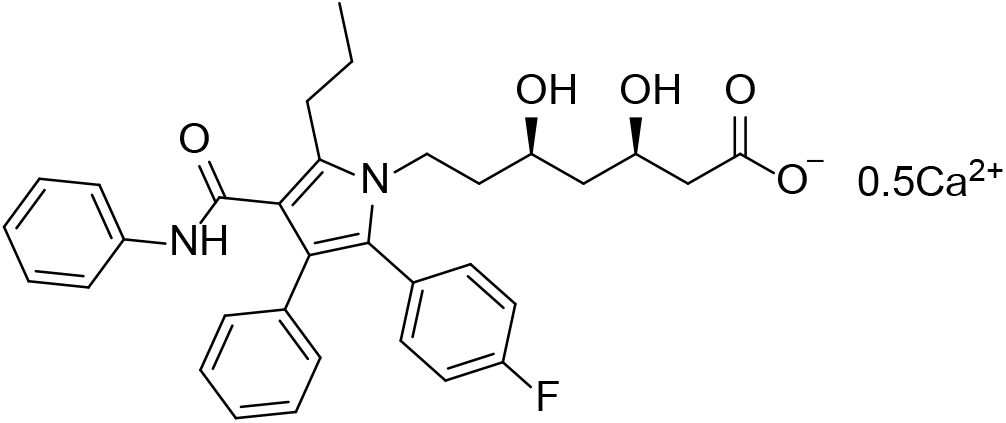

Prepared using General Procedure D (white powder, 107 mg, 24%). ^1^H NMR (600 MHz, DMSO-d_6_): δ 9.28 (s, 1H), 7.43-7.42 (m, 2H), 7.26-7.04 (m, 11H), 6.98-6.96 (m, 1H), 6.03 (bs, 1H), 4.77 (bs, 1H), 3.97-3.93 (m, 1H), 3.81-3.73 (m, 2H), 3.54-3.50 (m, 1H), 2.85-2.82 (m, 2H), 2.08-2.06 (m, 1H), 1.95-1.91 (m, 1H), 1.64-1.53 (m, 3H), 1.49-1.43 (m, 1H), 1.41-1.36 (m, 1H), 1.23-1.19 (m, 1H), 0.93 (t, *J* = 6.7 Hz, 3H); ^13^C NMR (151 MHz, DMSO-d_6_): δ 178.0, 164.8, 161.6 (d, *J* = 245 Hz), 139.5, 134.8, 133.9, 133.3 (d, *J* = 8 Hz), 129.6, 128.5, 128.3, 127.7, 125.6, 122.8, 120.8, 119.2, 117.1, 115.4 (d, *J* = 21 Hz), 66.3, 66.1, 43.9, 43.7, 40.8, 38.6, 26.6, 23.4, 14.2; FTIR (ATR): ʋ 3396, 1660, 1594, 1558, 1530 cm^-1^; HR-MS (ESI-): *m/z* calculated for C_33_H_34_N_2_O_5_F [M-0.5Ca]^-^: 557.2416, found: 557.2433.

### (β*R*,δ*R*)-2-(4-Fluorophenyl)-β,δ-dihydroxy-5-(2-hydroxyethyl)-3-phenyl-4-[(phenylamino)carbonyl]-1*H*-pyrrole-1-heptanoic acid hemicalcium salt (4)

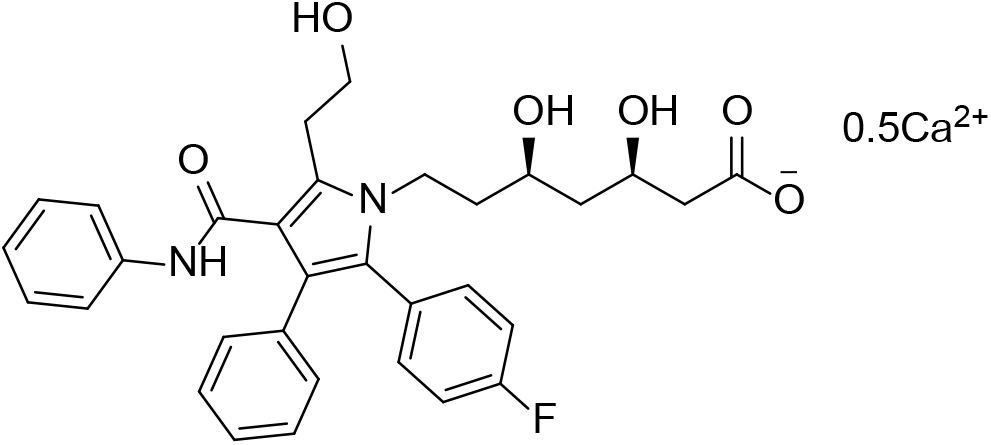

To a solution of 1,1-dimethylethyl (4*R*,6*R*)-6-[5-(2-benzyloxyethyl)-2-[2-(4-fluorophenyl)-3-phenyl-4-[(phenylamino)carbonyl]-1*H*-pyrrol-1-yl]ethyl]-2,2-dimethyl-1,3-dioxane-4-acetate (**22**) (100 mg, 0.134 mmol) in methanol (2 mL) was added 1M HCl (0.30 mL) and the solution was stirred at r.t. for 2 h. The solution was diluted with EtOAc (10 mL) and washed with water (2 x 5 mL), dried over MgSO_4_, filtered and concentrated, then purified by silica gel chromatography (20-100% EtOAc/hexanes) to yield 1,1-dimethylethyl (3*R*,5*R*)-7-[5-(2-benzyloxyethyl)-2-(4-fluorophenyl)-3-phenyl-4-phenylcarbamoylpyrrol-1-yl]-3,5-dihydroxyheptanoate as a colourless resin (68 mg, 72%). R_f_ 0.17 (40% EtOAc/hexanes); ^1^H NMR (400 MHz, CDCl_3_): δ 7.40 (s, 1H), 7.33-7.29 (m, 4H), 7.28-7.22 (m, 6H), 7.19-7.14 (m, 4H), 7.10-7.08 (m, 2H), 7.01-6.95 (m, 3H), 4.58 (s, 2H), 4.14-3.97 (m, 3H), 3.93 (t, *J* = 6.1 Hz, 2H), 3.69-3.63 (m, 1H), 3.46 (t, *J* = 6.1 Hz, 2H), 3.11 (bs, 2H), 2.30-2.27 (m, 2H), 1.65-1.51 (m, 2H), 1.46 (s, 9H), 1.38 (ddd, *J* = 14.2, 10.1, 10.1 Hz, 1H), 1.18 (ddd, *J* = 14.2, 2.2, 2.2 Hz, 1H); ^13^C NMR (101 MHz, CDCl_3_): δ 172.2, 163.8, 162.4 (d, *J* = 248 Hz), 138.7, 138.5, 135.7, 134.9, 133.1 (d, *J* = 8 Hz), 131.2, 130.1, 128.8, 128.6, 128.5, 128.2 (d, *J* = 3 Hz), 127.8, 127.7, 127.2, 123.3, 122.0, 119.4, 115.5 (d, *J* = 21 Hz), 115.3, 81.8, 73.1, 70.7, 69.4, 69.1, 42.4, 41.9, 40.9, 38.8, 28.2, 26.5.

To 1,1-dimethylethyl (3*R*,5*R*)-7-[5-(2-benzyloxyethyl)-2-(4-fluorophenyl)-3-phenyl-4-phenylcarbamoylpyrrol-1-yl]-3,5-dihydroxyheptanoate (68 mg, 0.096 mmol) and 20% Pd(OH)_2_/C (8.4 mg) was added ethanol (2 mL), then the atmosphere was evacuated and filled with H_2_ (x 3) and stirred vigorously under a balloon H_2_ for 5 h. The mixture was filtered through celite and purified by silica gel chromatography (50-100% EtOAc/hexanes; R_f_ 0.48 (EtOAc)) to yield a colourless resin, which was dissolved in methanol (0.3 mL) and 1M NaOH (0.1 mL) was added. The solution was stirred at r.t. for 1 h, then a solution of calcium acetate hydrate (8 mg, 0.04 mmol, 0.6 equiv) in water (0.5 mL) was added dropwise. After stirring for an additional 30 min, the precipitate was filtered, washed with water (5 x 1 mL) and dried to yield the title compound as a cream solid (22 mg, 39%). ^1^H NMR (600 MHz, DMSO-d_6_): δ 9.71 (s, 1H), 7.49-7.47 (m, 2H), 7.27-7.22 (m, 4H), 7.19-7.16 (m, 2H), 7.13-7.10 (m, 2H), 7.07-7.04 (m, 3H), 7.00-6.97 (m, 1H), 6.18 (bs, 1H), 5.44 (bt, J = 4.5 Hz, 1H), 4.73 (bd, *J* = 3.5 Hz, 1H), 4.00-3.95 (m, 1H), 3.86-3.80 (m, 1H), 3.73-3.71 (m, 2H), 3.53-3.49 (m, 1H), 3.05 (t, *J* = 6.5 Hz, 2H), 2.04 (dd, *J* = 15.1, 4.4 Hz, 1H), 1.92 (dd, *J* = 15.1, 7.8 Hz, 1H), 1.58-1.52 (m, 1H), 1.50-1.43 (m, 1H), 1.40-1.35 (m, 1H), 1.24-1.20 (m, 1H); ^13^C NMR (151 MHz, DMSO-d_6_): δ 177.6, 164.2, 161.6 (d, *J* = 245 Hz), 139.5, 134.3, 133.3 (d, *J* = 8 Hz), 130.7, 129.7, 129.0, 128.5, 128.5 (d, *J* = 3 Hz), 127.6, 125.6, 122.9, 121.4, 119.1, 118.1, 115.3 (d, *J* = 21 Hz), 66.3, 66.1, 60.9, 44.0, 43.9, 40.9, 38.4, 28.3; FTIR (ATR): ʋ 3393, 1641, 1594, 1558, 1532 cm^-1^; HR-MS (ESI+): *m/z* calculated for C_32_H_34_N_2_O_6_F [M-0.5Ca+2H]^+^: 561.2401, found: 561.2390.

### (β*R*,δ*R*)-2-(4-Fluorophenyl)-β,δ-dihydroxy-5-(2-methylpropyl)-3-phenyl-4-[(phenylamino)carbonyl]-1*H*-pyrrole-1-heptanoic acid hemicalcium salt (5)

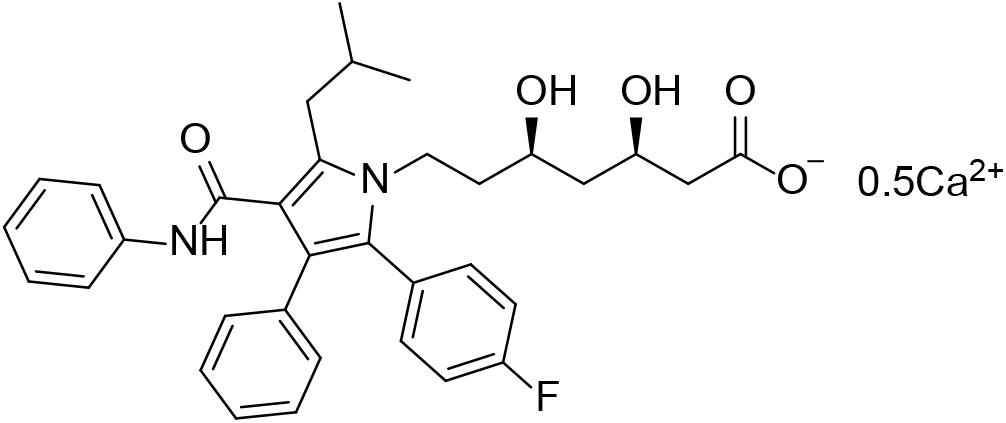

Prepared using General Procedure D (white powder, 206 mg, 46%). ^1^H NMR (600 MHz, DMSO-d_6_): δ 9.39 (s, 1H), 7.44-7.43 (m, 2H), 7.25-7.16 (m, 6H), 7.12-7.10 (m, 2H), 7.05-7.01 (m, 3H), 6.98-6.96 (m, 1H), 5.89 (bs, 1H), 4.74 (bs, 1H), 3.98-3.95 (m, 1H), 3.83-3.77 (m, 1H), 3.75-3.72 (m, 1H), 3.50-3.46 (m, 1H), 2.78-2.77 (m, 2H), 2.06 (dd, *J* = 15.2, 3.4 Hz, 1H), 1.93 (dd, *J* = 15.2, 8.0 Hz, 1H), 1.90-1.83 (m, 1H), 1.52-1.46 (m, 1H), 1.43-1.34 (m, 2H), 1.21-1.16 (m, 1H), 0.92-0.90 (m, 6H); ^13^C NMR (151 MHz, DMSO-d_6_): δ 178.2, 165.0, 161.6 (d, *J* = 245 Hz), 139.5, 134.8, 133.3 (d, *J* = 8 Hz), 132.8, 129.5, 128.6 (d, *J* = 3 Hz), 128.5, 127.7, 125.5, 122.8, 120.9, 119.3, 117.8, 115.4 (d, *J* = 21 Hz), 66.2, 66.0, 43.9, 43.7, 40.9, 38.4, 33.3, 29.2, 22.5, 22.5; FTIR (ATR): ʋ 3404, 1728, 1662, 1595, 1528 cm^-1^; HR-MS (ESI-): *m/z* calculated for C_34_H_36_N_2_O_5_F [M-0.5Ca]^-^: 571.2608, found: 571.2611.

### (β*R*,δ*R*)-5-Butyl-2-(4-fluorophenyl)-β,δ-dihydroxy-3-phenyl-4-[(phenylamino)carbonyl]-1*H*-pyrrole-1-heptanoic acid hemicalcium salt (6)

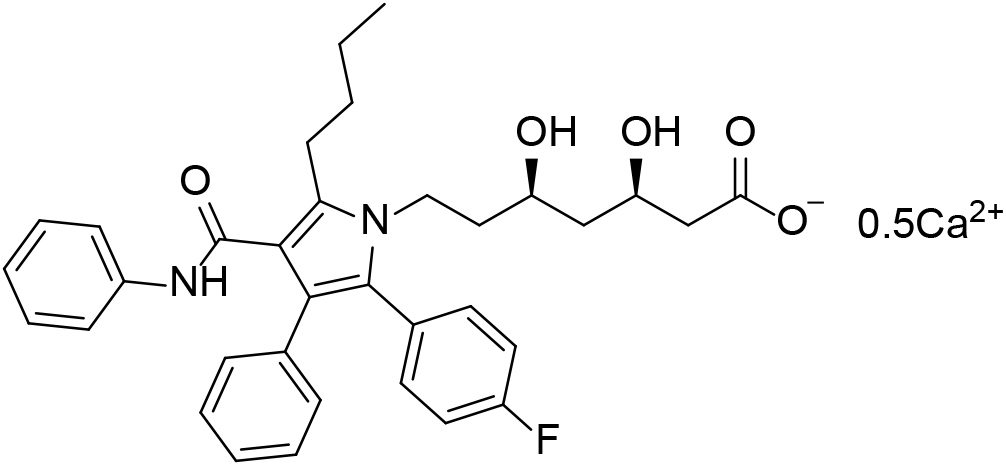

Prepared using General Procedure D (white powder, 372 mg, 84%). ^1^H NMR (500 MHz, DMSO-d_6_): δ 9.29 (s, 1H), 7.44-7.42 (m, 2H), 7.26-7.16 (m, 6H), 7.14-7.11 (m, 2H), 7.07-7.04 (m, 3H), 6.98-6.95 (m, 1H), 5.95 (bs, 1H), 4.75 (bs, 1H), 3.98-3.92 (m, 1H), 3.82-3.73 (m, 2H), 3.54-3.49 (m, 1H), 2.87-2.84 (m, 2H), 2.09-2.05 (m, 1H), 1.93 (dd, *J* = 15.2, 8.1 Hz, 1H), 1.61-1.52 (m, 3H), 1.49-1.43 (m, 1H), 1.42-1.31 (m, 3H), 1.23-1.18 (m, 1H), 0.86 (t, *J* = 7.3 Hz, 3H); ^13^C NMR (126 MHz, DMSO-d_6_): δ 178.1, 164.7, 161.6 (d, *J* = 245 Hz), 139.5, 134.8, 134.0, 133.3 (d, *J* = 8 Hz), 129.6, 128.5 (d, *J* = 3 Hz), 128.5, 128.3, 127.7, 125.6, 122.8, 120.8, 119.2, 117.1, 115.4 (d, *J* = 21 Hz), 66.3, 66.1, 43.9, 43.7, 40.8, 38.6, 32.2, 24.2, 22.1, 13.7; FTIR (ATR): ʋ 3403, 1662, 1594, 1558, 1531 cm^-1^; HR-MS (ESI+): *m/z* calculated for C_34_H_38_N_2_O_5_F [M-0.5Ca+2H]^+^: 573.2765, found: 573.2761.

### (β*R*,δ*R*)-5-Cyclobutyl-2-(4-fluorophenyl)-β,δ-dihydroxy-5-cyclobutyl-3-phenyl-4-[(phenylamino)carbonyl]-1*H*-pyrrole-1-heptanoic acid hemicalcium salt (7)

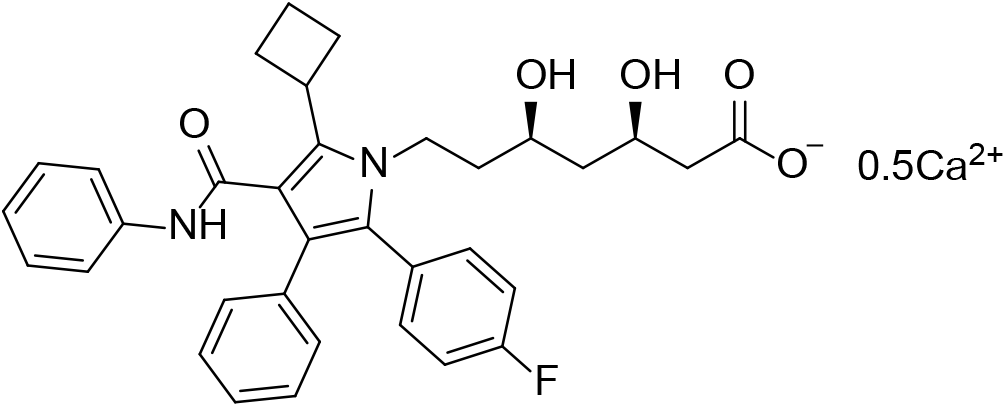

Prepared using General Procedure D (off-white powder, 127 mg, 29%). ^1^H NMR (500 MHz, DMSO-d_6_): δ 10.06 (s, 1H), 7.59-7.57 (m, 2H), 7.25-7.16 (m, 6H), 7.10-7.05 (m, 4H), 7.02-6.98 (m, 2H), 5.93 (bs, 1H), 4.72 (bs, 1H), 3.94-3.87 (m, 1H), 3.80-3.69 (m, 3H), 3.53-3.49 (m, 1H), 2.42-2.27 (m, 4H), 2.09-2.05 (m, 1H), 1.96-1.85 (m, 2H), 1.71-1.66 (m, 1H), 1.57-1.50 (m, 1H), 1.48-1.36 (m, 2H), 1.24-1.19 (m, 1H); ^13^C NMR (126 MHz, DMSO-d_6_): δ 178.2, 165.9, 161.6 (d, *J* = 245 Hz), 139.5, 134.9, 133.3 (d, *J* = 8 Hz), 133.3, 129.2, 128.6 (d, *J* = 3 Hz), 128.5, 127.9, 127.6, 125.3, 123.0, 120.5, 119.3, 118.3, 115.4 (d, *J* = 21 Hz), 66.3, 66.2, 44.0, 43.8, 41.1, 38.5, 32.5, 28.9, 28.8, 18.5; FTIR (ATR): ʋ 3407, 1661, 1594, 1559, 1508 cm^-1^; HR-MS (ESI-): *m/z* calculated for C_34_H_34_N_2_O_5_F [M-0.5Ca]^-^: 569.2452, found: 569.2448.

### (β*R*,δ*R*)-5-Cyclopentyl-2-(4-fluorophenyl)-β,δ-dihydroxy-3-phenyl-4-[(phenylamino)carbonyl]-1*H*-pyrrole-1-heptanoic acid hemicalcium salt (8)

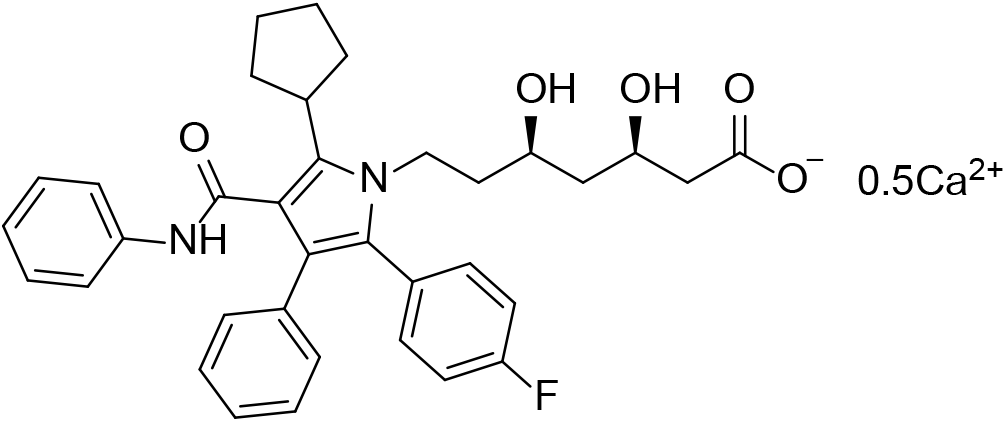

Prepared using General Procedure D (white powder, 159 mg, 36%). ^1^H NMR (600 MHz, DMSO-d_6_): δ 9.76 (s, 1H), 7.49-7.47 (m, 2H), 7.25-7.16 (m, 6H), 7.09-7.05 (m, 4H), 7.00-6.96 (m, 2H), 5.95 (bs, 1H), 4.73 (bs, 1H), 3.98-3.94 (m, 1H), 3.81-3.74 (m, 2H), 3.55-3.51 (m, 1H), 3.27-3.22 (m, 1H), 2.09-1.92 (m, 6H), 1.71-1.65 (m, 2H), 1.63-1.49 (m, 4H), 1.42-1.37 (m, 1H), 1.26-1.21 (m, 1H); ^13^C NMR (151 MHz, DMSO-d_6_): δ 178.2, 166.0, 161.6 (d, *J* = 245 Hz), 139.4, 134.9, 133.9, 133.4 (d, *J* = 8 Hz), 129.2, 128.8 (d, *J* = 3 Hz), 128.4, 127.6, 127.6, 125.4, 123.0, 120.8, 119.4, 117.6, 115.4 (d, *J* = 21 Hz), 66.3, 66.2, 43.9, 43.7, 41.0, 36.6, 32.5, 25.5, 25.5; FTIR (ATR): ʋ 3395, 1652, 1594, 1558, 1508 cm^-1^; HR-MS (ESI-): *m/z* calculated for C_35_H_36_N_2_O_5_F [M-0.5Ca]^-^: 583.2608, found: 583.2594.

### (β*R*,δ*R*)-5-Cyclohexyl-2-(4-fluorophenyl)-β,δ-dihydroxy-3-phenyl-4-[(phenylamino)carbonyl]-1*H*-pyrrole-1-heptanoic acid hemicalcium salt (9)

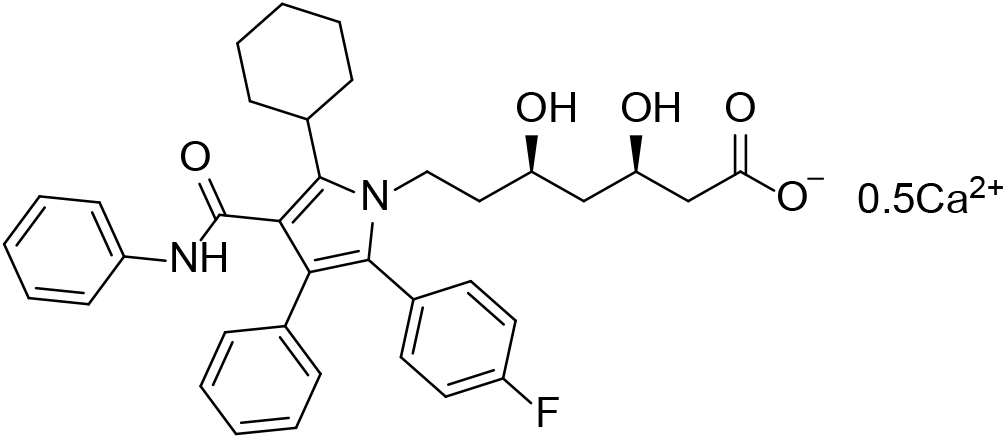

Prepared using General Procedure D (white powder, 57 mg, 23%). ^1^H NMR (500 MHz, DMSO-d_6_): δ 9.82 (s, 1H), 7.51-7.50 (m, 2H), 7.25-7.16 (m, 6H), 7.07-7.05 (m, 4H), 7.00-6.97 (m, 2H), 6.51 (bs, 1H), 4.80 (bs, 1H), 3.95-3.89 (m, 1H), 3.79-3.72 (m, 2H), 3.57-3.53 (m, 1H), 2.85-2.80 (m, 1H), 2.07-2.03 (m, 1H), 1.91-1.80 (m, 5H), 1.76-1.71 (m, 2H), 1.65-1.60 (m, 2H), 1.57-1.49 (m, 1H), 1.43-1.29 (m, 3H), 1.25-1.20 (m, 1H), 1.12-1.04 (m, 1H); ^13^C NMR (126 MHz, DMSO-d_6_): δ 177.2, 166.3, 161.6 (d, *J* = 245 Hz), 139.5, 135.1, 135.0, 133.4 (d, *J* = 8 Hz), 129.1, 128.8 (d, *J* = 3 Hz), 128.4, 127.6, 127.2, 125.3, 122.9, 120.5, 119.5, 117.8, 115.4 (d, *J* = 21 Hz), 66.4, 66.3, 43.8, 40.7, 36.3, 32.1, 26.7, 25.6; FTIR (ATR): ʋ 3395, 1652, 1594, 1558, 1508 cm^-1^; HR-MS (ESI-): *m/z* calculated for C_36_H_38_N_2_O_5_F [M-0.5Ca]^-^: 597.2765, found: 597.2770.

---

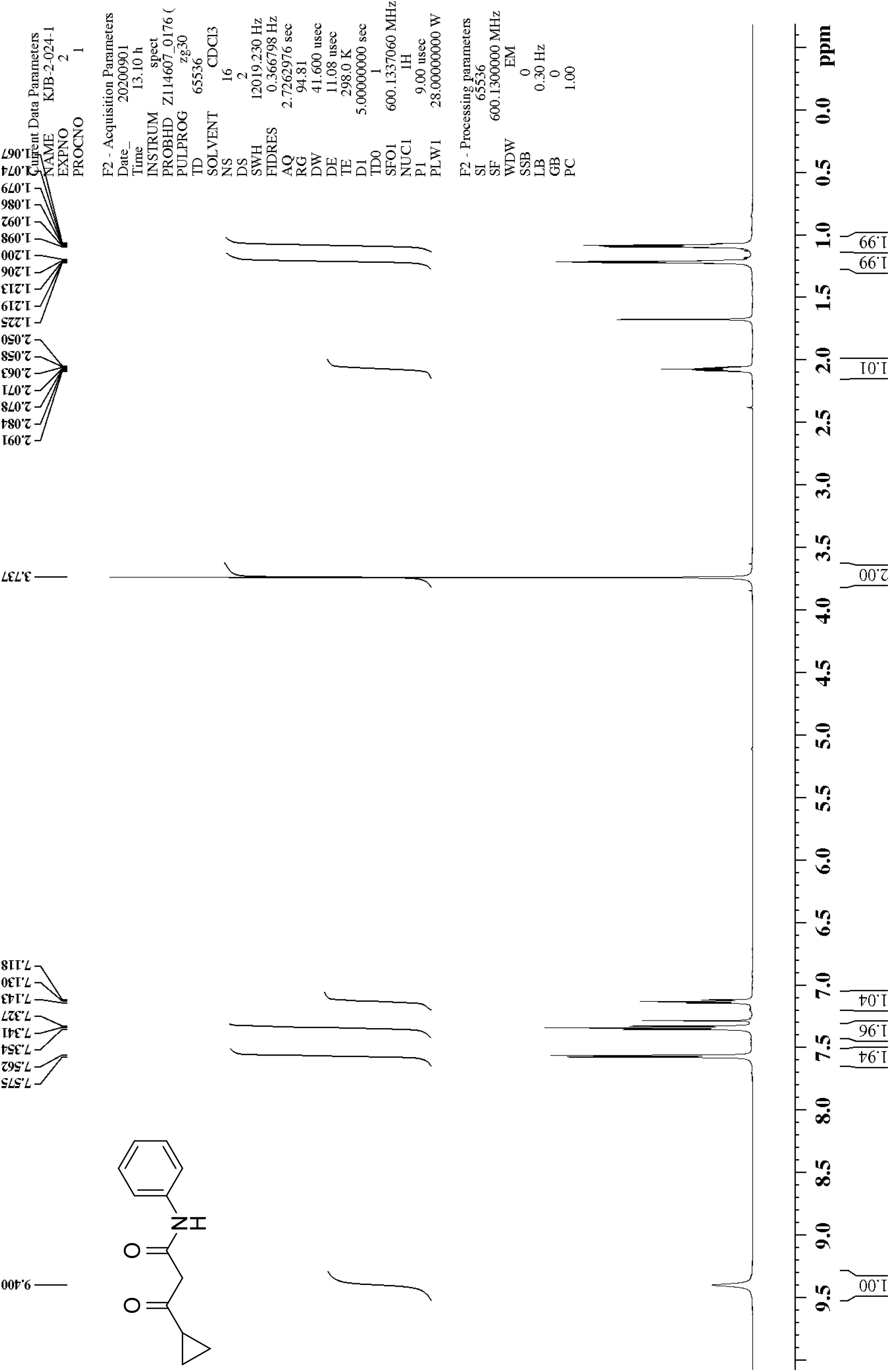
^1^H NMR of β-oxo-*N*-phenylcyclopropanepropanamide (10)

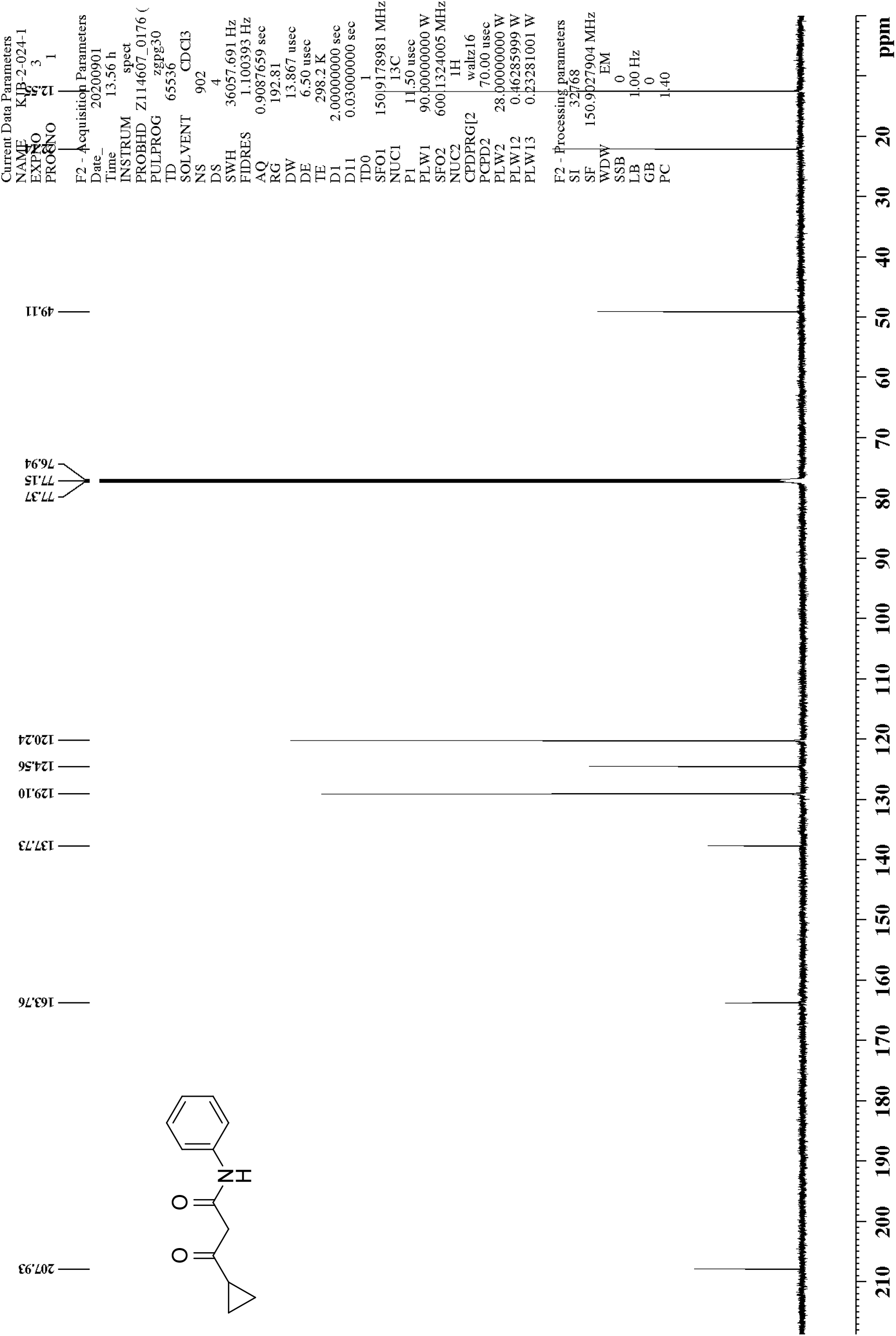
^13^C NMR of β-oxo-*N-*phenylcyclopropanepropanamide (10)

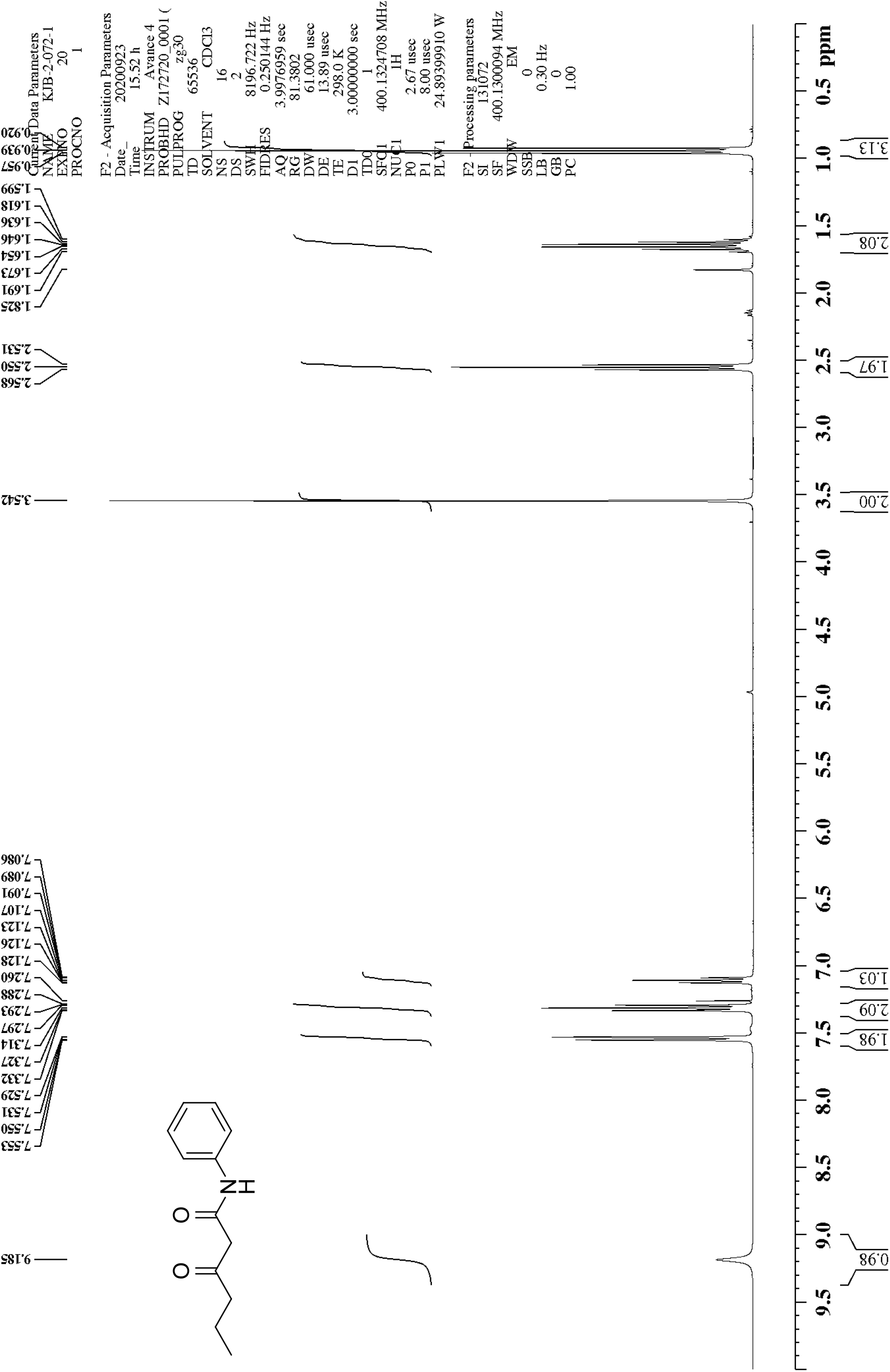
^1^H NMR of 3-oxo-*N-*phenylhexanamide (12)

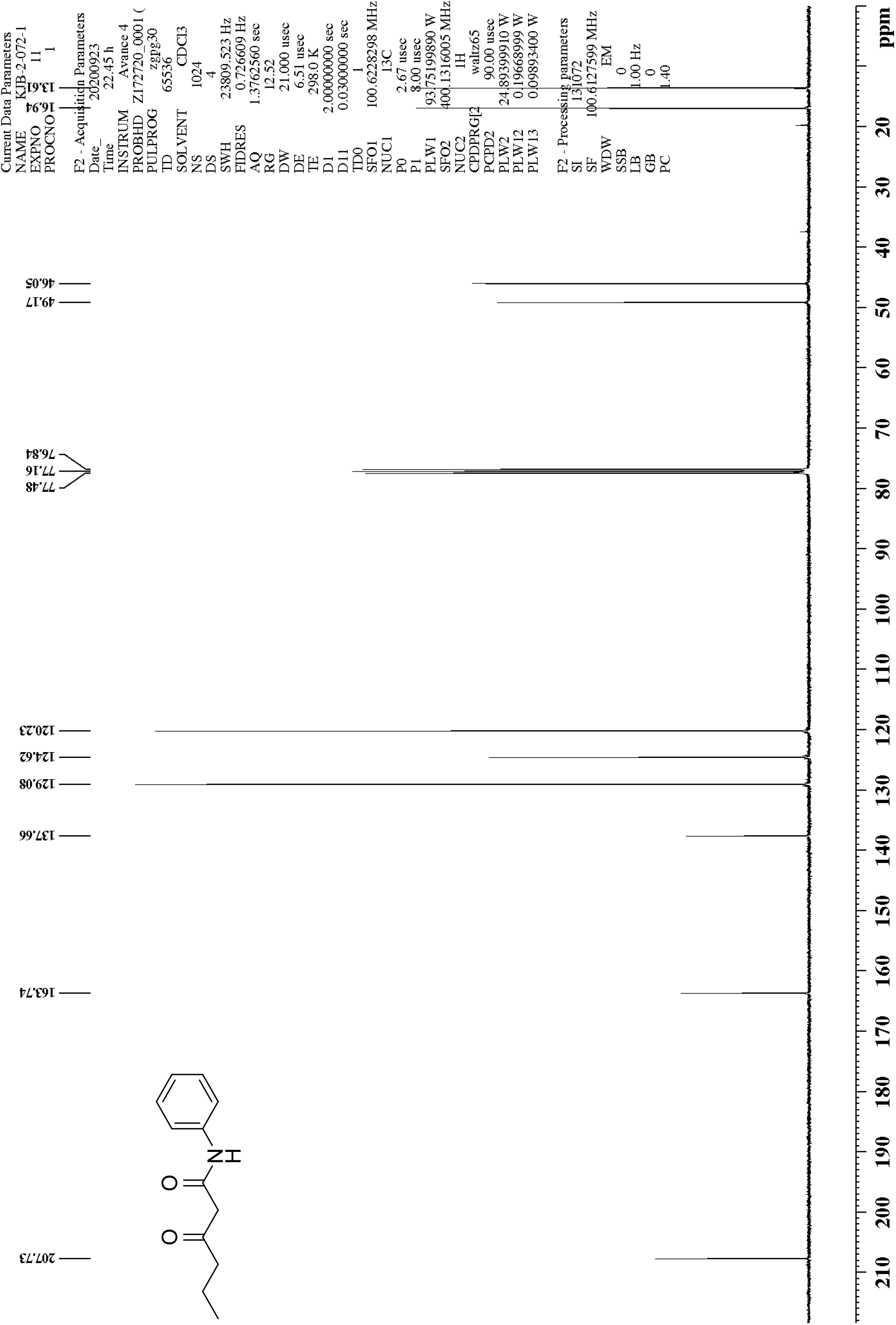
^13^C NMR of 3-oxo-*N-*phenylhexanamide (12)

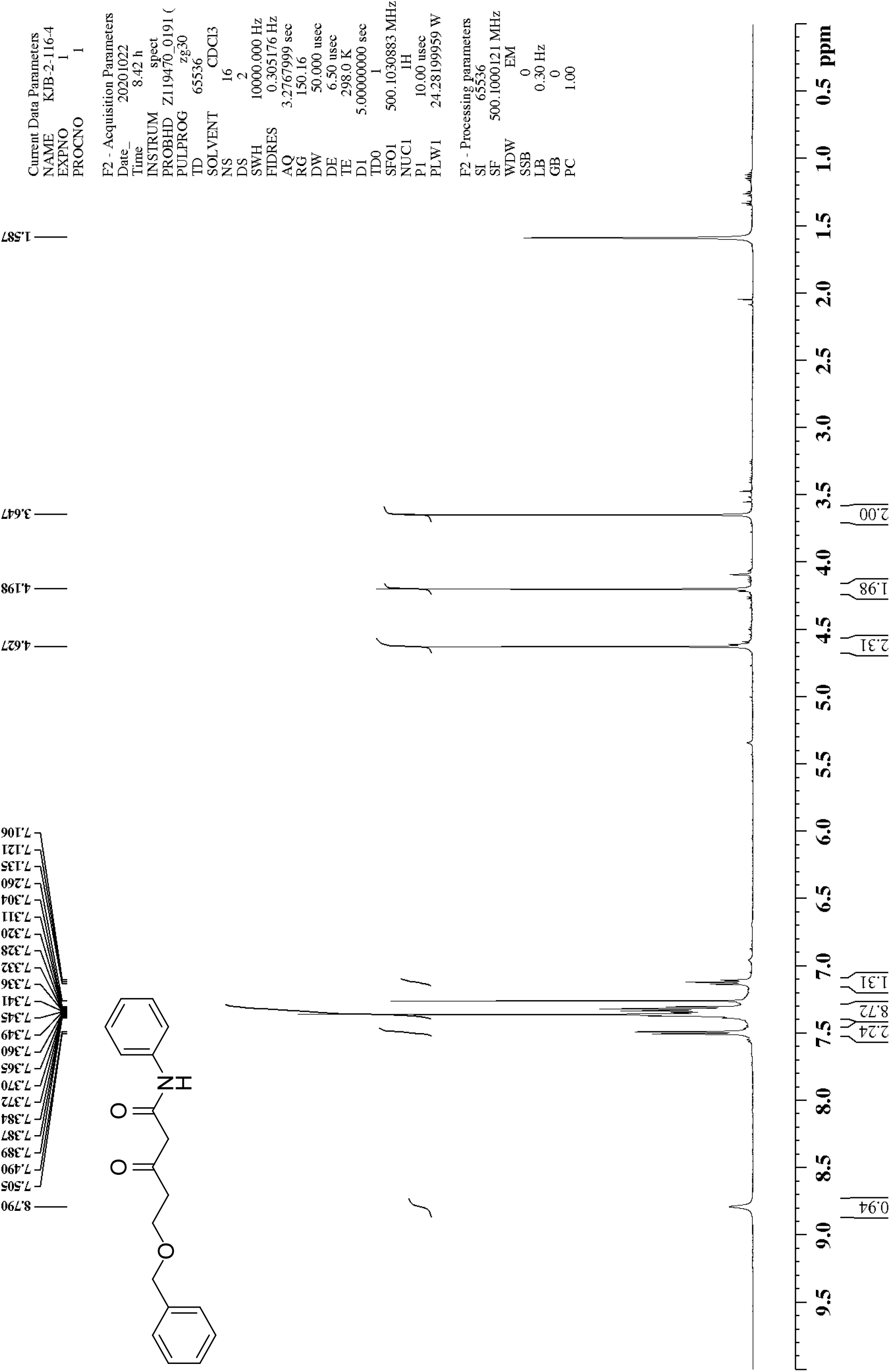
^1^H NMR of 3-oxo-*N-*phenyl-5-(benzyloxy)pentanamide (13)

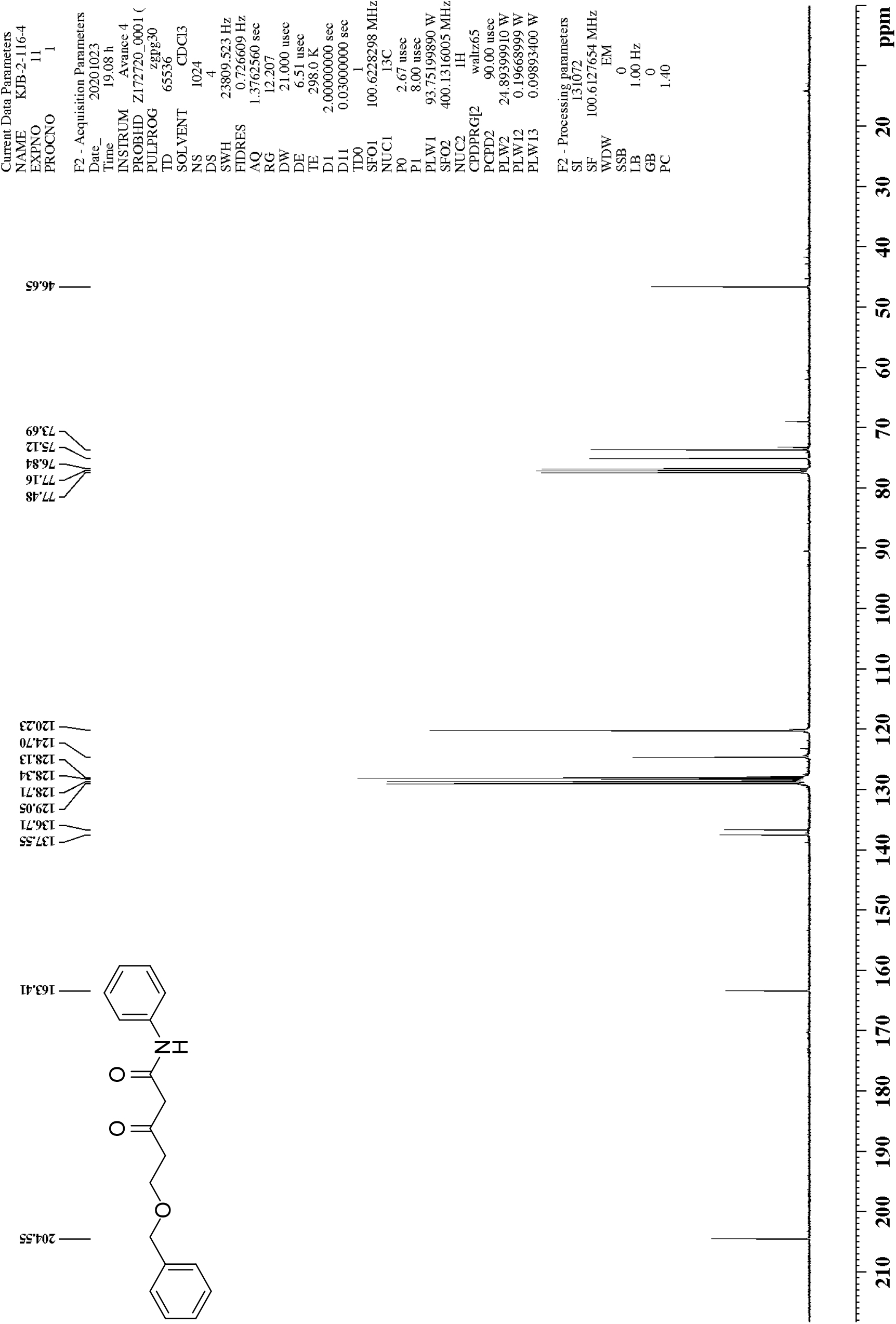
^13^C NMR of 3-oxo-*N-*phenyl-5-(benzyloxy)pentanamide (13)

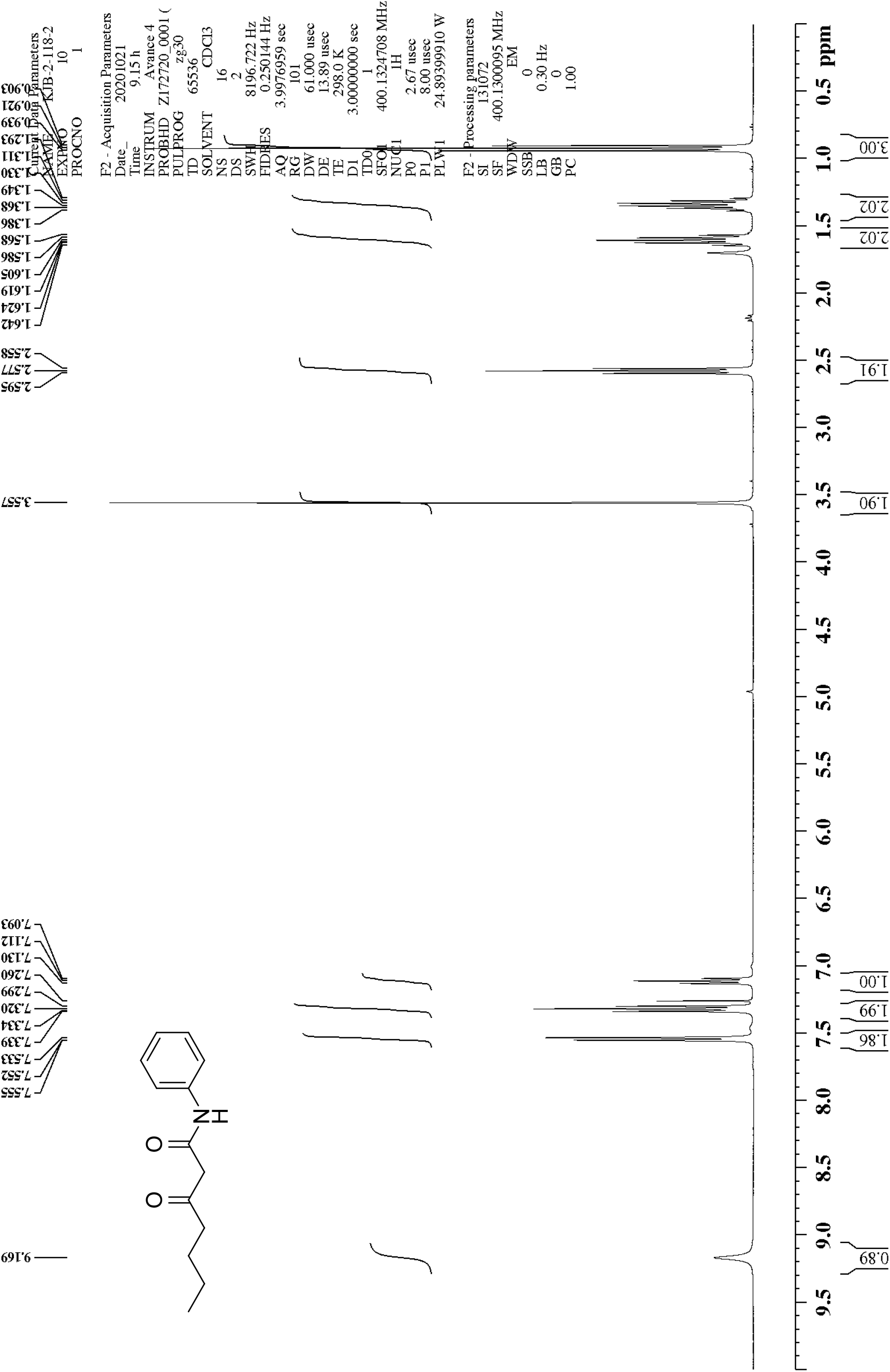
^1^H NMR of 3-oxo-*N-*phenylheptanamide (15)

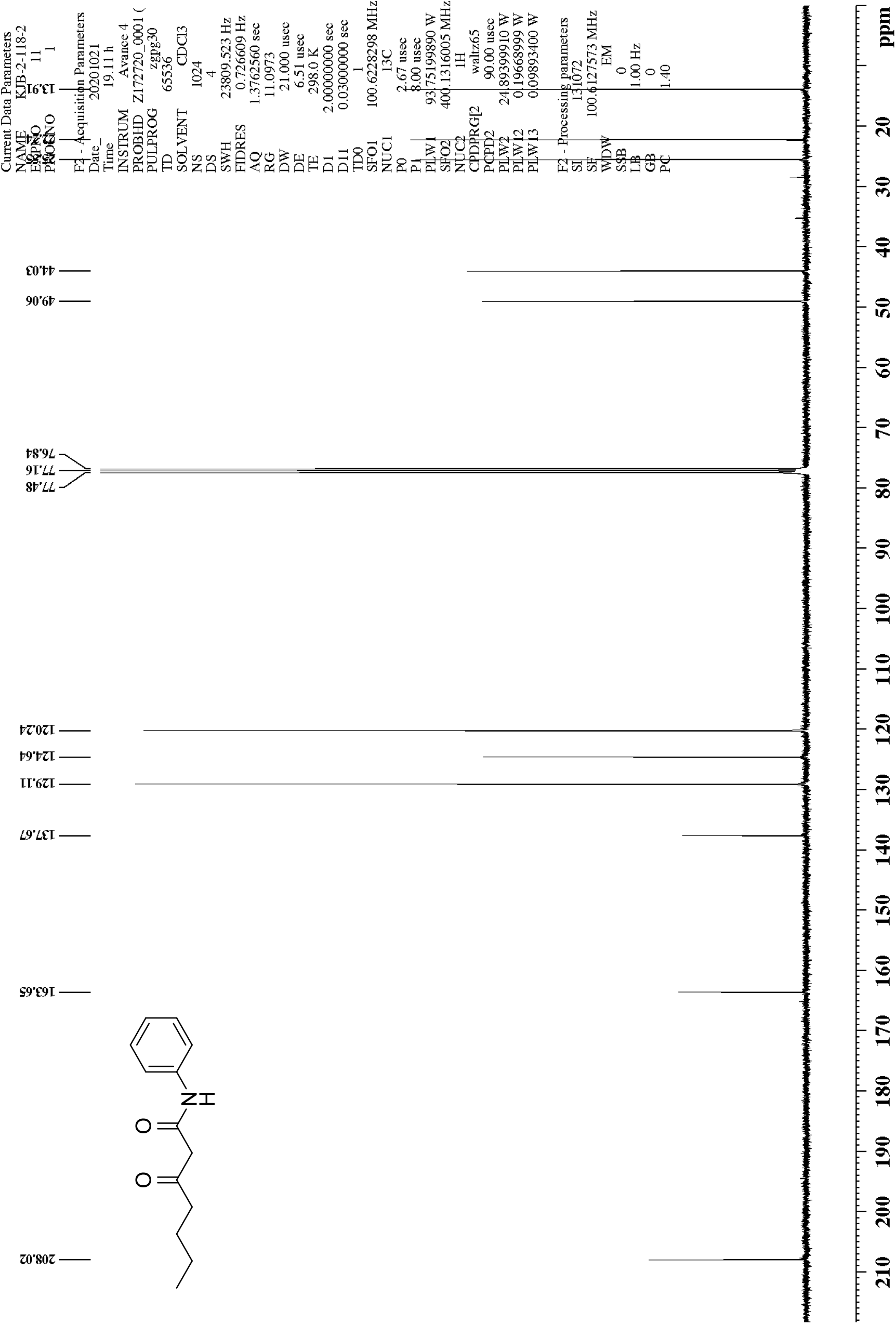
^13^C NMR of 3-oxo-*N-*phenylheptanamide (15)

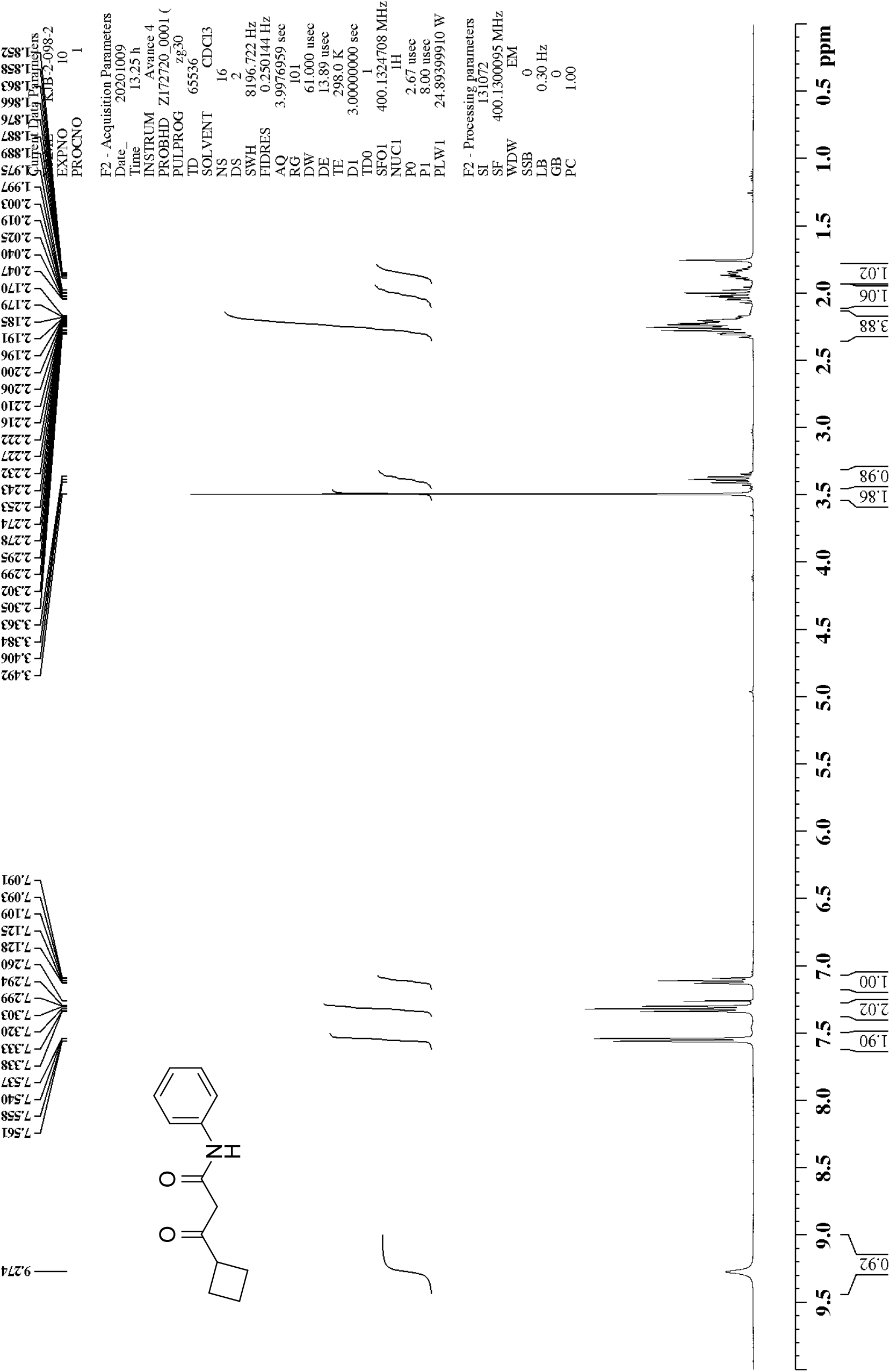
^1^H NMR of β-oxo-*N-*phenylcyclobutanepropanamide (16)

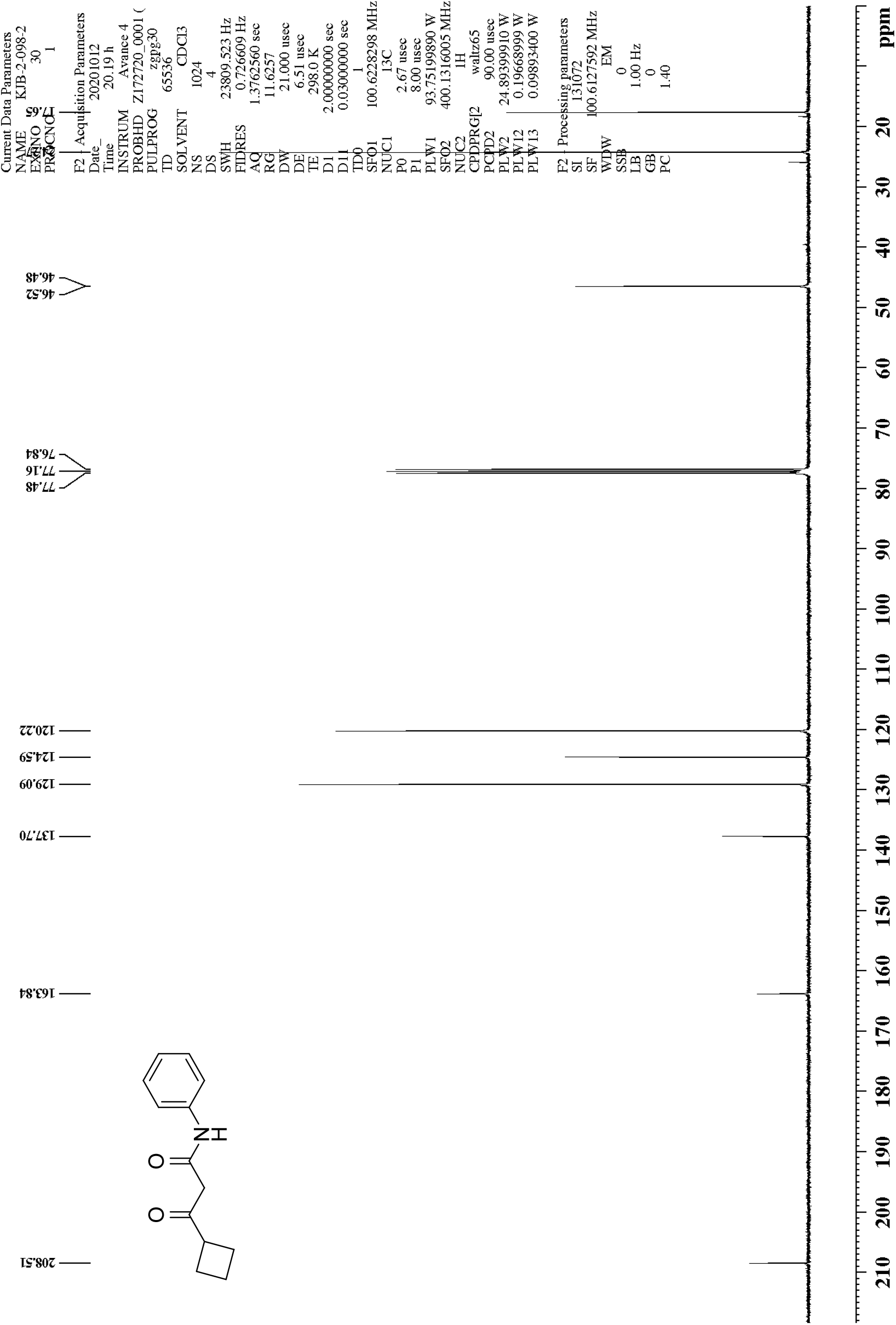
^13^C NMR of β-oxo-*N-*phenylcyclobutanepropanamide (16)

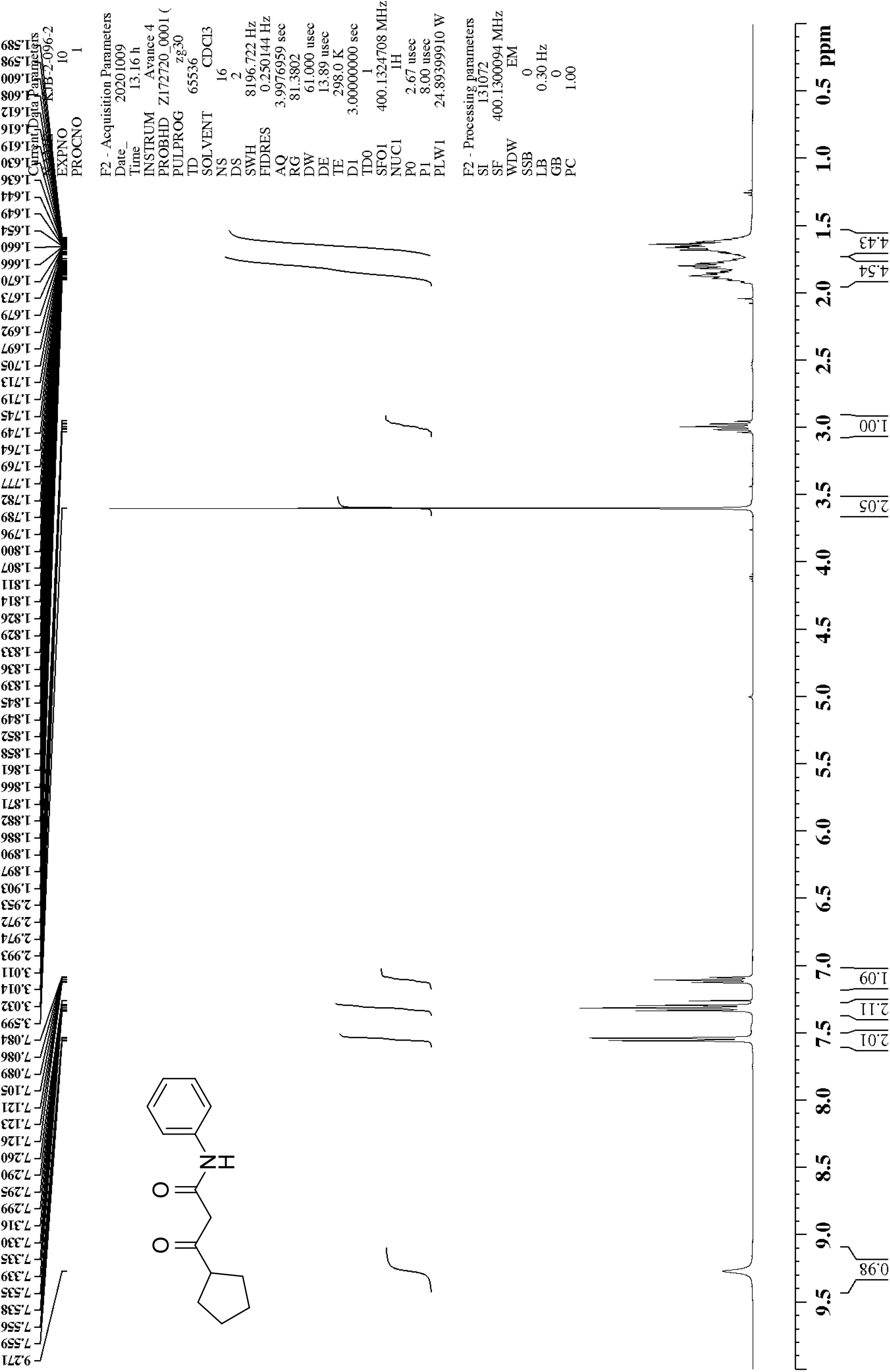
^1^H NMR of β-oxo-*N-*phenylcyclopentanepropanamide (17)

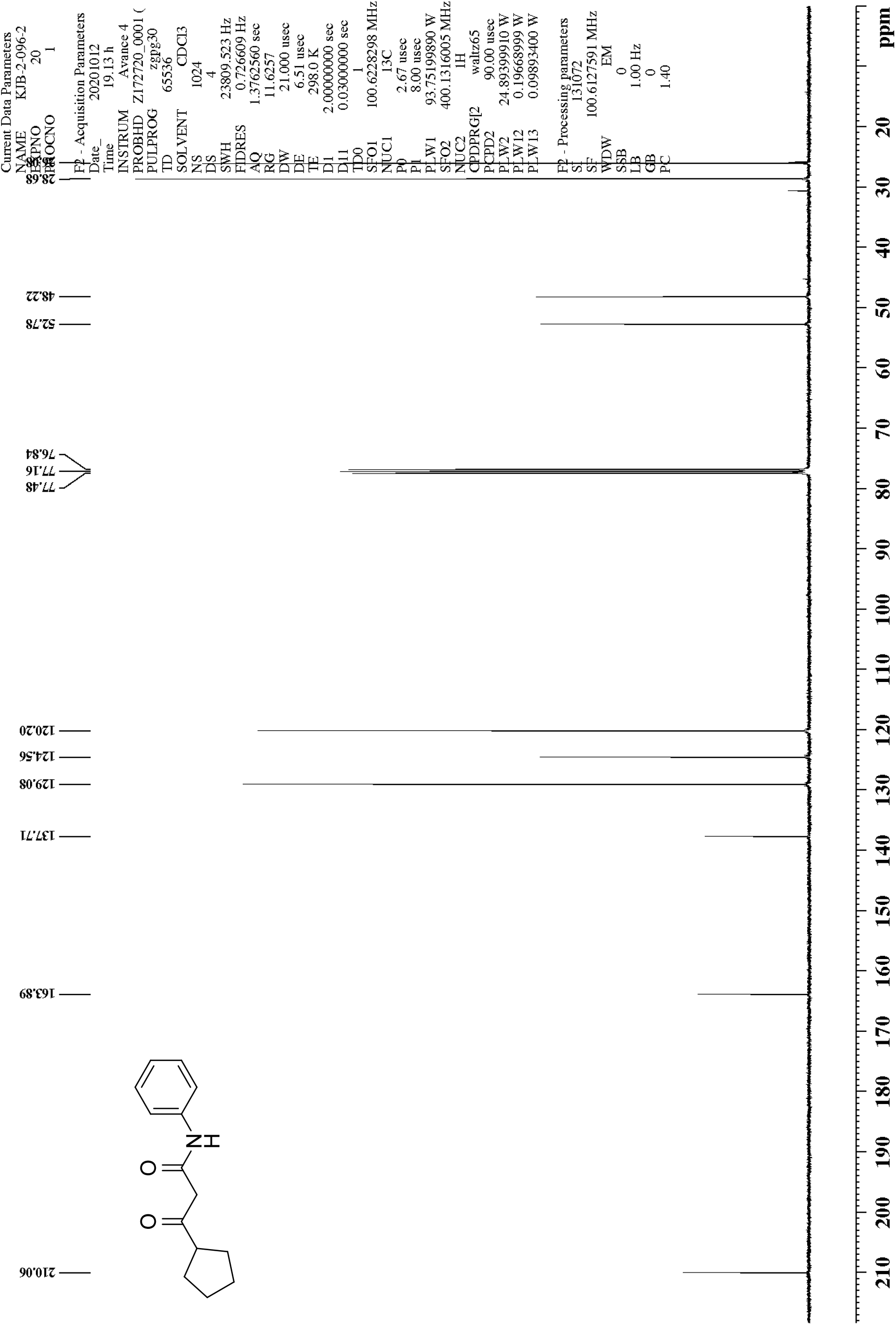
^13^C NMR of β-oxo-*N-*phenylcyclopentanepropanamide (17)

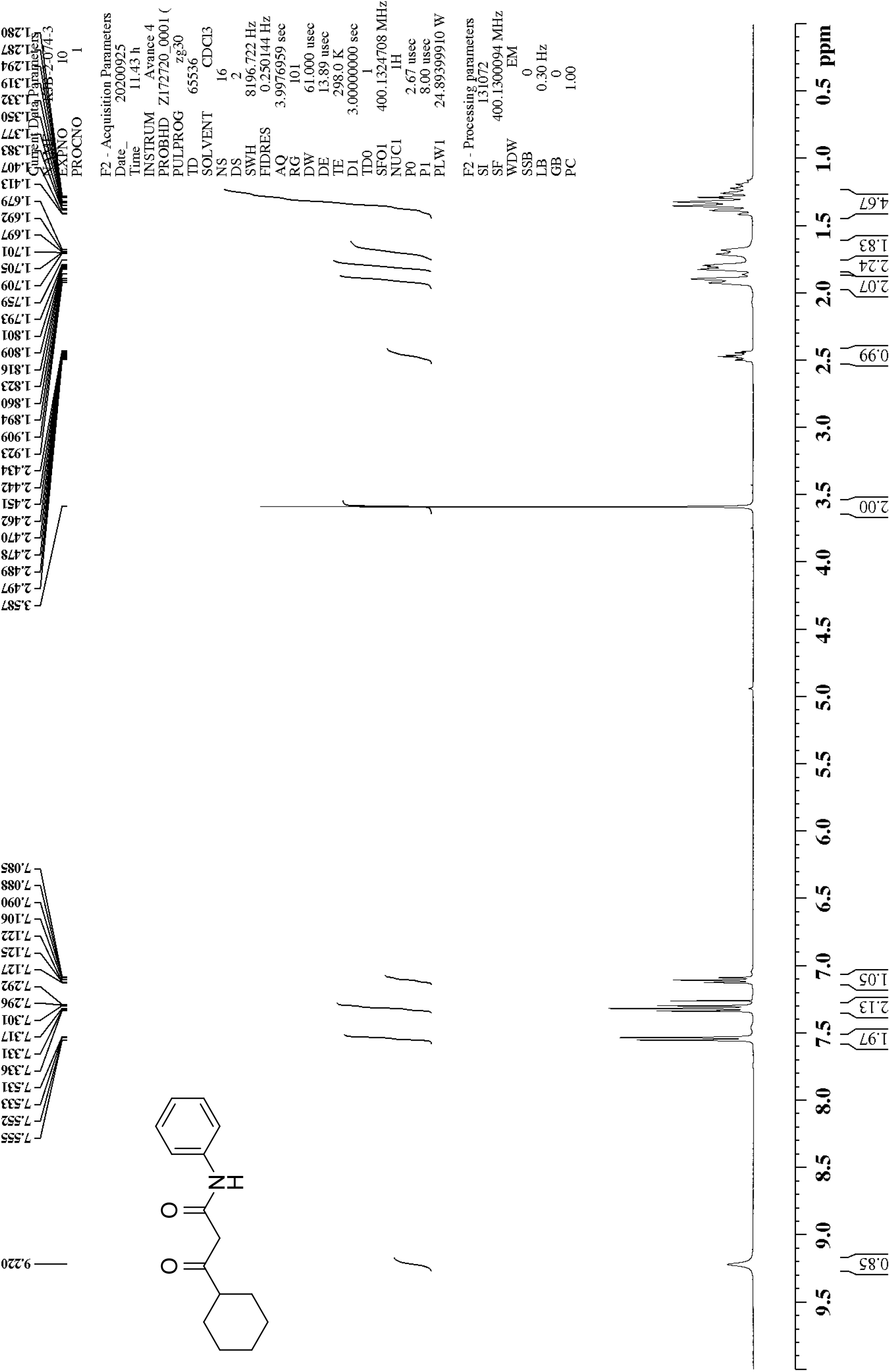
^1^H NMR of β-oxo-*N-*phenylcyclohexanepropanamide (18)

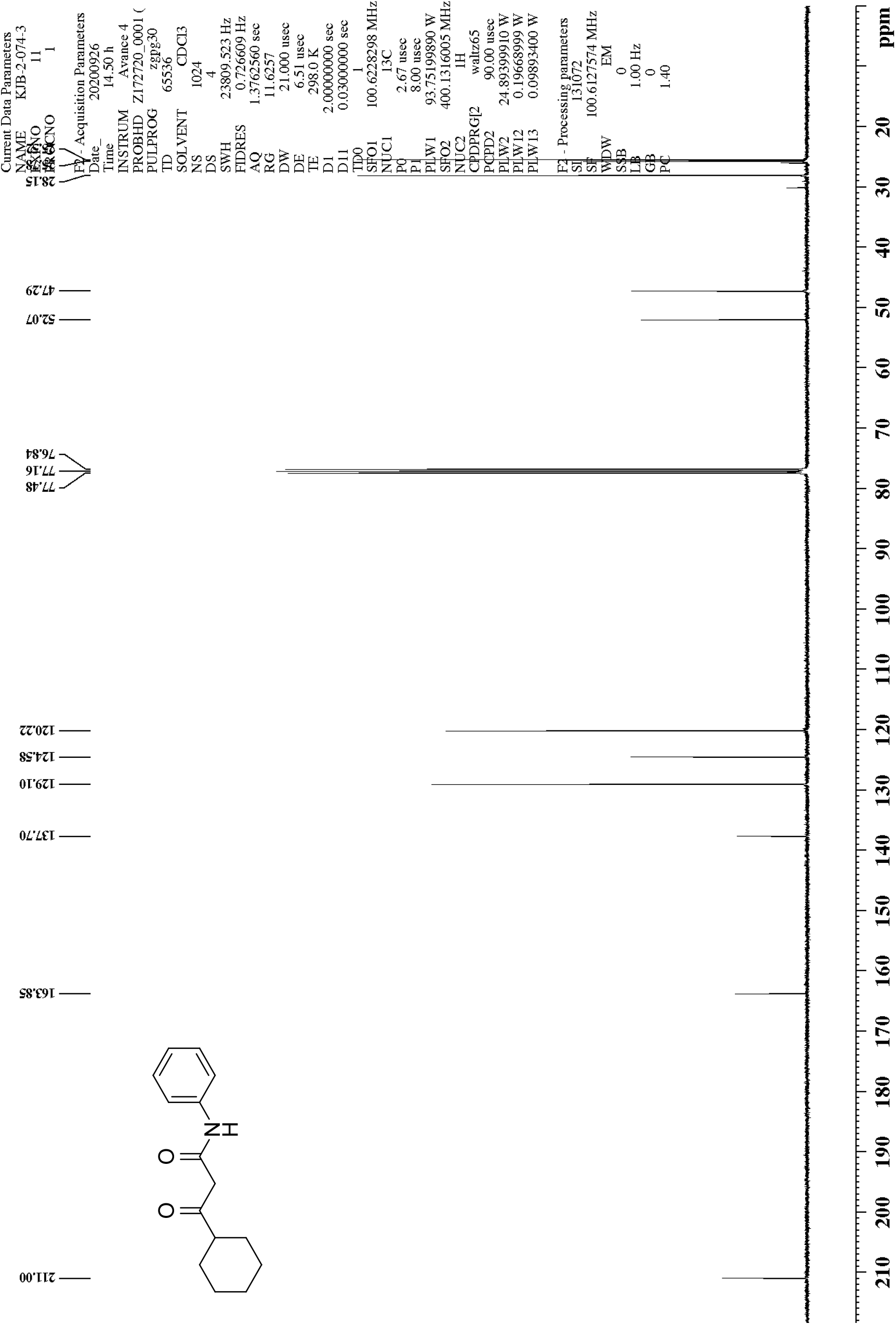
^13^C NMR of β-oxo-*N-*phenylcyclohexanepropanamide (18)

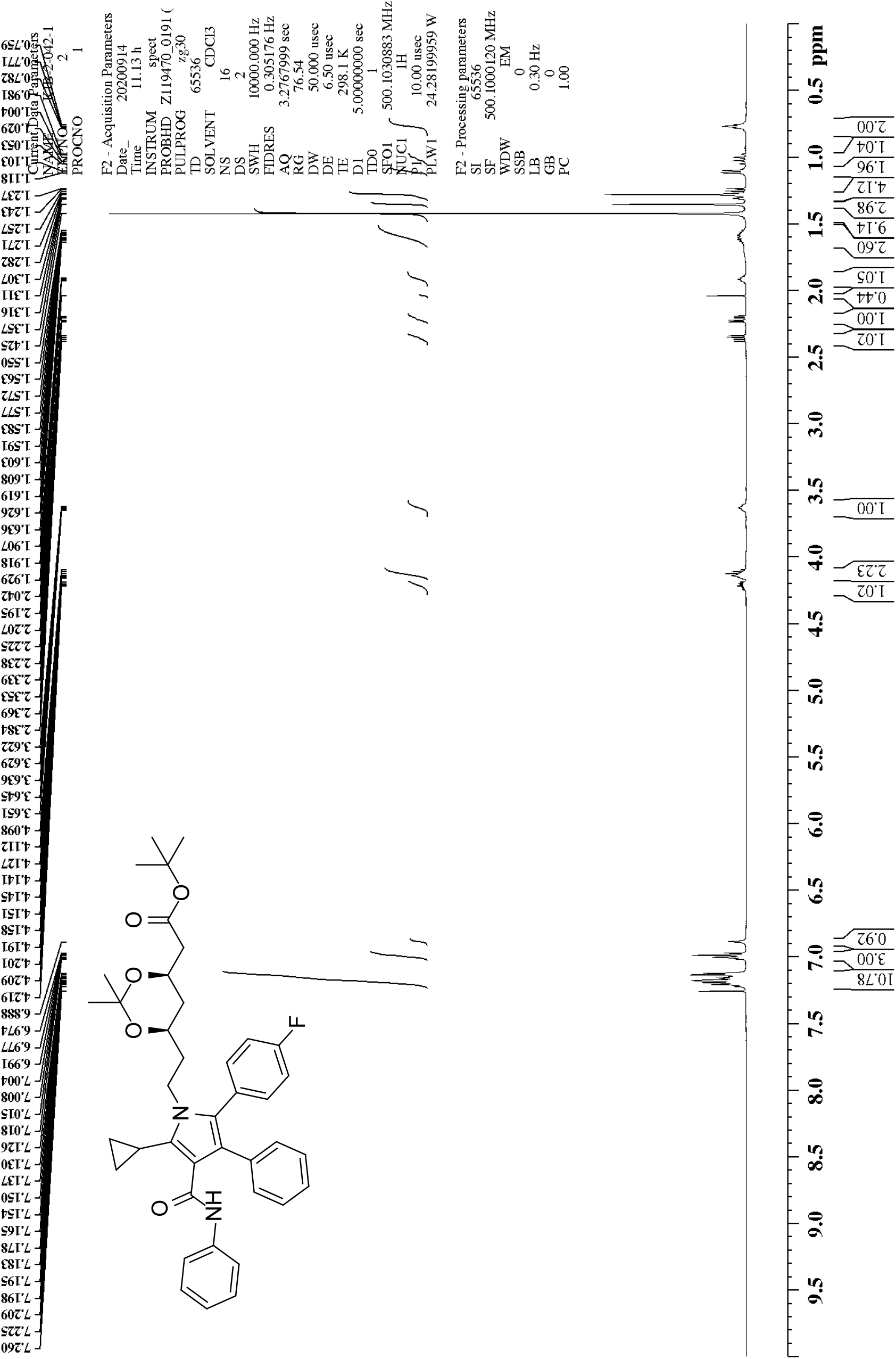
^1^H NMR of 1,1-dimethylethyl (4*R*,6*R*)-6-[2-[5-cyclopropyl-2-(4-fluorophenyl)-3-phenyl-4-[(phenylamino)carbonyl]-1*H*-pyrrol-1-yl]ethyl]-2,2-dimethyl-1,3-dioxane-4-acetate (19)

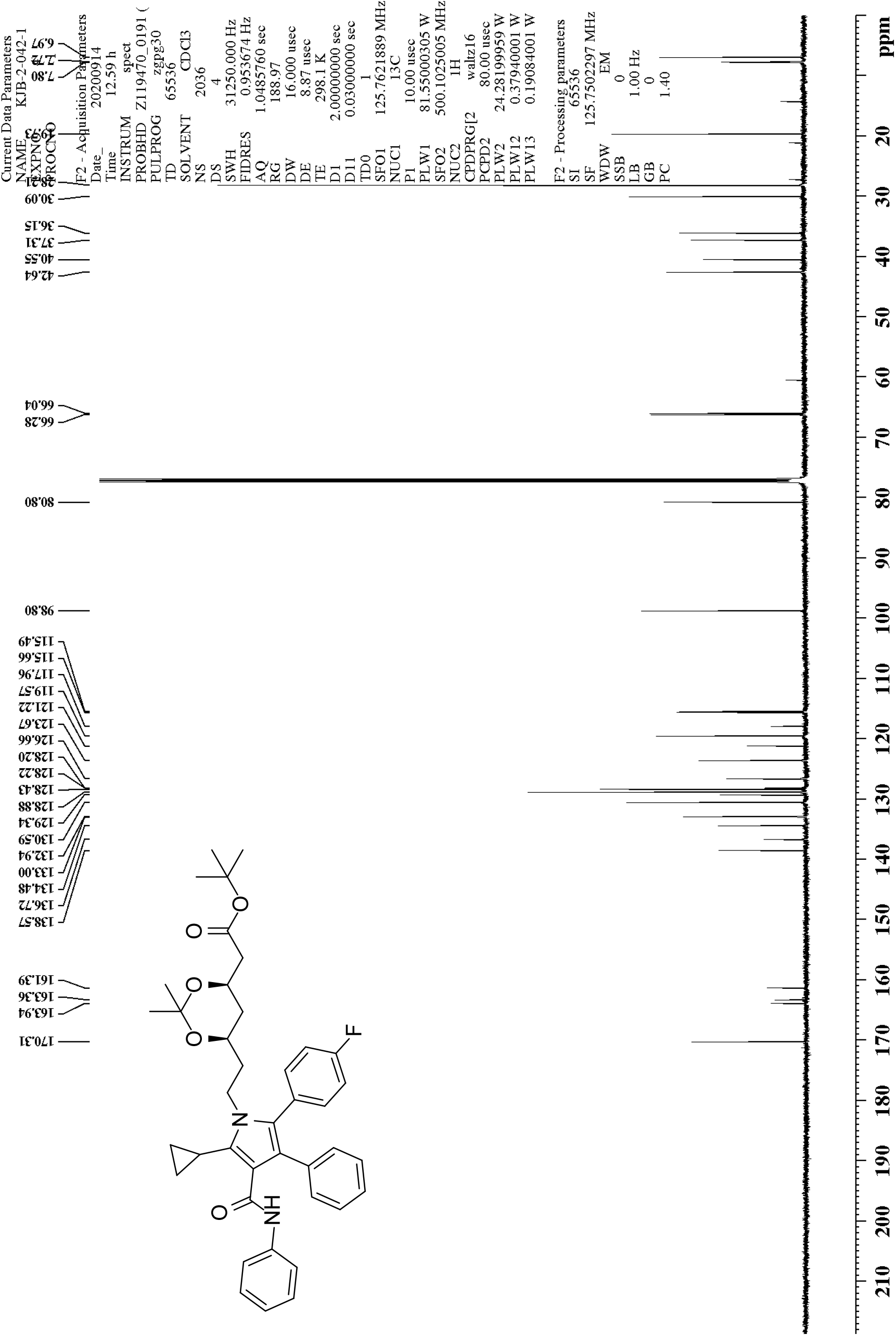
^13^C NMR of 1,1-dimethylethyl (4*R*,6*R*)-6-[2-[5-cyclopropyl-2-(4-fluorophenyl)-3-phenyl-4-[(phenylamino)carbonyl]-1*H*-pyrrol-1-yl]ethyl]-2,2-dimethyl-1,3-dioxane-4-acetate (19)

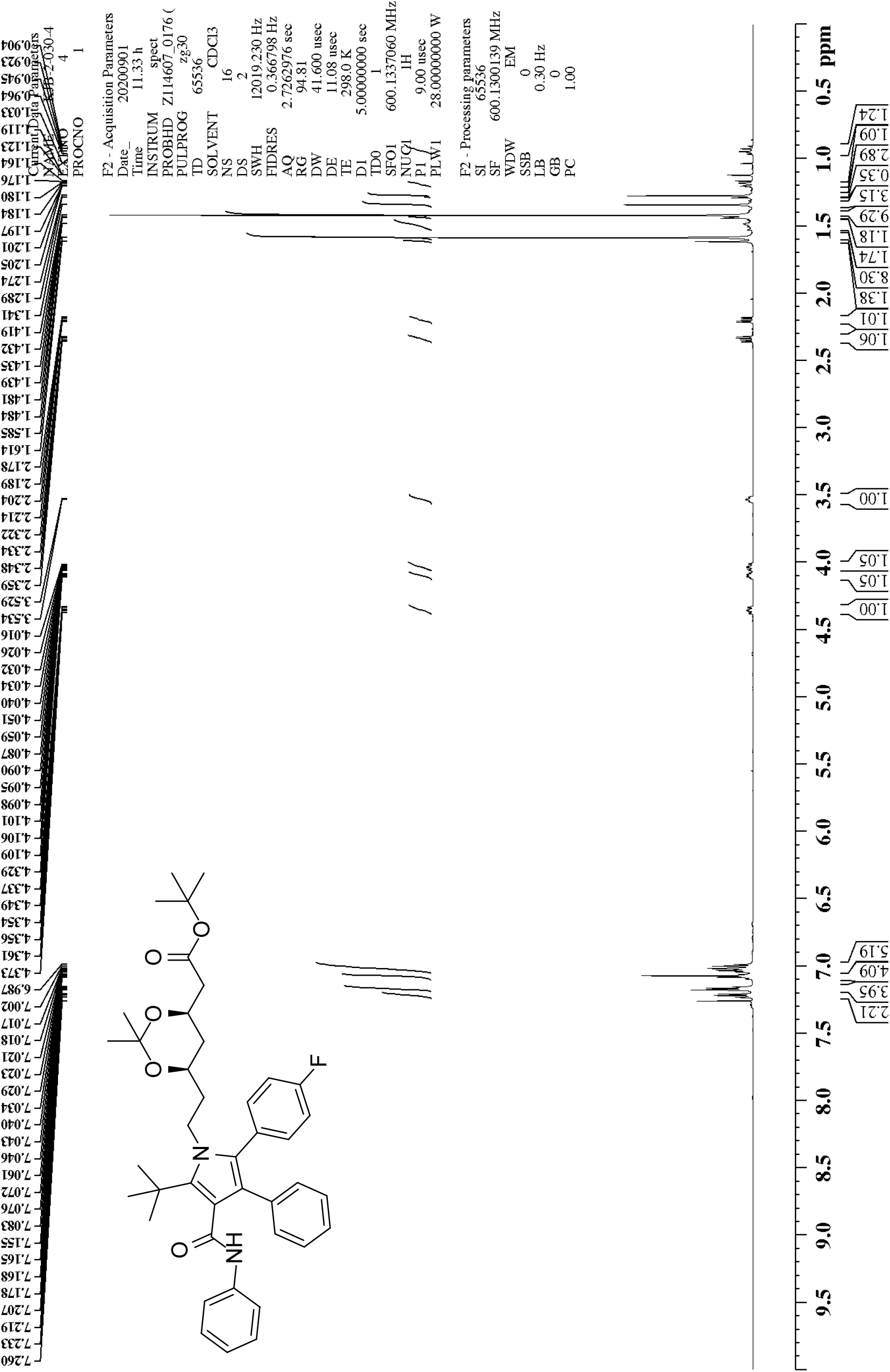
^1^H NMR of 1,1-dimethylethyl (4*R*,6*R*)-6-[2-[2-(4-fluorophenyl)-5-(1,1-dimethylethyl)-3-phenyl-4-[(phenylamino)carbonyl]-1*H*-pyrrol-1-yl]ethyl]-2,2-dimethyl-1,3-dioxane-4-acetate (20)

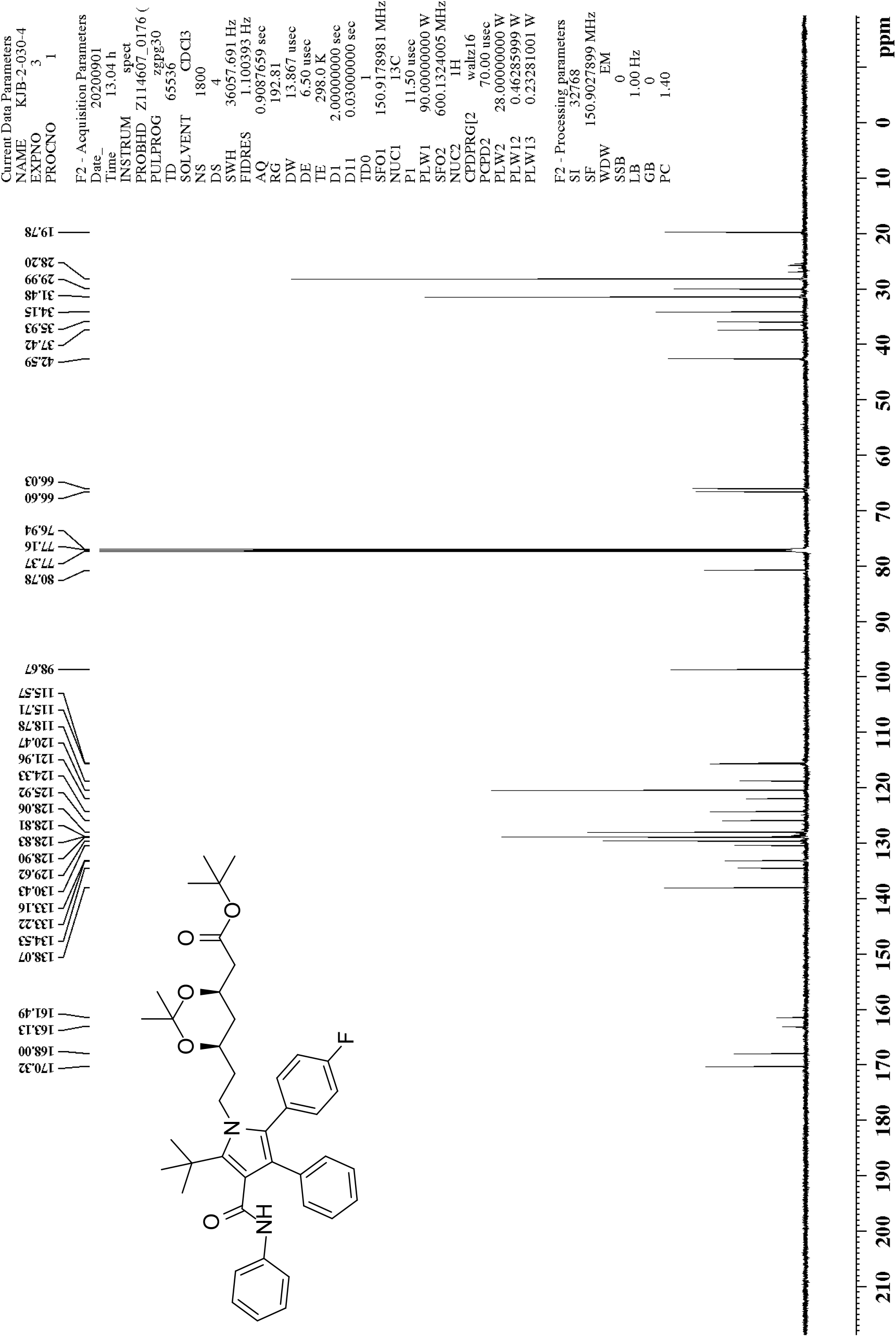
^13^C NMR of 1,1-dimethylethyl (4*R*,6*R*)-6-[2-[2-(4-fluorophenyl)-5-(1,1-dimethylethyl)-3-phenyl-4-[(phenylamino)carbonyl]-1*H*-pyrrol-1-yl]ethyl]-2,2-dimethyl-1,3-dioxane-4-acetate (20)

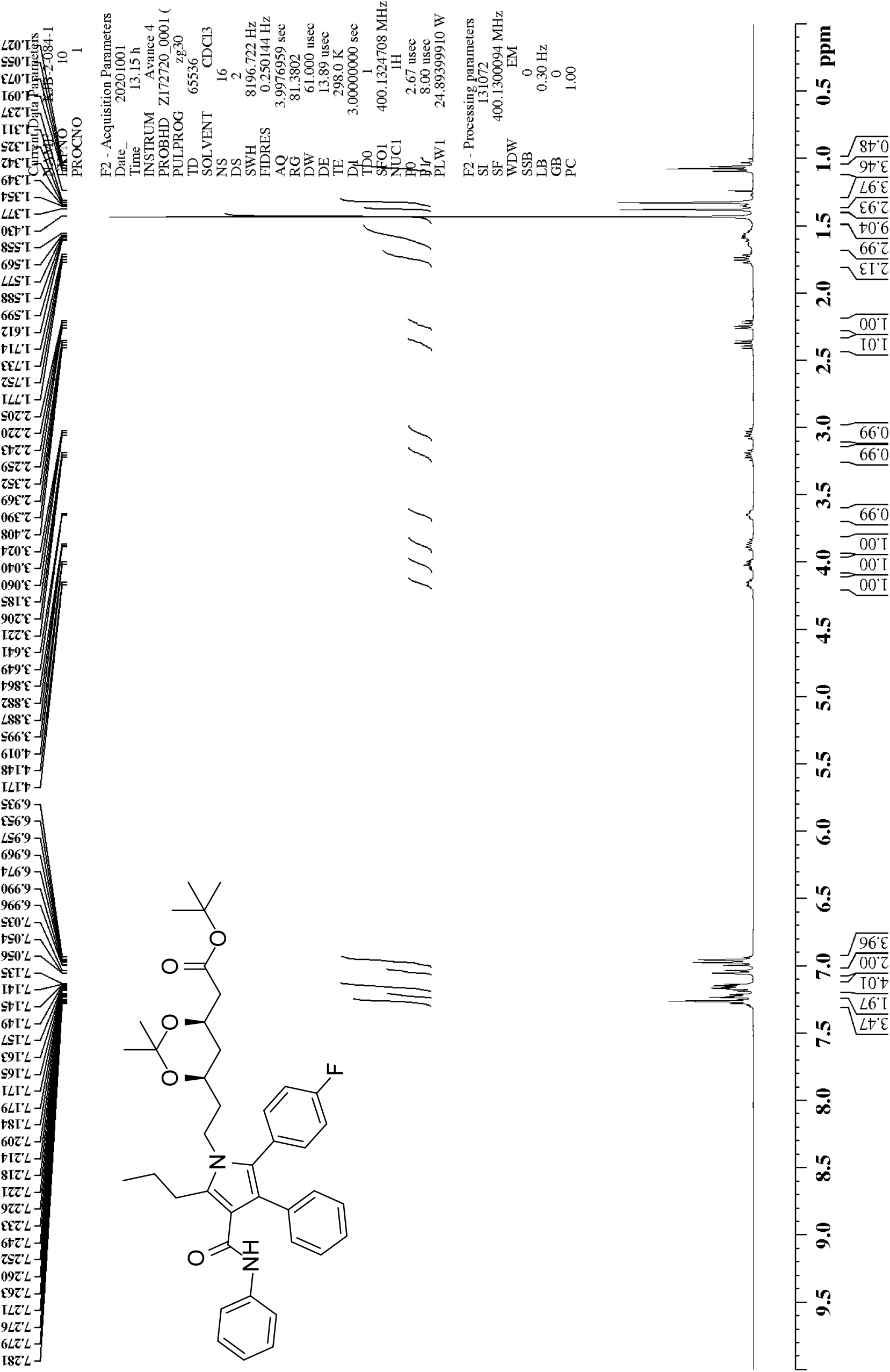
^1^H NMR of 1,1-dimethylethyl (4*R*,6*R*)-6-[2-[2-(4-fluorophenyl)-3-phenyl-4-[(phenylamino)carbonyl-5-propyl]-1*H*-pyrrol-1-yl]ethyl]-2,2-dimethyl-1,3-dioxane-4-acetate (21)

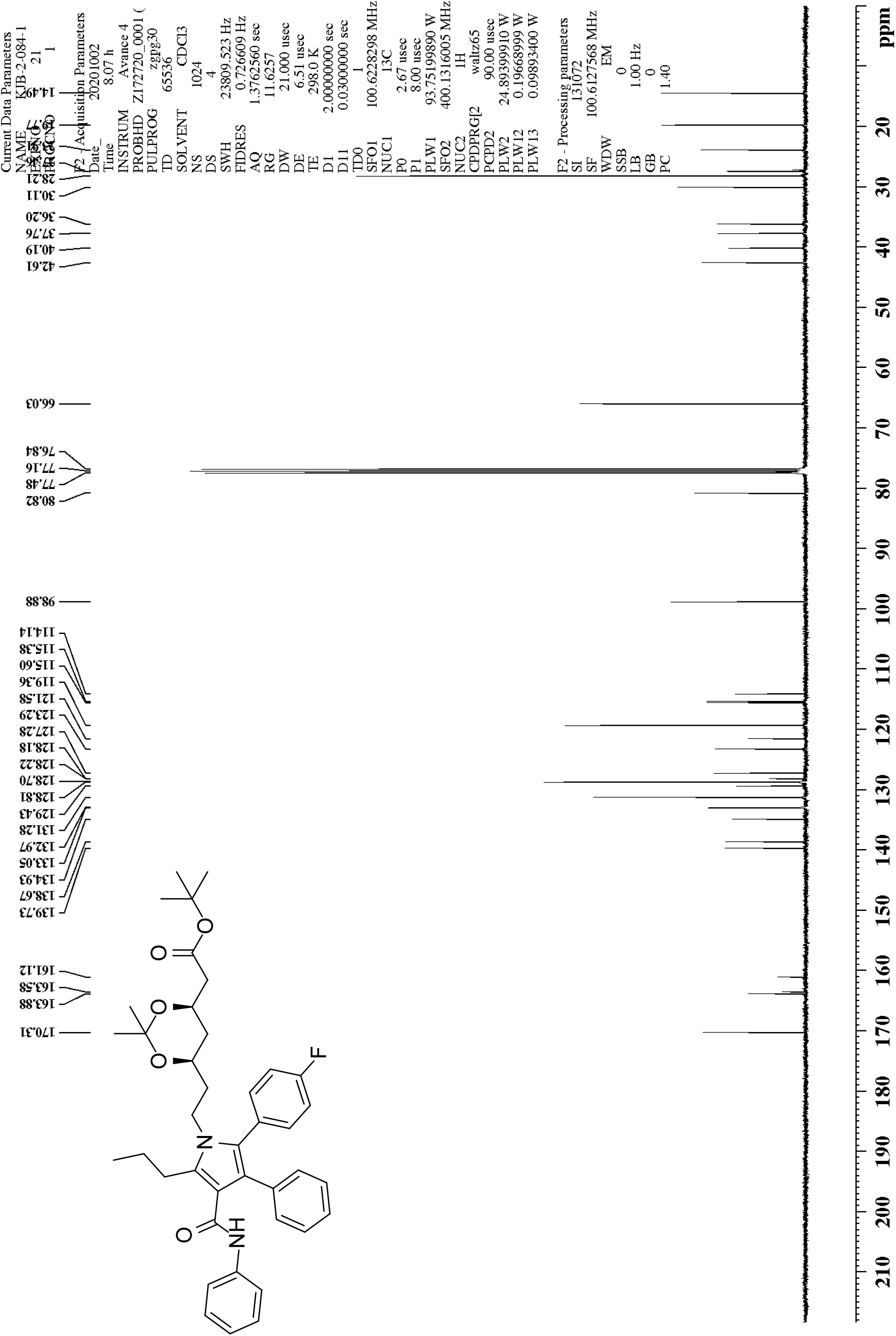
^13^C NMR of 1,1-dimethylethyl (4*R*,6*R*)-6-[2-[2-(4-fluorophenyl)-3-phenyl-4-[(phenylamino)carbonyl-5-propyl]-1*H*-pyrrol-1-yl]ethyl]-2,2-dimethyl-1,3-dioxane-4-acetate (21)

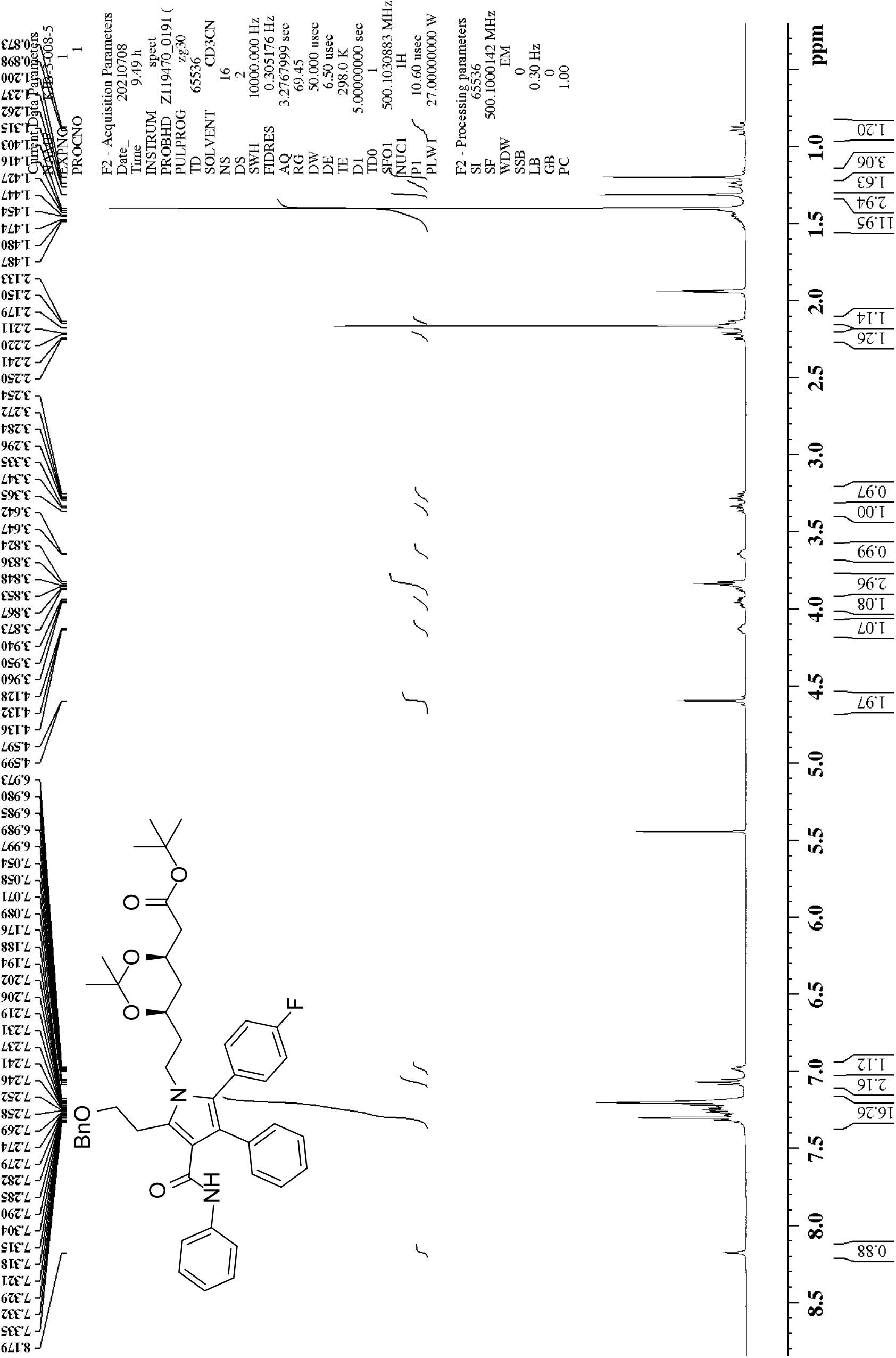
^1^H NMR of 1,1-dimethylethyl (4*R*,6*R*)-6-[5-(2-benzyloxyethyl)-2-[2-(4-fluorophenyl)-3-phenyl-4-[(phenylamino)carbonyl]-1*H*-pyrrol-1-yl]ethyl]-2,2-dimethyl-1,3-dioxane-4-acetate (22)

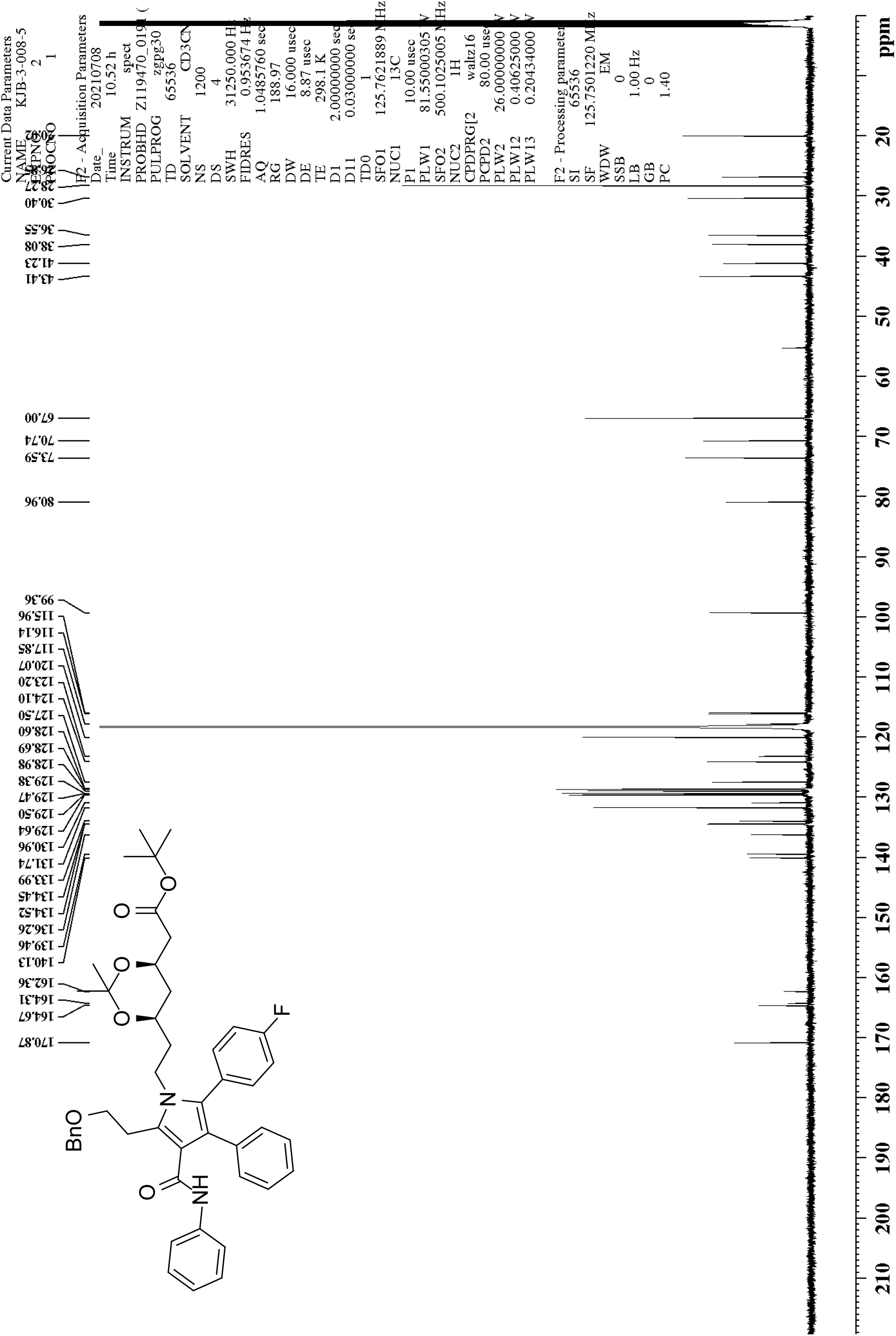
^13^C NMR of 1,1-dimethylethyl (4*R*,6*R*)-6-[5-(2-benzyloxyethyl)-2-[2-(4-fluorophenyl)-3-phenyl-4-[(phenylamino)carbonyl]-1*H*-pyrrol-1-yl]ethyl]-2,2-dimethyl-1,3-dioxane-4-acetate (22)

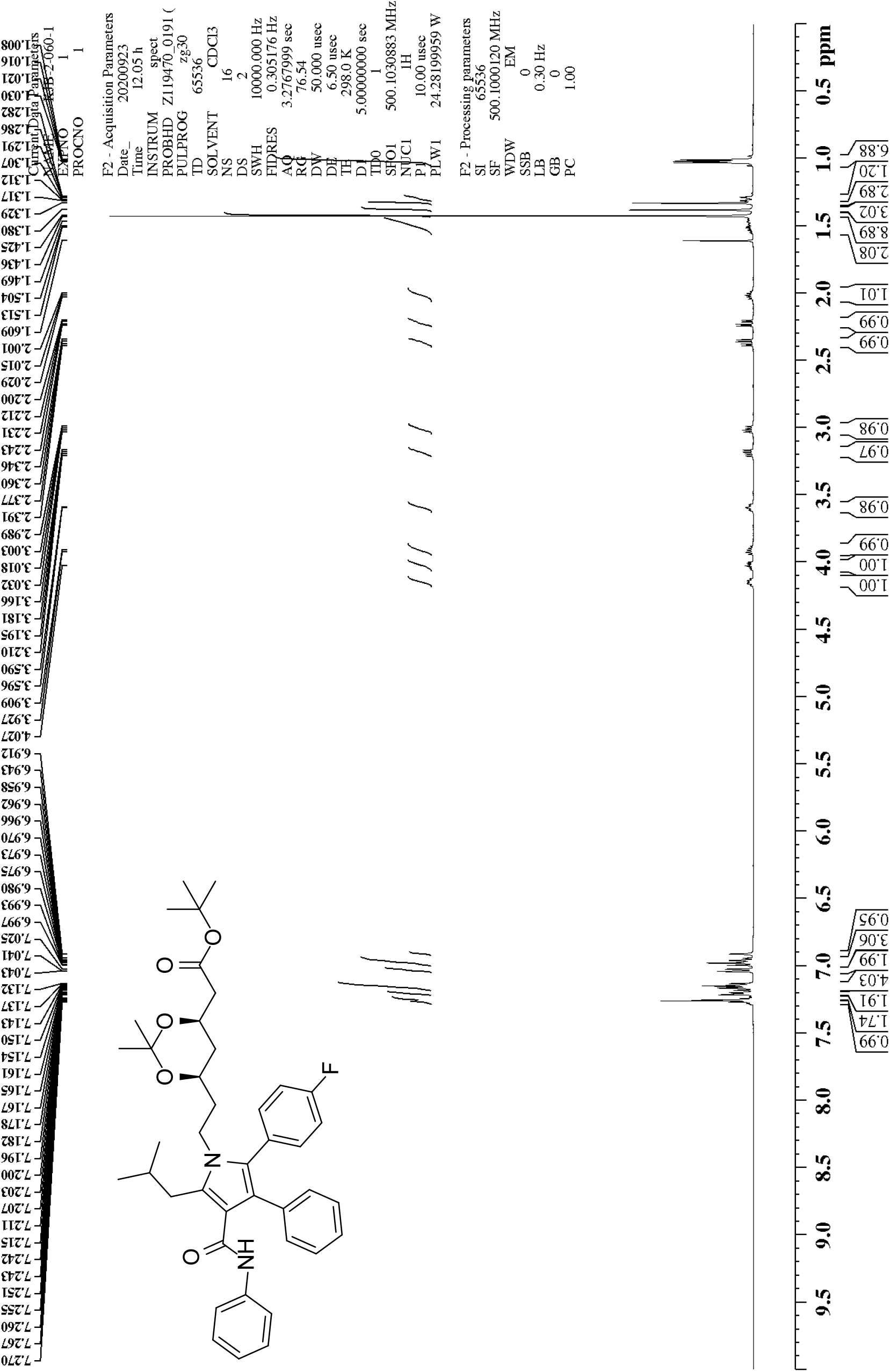
1H NMR of 1,1-dimethylethyl (4*R*,6*R*)-6-[2-[2-(4-fluorophenyl)-5-(2-methylpropyl)-3-phenyl-4-[(phenylamino)carbonyl]-1*H*-pyrrol-1-yl]ethyl]-2,2-dimethyl-1,3-dioxane-4-acetate (23)

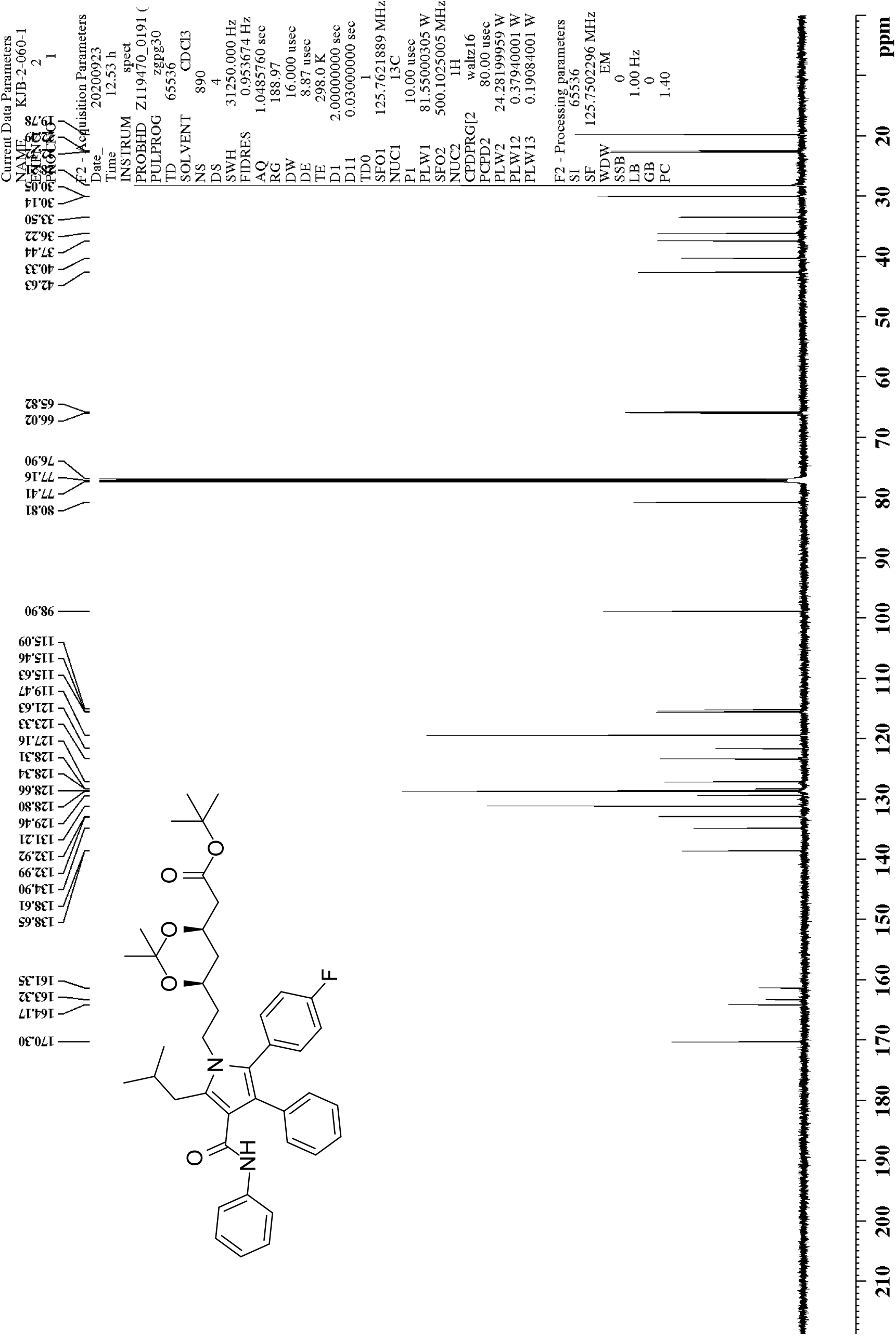
^13^C NMR of 1,1-dimethylethyl (4*R*,6*R*)-6-[2-[2-(4-fluorophenyl)-5-(2-methylpropyl)-3-phenyl-4-[(phenylamino)carbonyl]-1*H*-pyrrol-1-yl]ethyl]-2,2-dimethyl-1,3-dioxane-4-acetate (23)

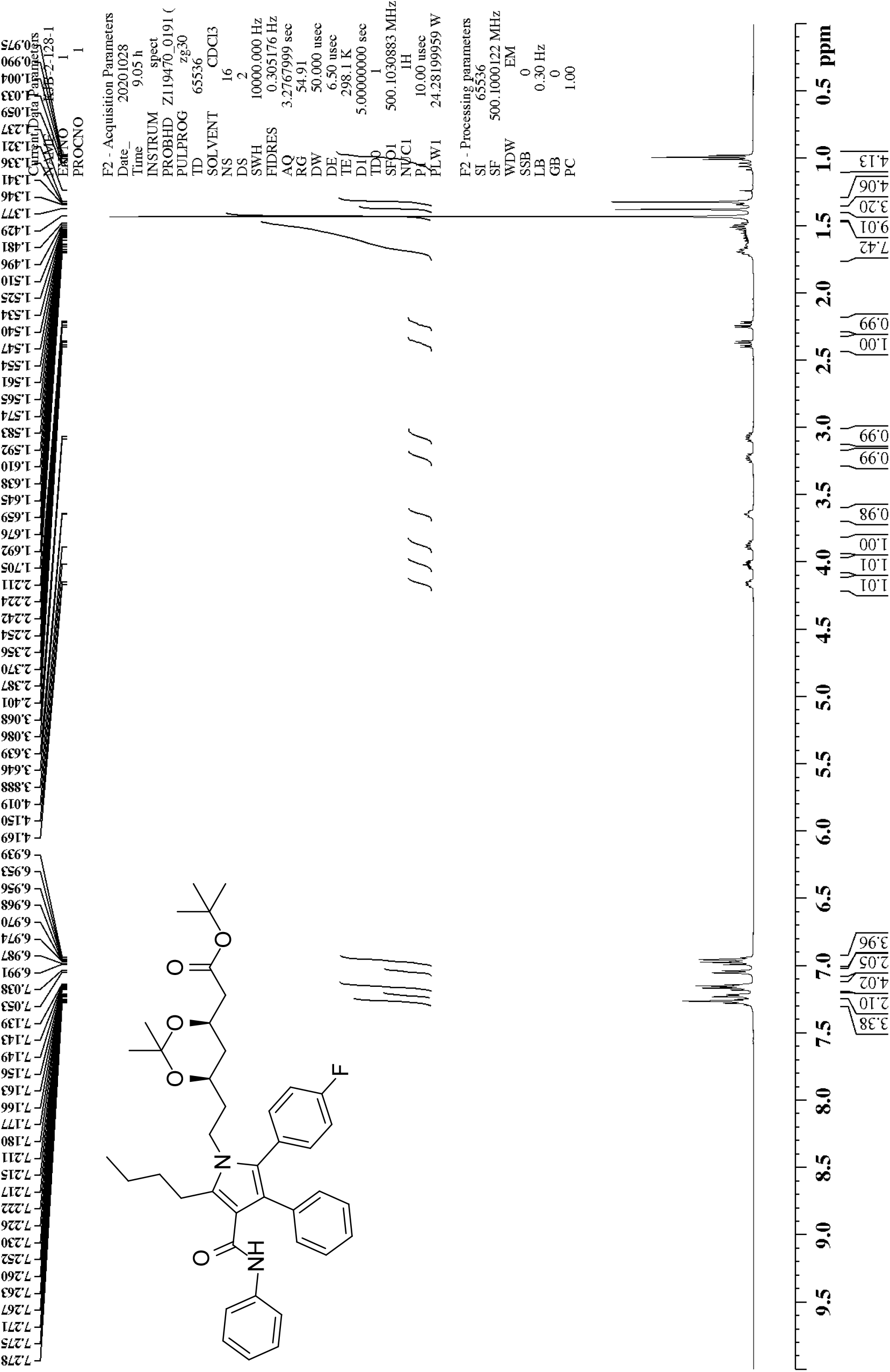
^1^H NMR of 1,1-dimethylethyl (4*R*,6*R*)-6-[2-[5-butyl-2-(4-fluorophenyl)-3-phenyl-4-[(phenylamino)carbonyl]-1*H*-pyrrol-1-yl]ethyl]-2,2-dimethyl-1,3-dioxane-4-acetate (24)

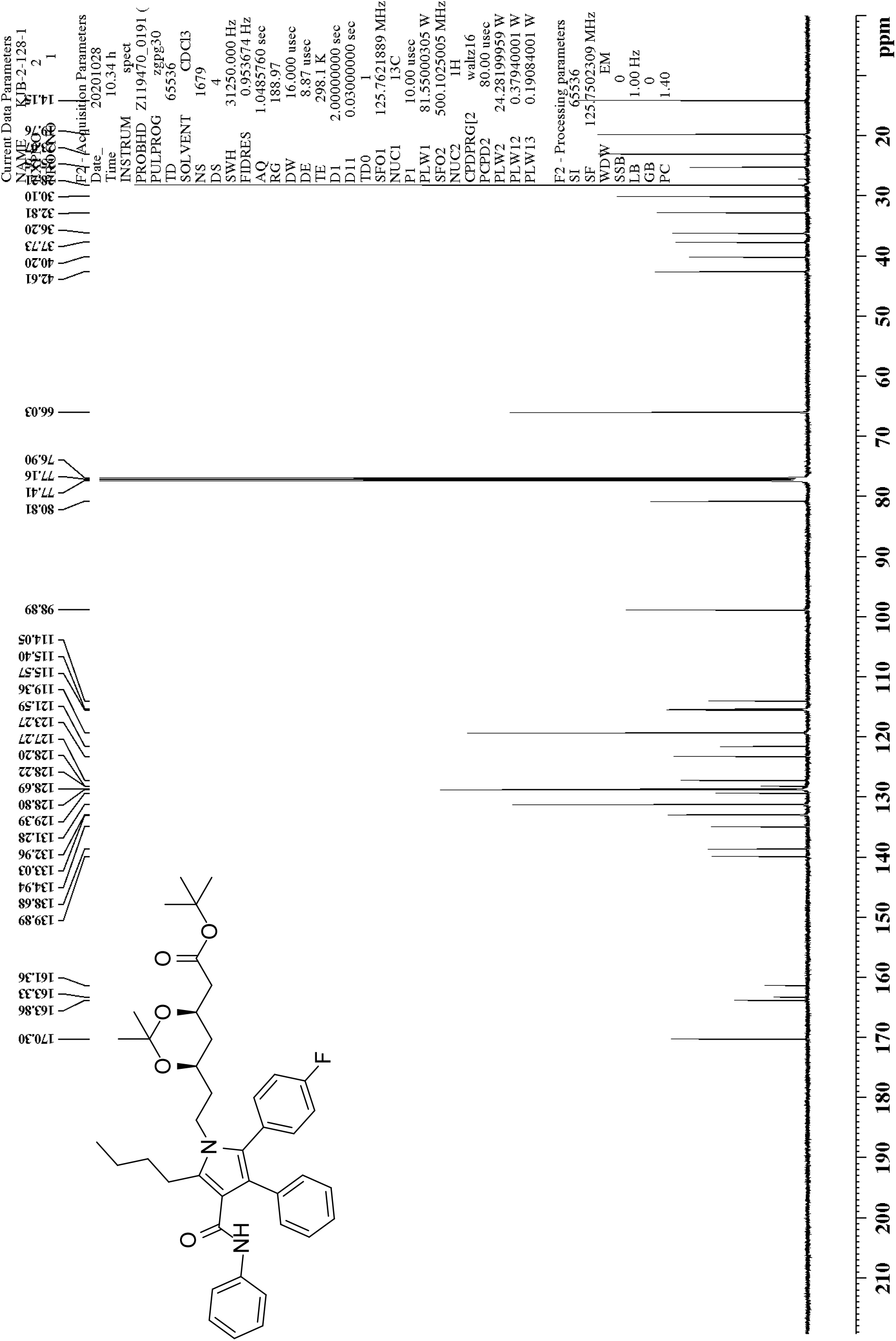
^13^C NMR of 1,1-dimethylethyl (4*R*,6*R*)-6-[2-[5-butyl-2-(4-fluorophenyl)-3-phenyl-4-[(phenylamino)carbonyl]-1*H*-pyrrol-1-yl]ethyl]-2,2-dimethyl-1,3-dioxane-4-acetate (24)

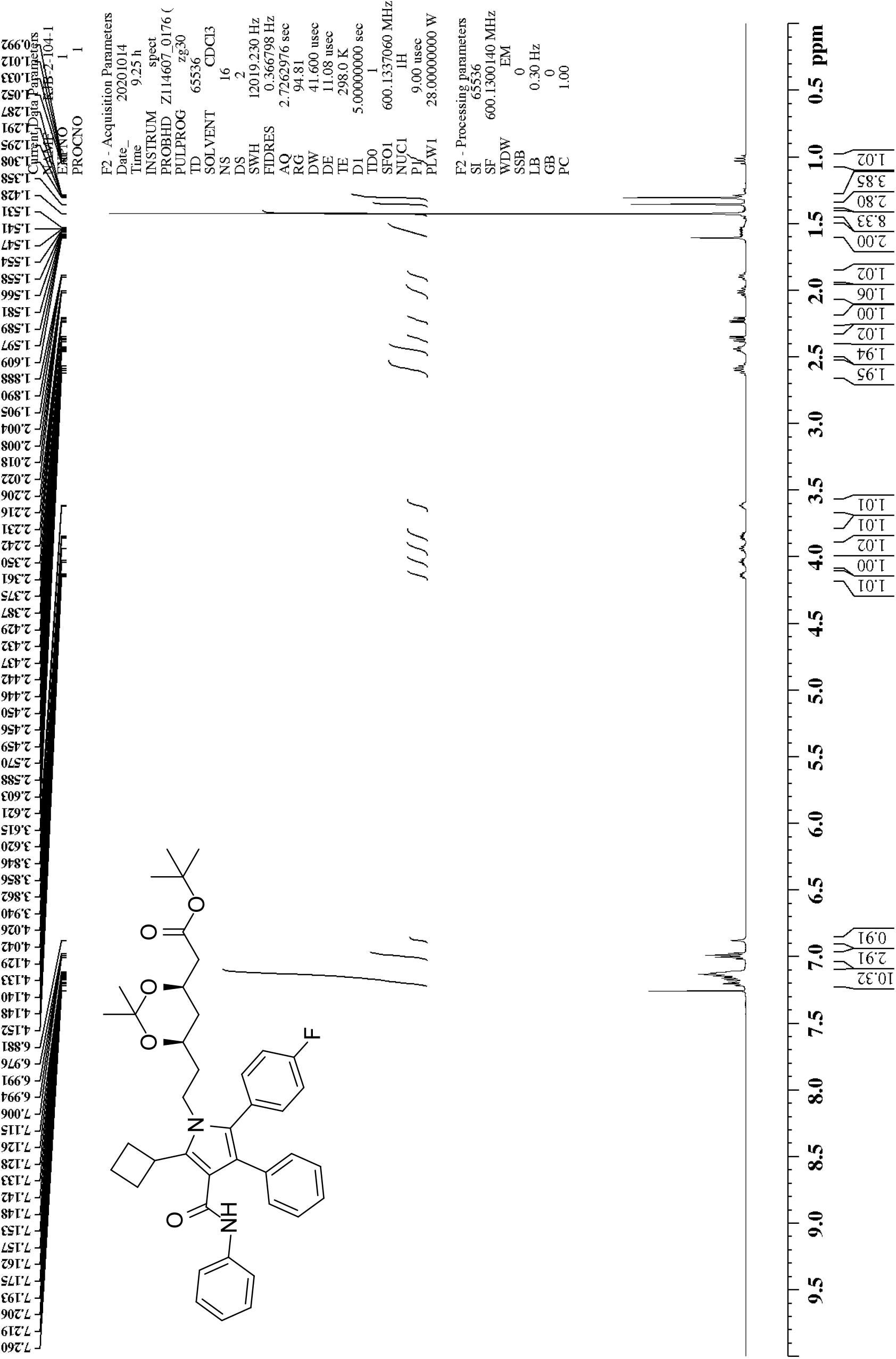
^1^H NMR of 1,1-dimethylethyl (4*R*,6*R*)-6-[2-[5-cyclobutyl-2-(4-fluorophenyl)-3-phenyl-4-[(phenylamino)carbonyl]-1*H*-pyrrol-1-yl]ethyl]-2,2-dimethyl-1,3-dioxane-4-acetate (25)

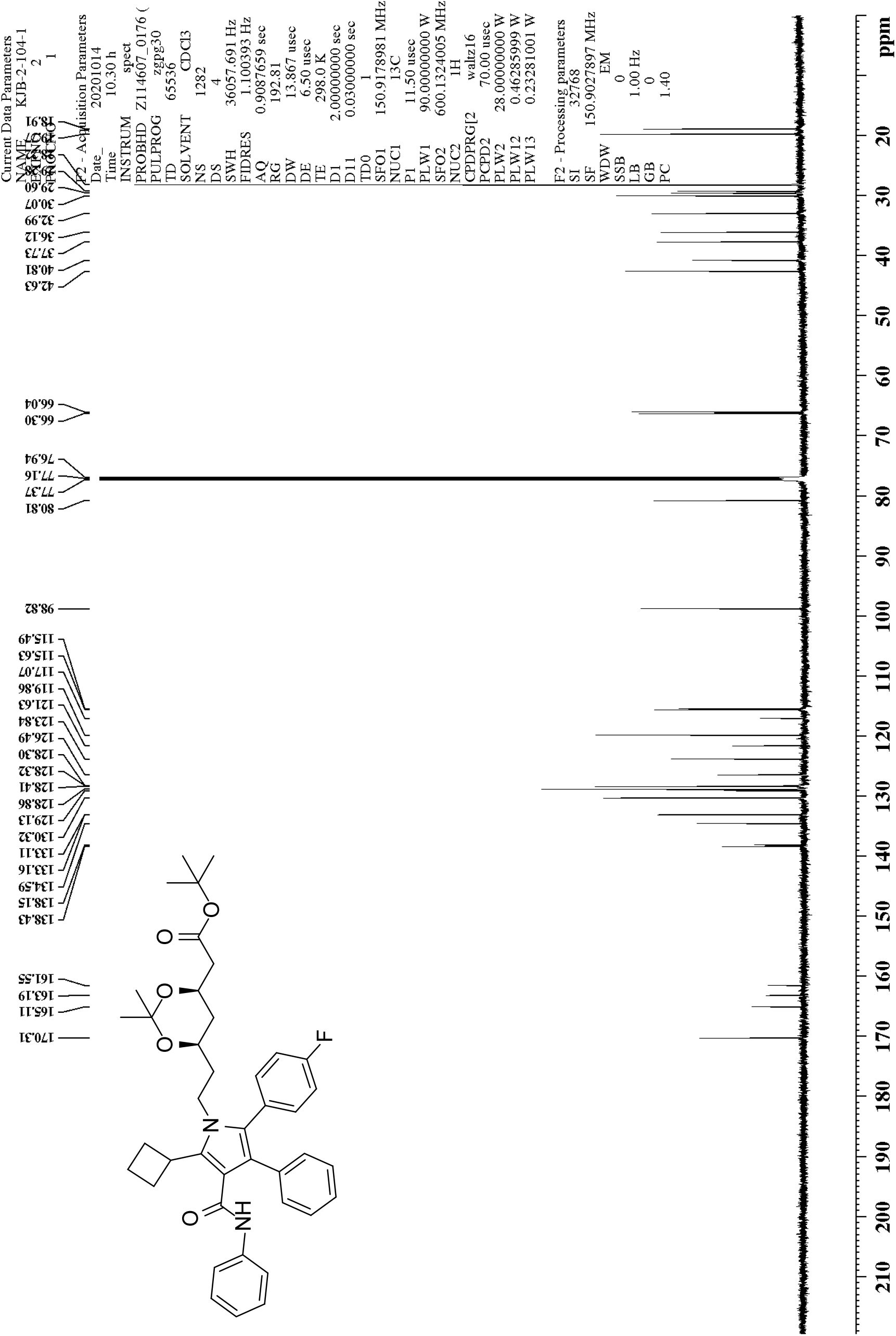
^13^C NMR of 1,1-dimethylethyl (4*R*,6*R*)-6-[2-[5-cyclobutyl-2-(4-fluorophenyl)-3-phenyl-4-[(phenylamino)carbonyl]-1*H*-pyrrol-1-yl]ethyl]-2,2-dimethyl-1,3-dioxane-4-acetate (25)

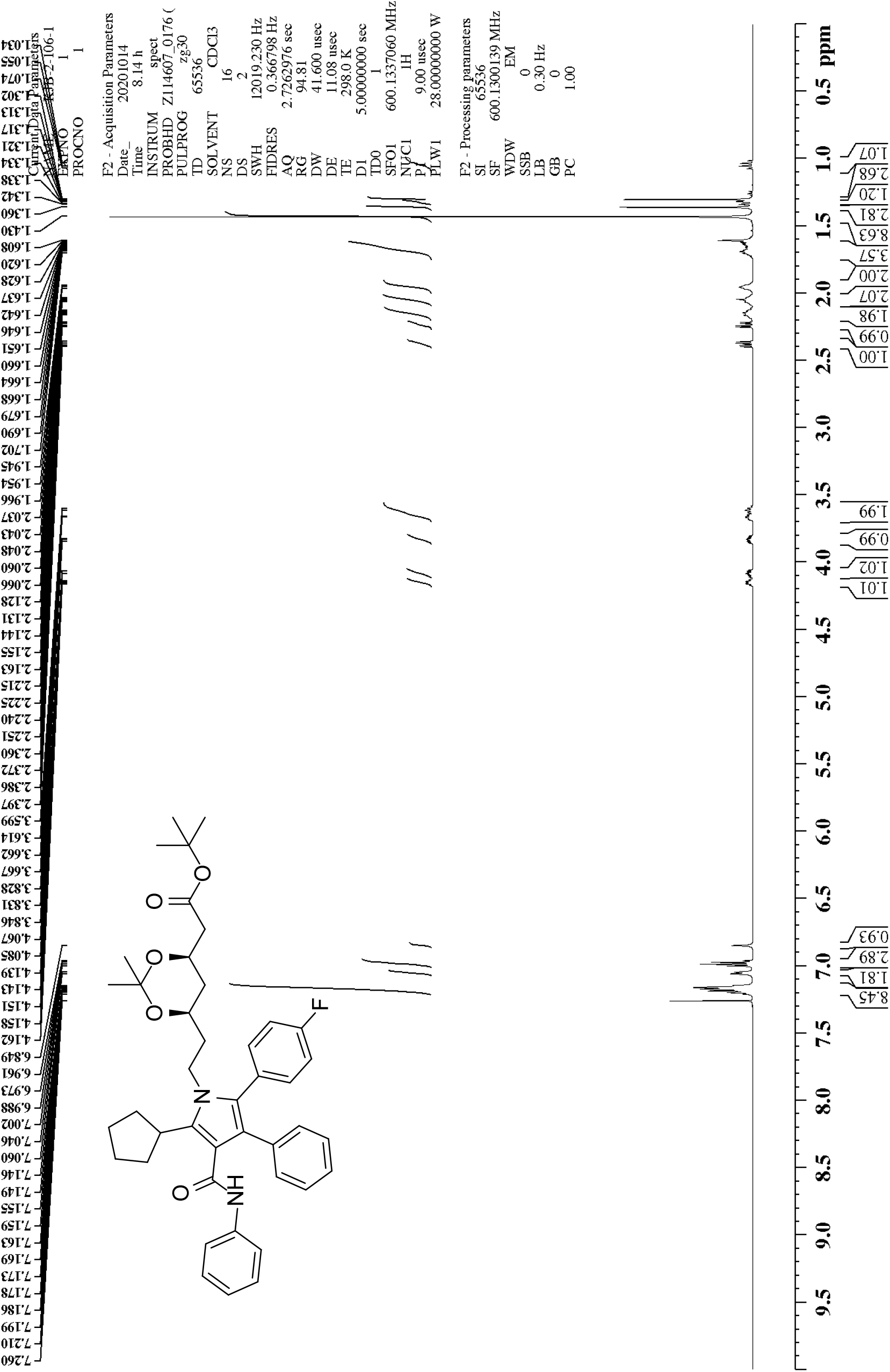
^1^H NMR of 1,1-dimethylethyl (4*R*,6*R*)-6-[2-[5-cyclopentyl-2-(4-fluorophenyl)-3-phenyl-4-[(phenylamino)carbonyl]-1*H*-pyrrol-1-yl]ethyl]-2,2-dimethyl-1,3-dioxane-4-acetate (26)

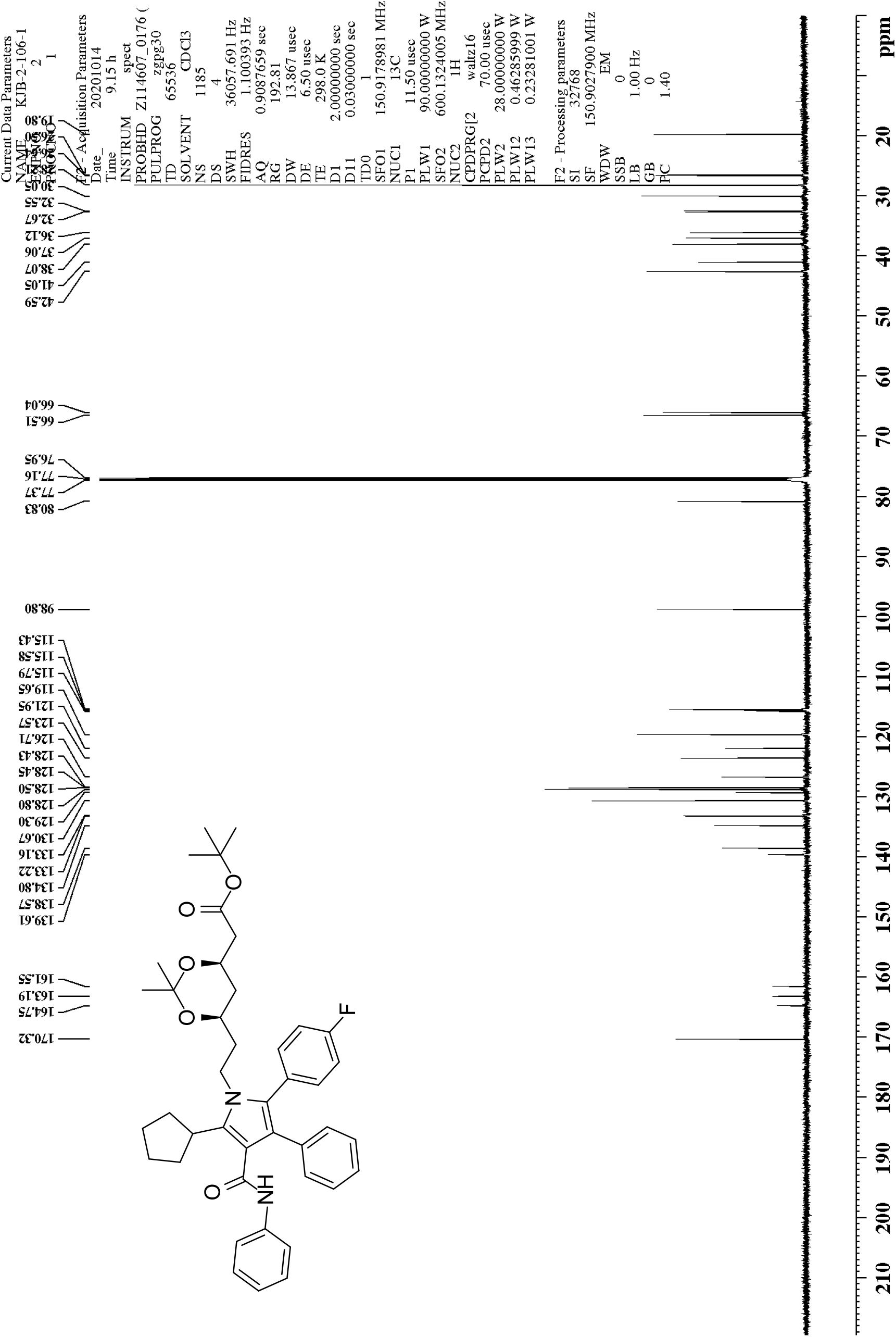
^13^C NMR of 1,1-dimethylethyl (4*R*,6*R*)-6-[2-[5-cyclopentyl-2-(4-fluorophenyl)-3-phenyl-4-[(phenylamino)carbonyl]-1*H*-pyrrol-1-yl]ethyl]-2,2-dimethyl-1,3-dioxane-4-acetate (26)

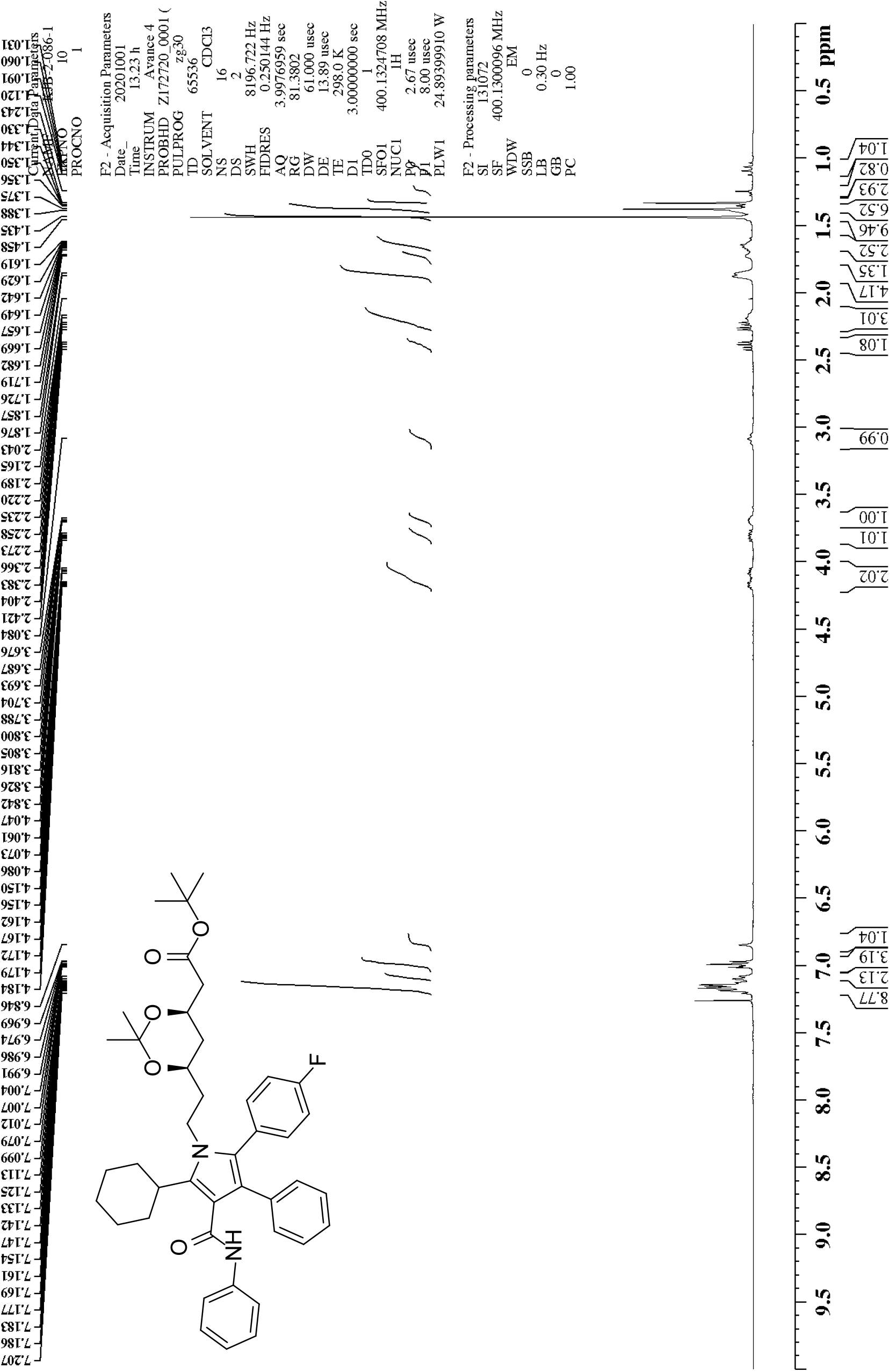
^1^H NMR of 1,1-dimethylethyl (4*R*,6*R*)-6-[2-[5-cyclohexyl-2-(4-fluorophenyl)-3-phenyl-4-[(phenylamino)carbonyl]-1*H*-pyrrol-1-yl]ethyl]-2,2-dimethyl-1,3-dioxane-4-acetate (27)

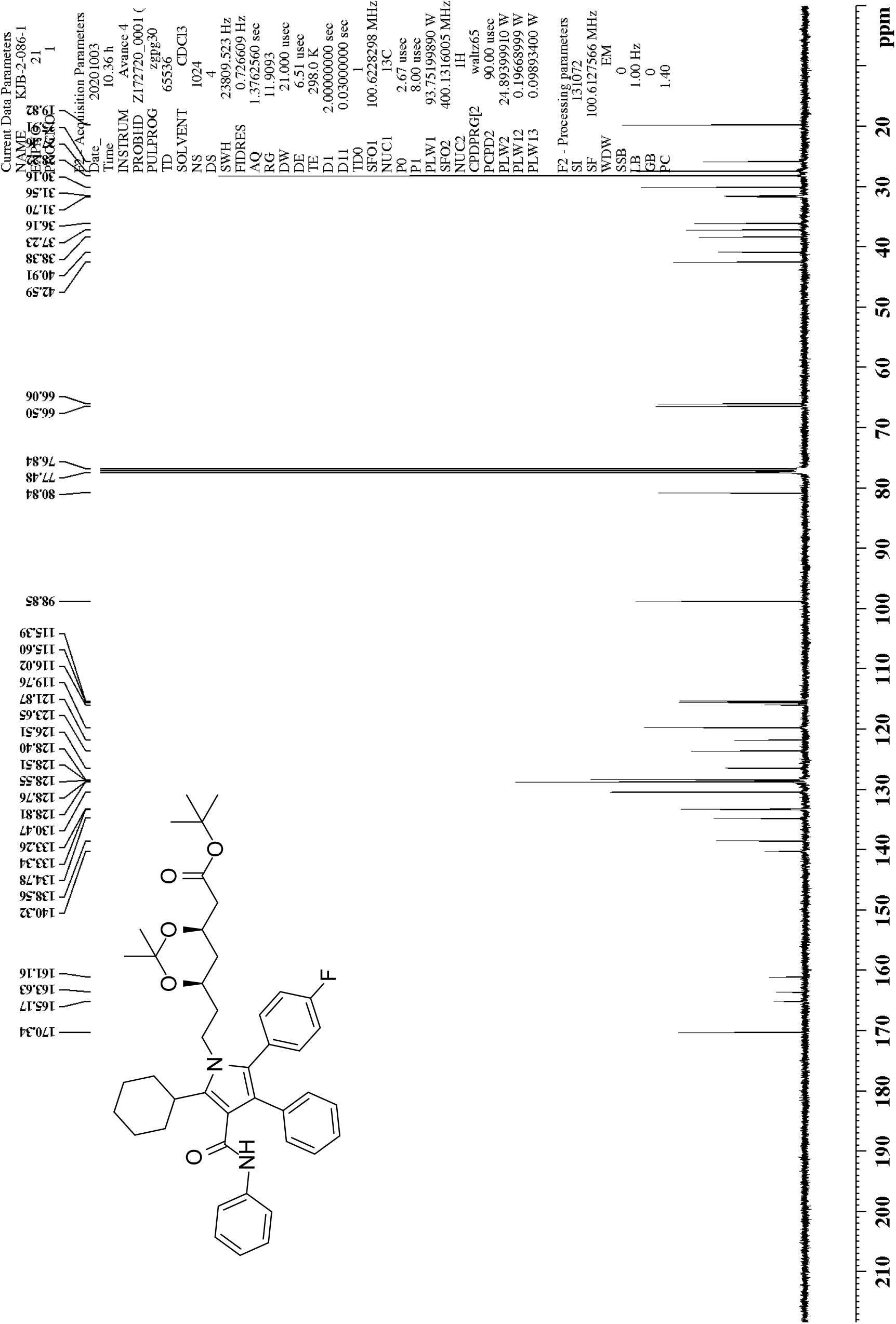
^13^C NMR of 1,1-dimethylethyl (4*R*,6*R*)-6-[2-[5-cyclohexyl-2-(4-fluorophenyl)-3-phenyl-4-[(phenylamino)carbonyl]-1*H*-pyrrol-1-yl]ethyl]-2,2-dimethyl-1,3-dioxane-4-acetate (27)

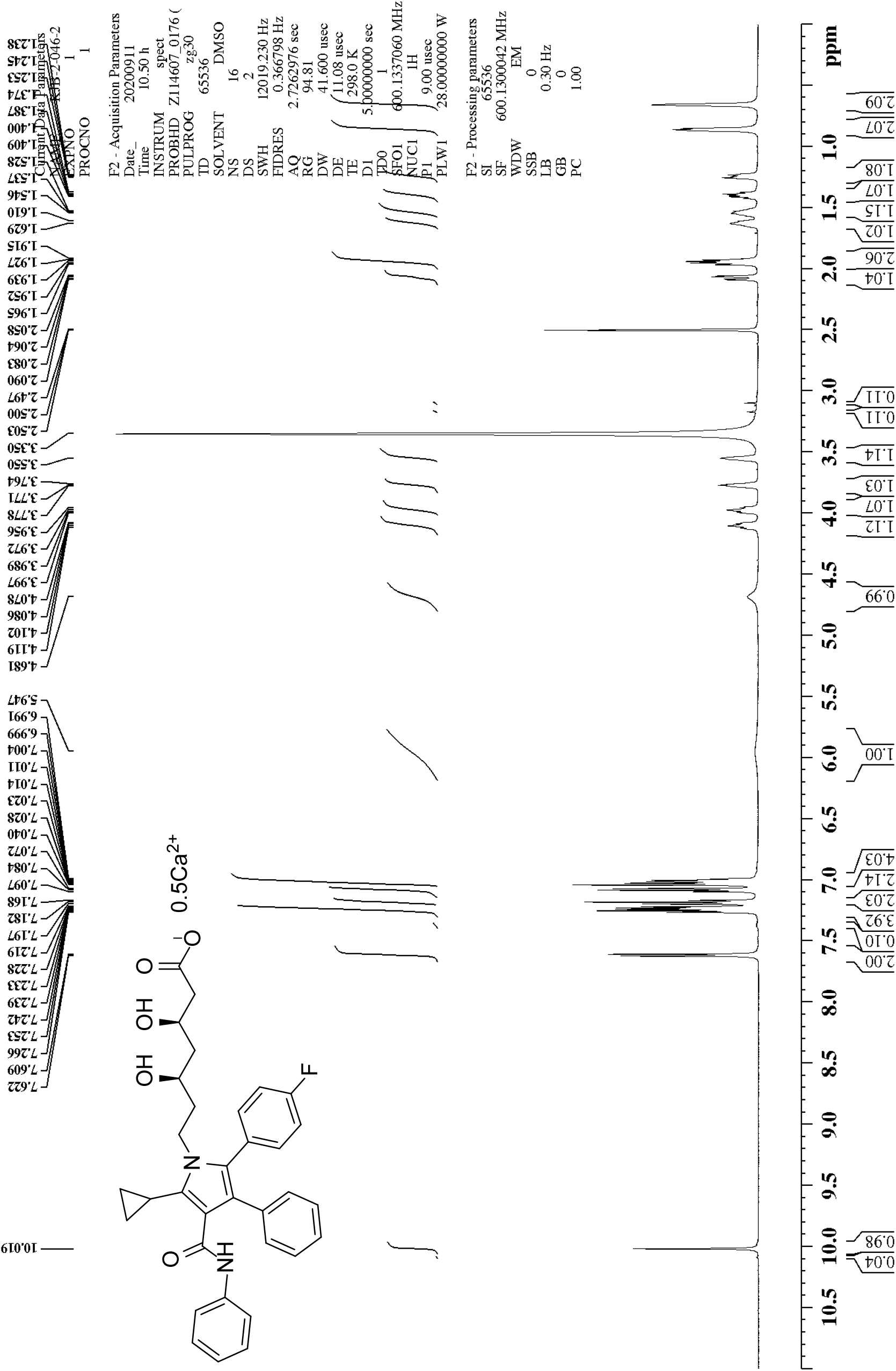
^1^H NMR of (β*R*,δ*R*)-5-cyclopropyl-2-(4-fluorophenyl)-β,δ-dihydroxy-3-phenyl-4-[(phenylamino)carbonyl]-1*H*-pyrrole-1-heptanoic acid hemicalcium salt (1)

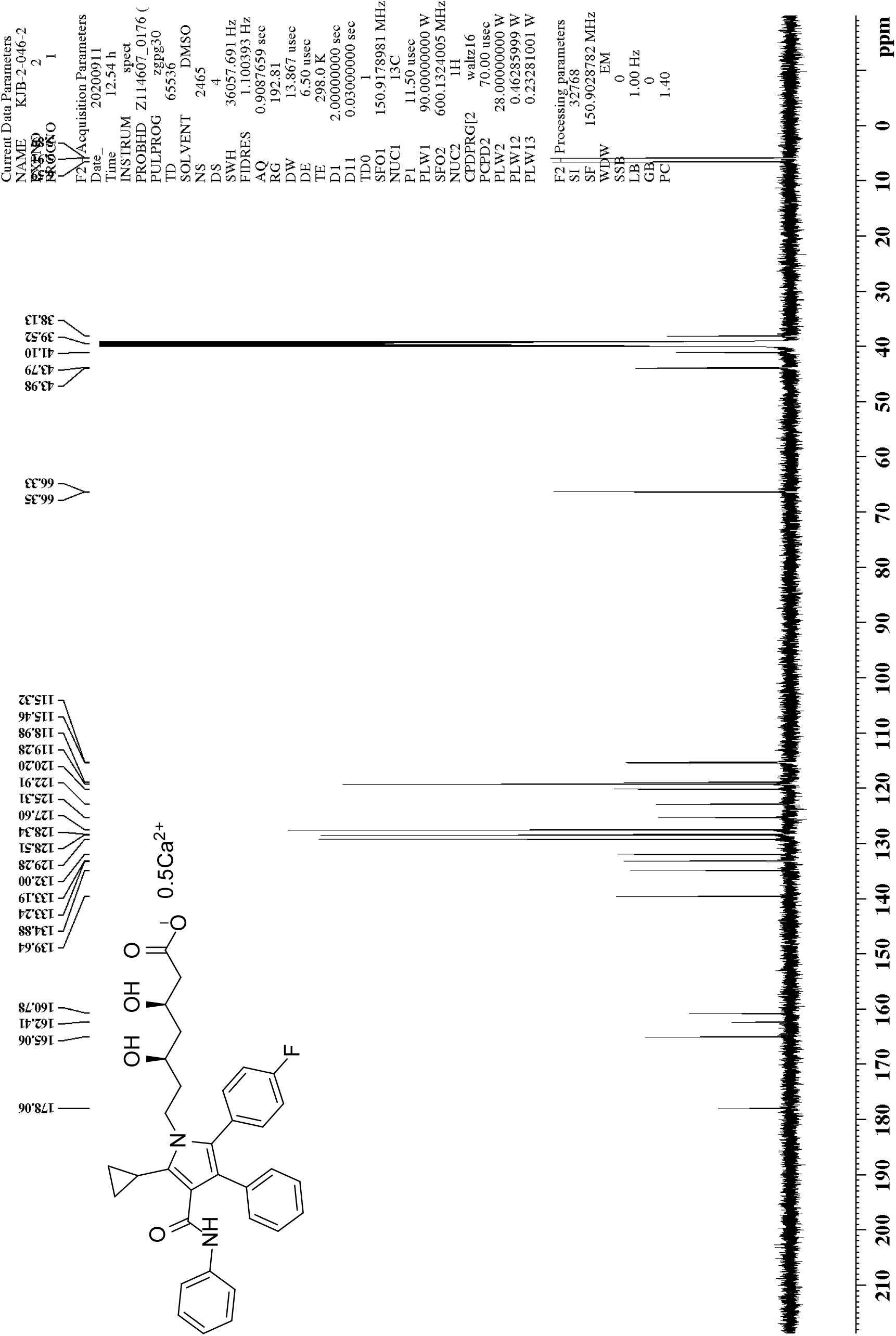
^13^C NMR of (β*R*,δ*R*)-5-cyclopropyl-2-(4-fluorophenyl)-β,δ-dihydroxy-3-phenyl-4-[(phenylamino)carbonyl]-1*H*-pyrrole-1-heptanoic acid hemicalcium salt (1)

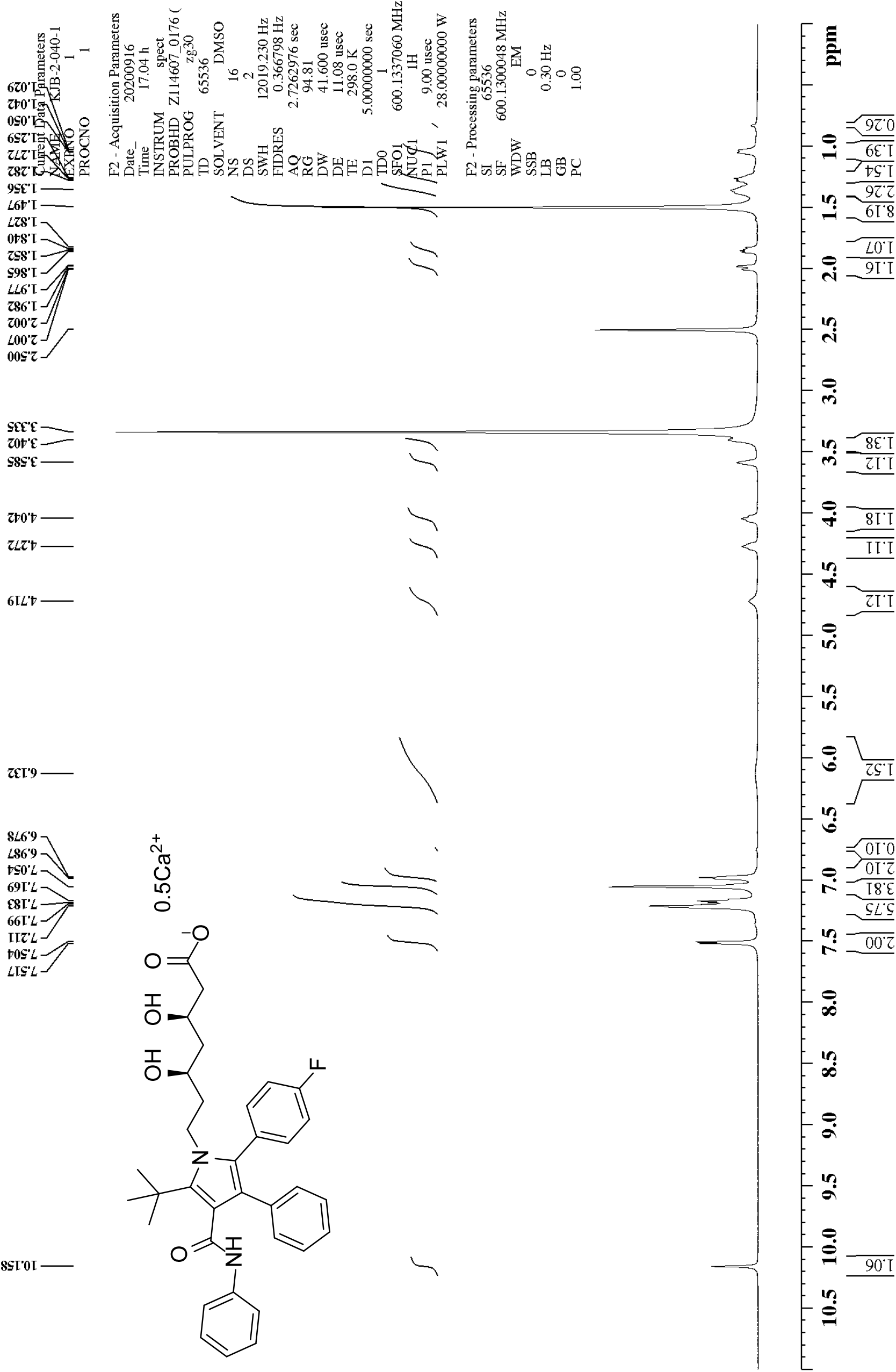
^1^H NMR of (β*R*,δ*R*)-2-(4-fluorophenyl)-β,δ-dihydroxy-5-(1,1-dimethylethyl)-3-phenyl-4-[(phenylamino)carbonyl]-1*H*-pyrrole-1-heptanoic acid hemicalcium salt (2)

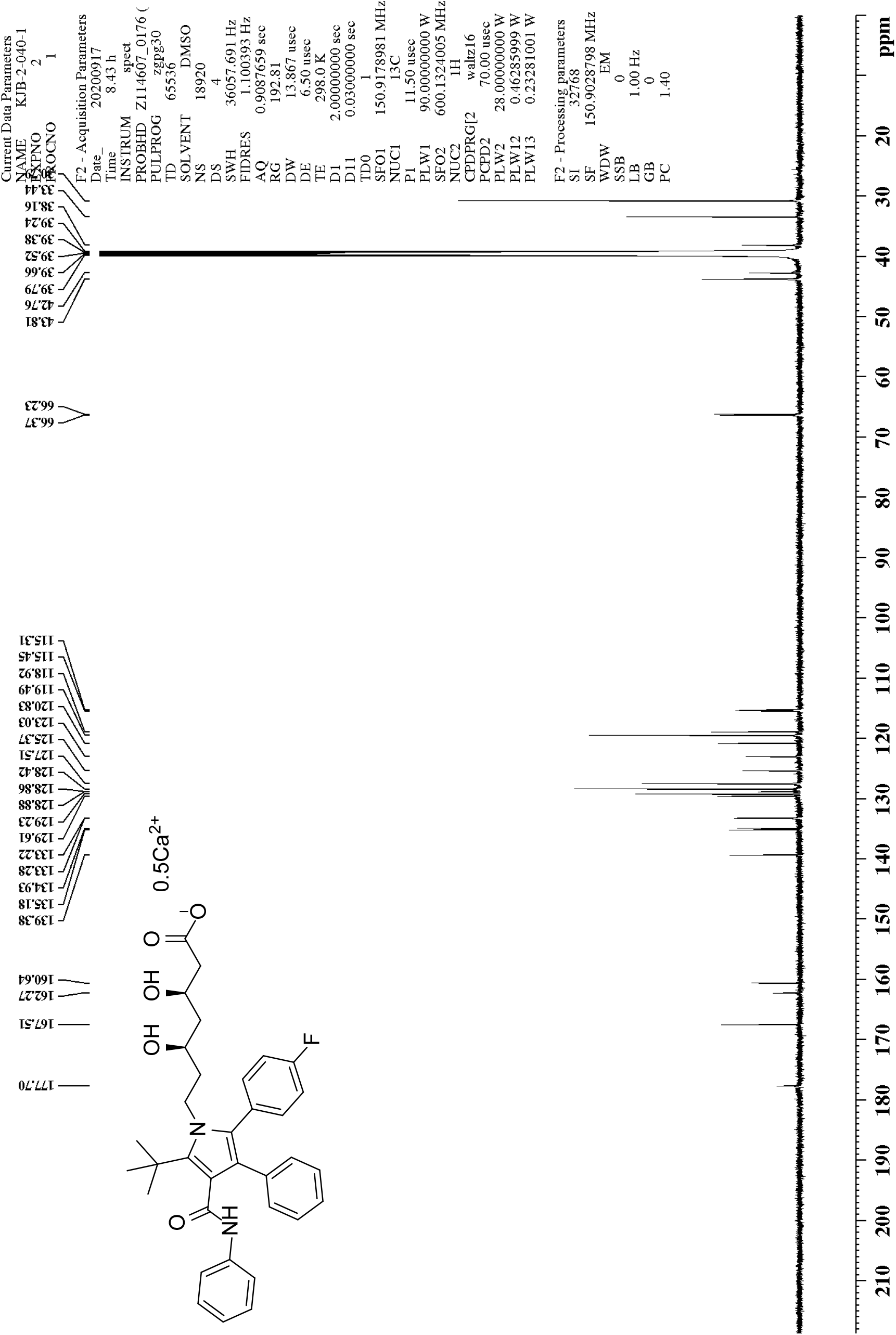
^13^C NMR of (β*R*,δ*R*)-2-(4-fluorophenyl)-β,δ-dihydroxy-5-(1,1-dimethylethyl)-3-phenyl-4-[(phenylamino)carbonyl]-1*H*-pyrrole-1-heptanoic acid hemicalcium salt (2)

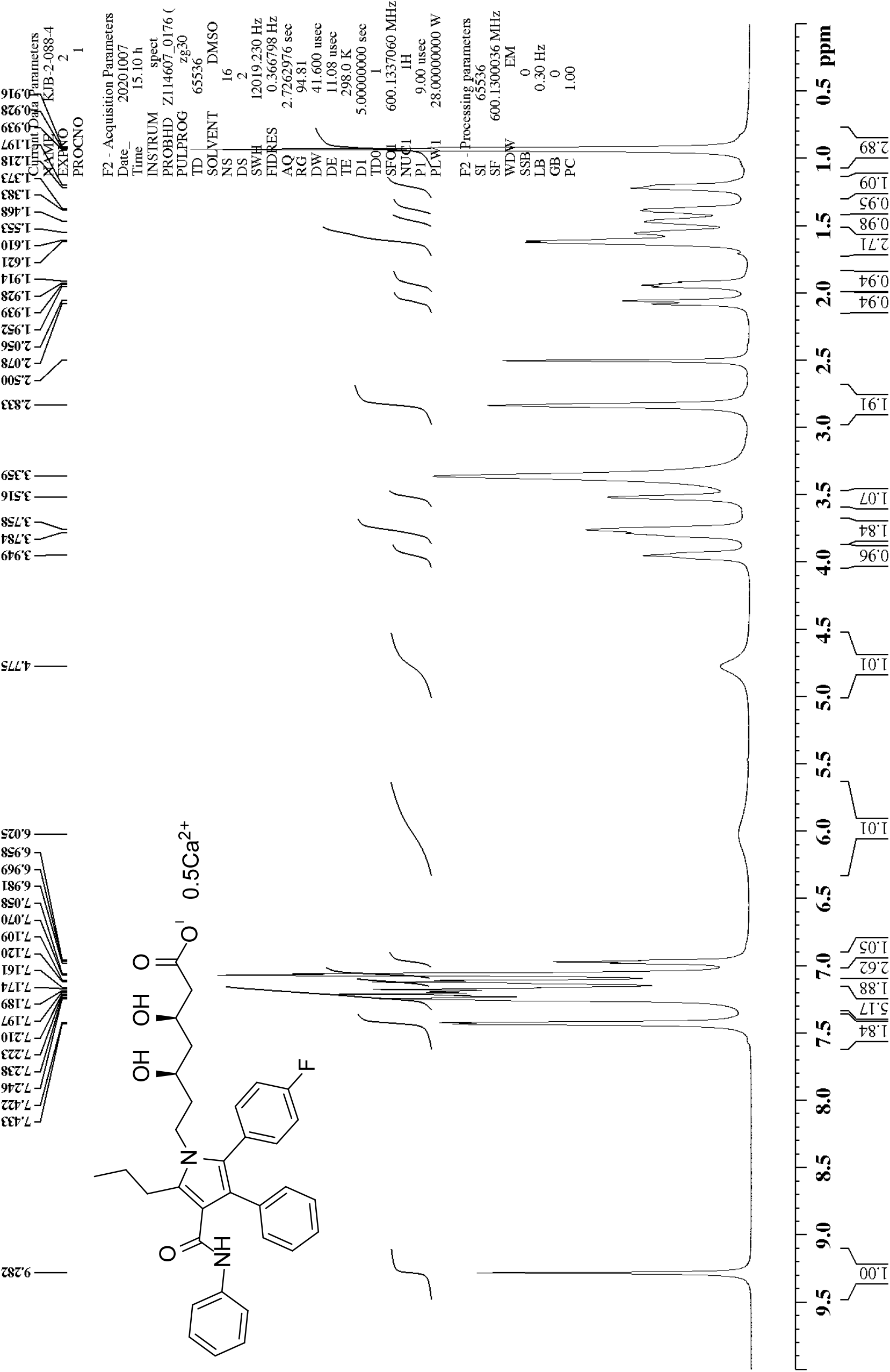
^1^H NMR of (β*R*,δ*R*)-2-(4-fluorophenyl)-β,δ-dihydroxy-3-phenyl-4-[(phenylamino)carbonyl-5-propyl]-1*H*-pyrrole-1-heptanoic acid hemicalcium salt (3)

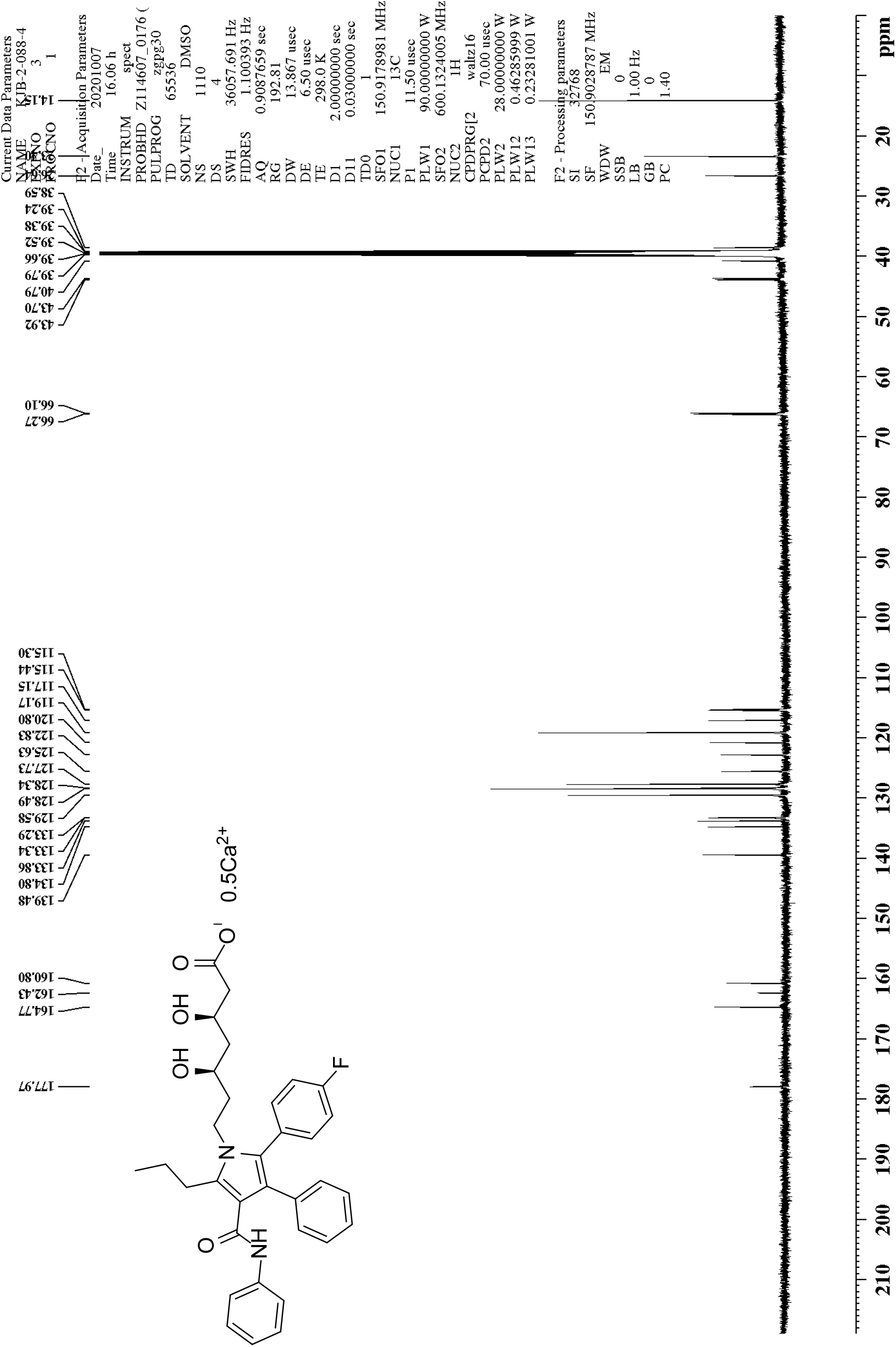
^13^C NMR of (β*R*,δ*R*)-2-(4-fluorophenyl)-β,δ-dihydroxy-3-phenyl-4-[(phenylamino)carbonyl-5-propyl]-1*H*-pyrrole-1-heptanoic acid hemicalcium salt (3)

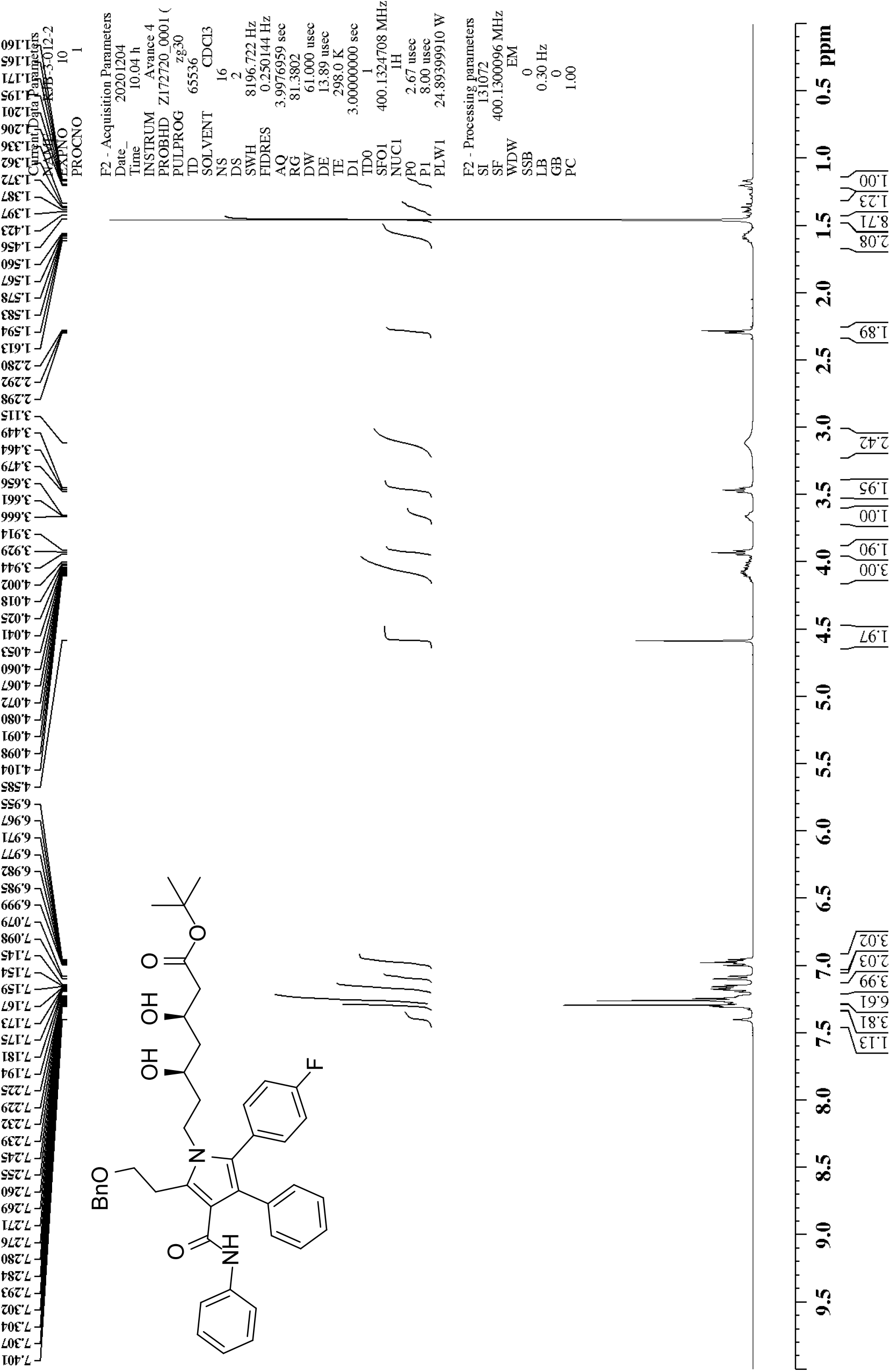
^1^H NMR of 1,1-dimethylethyl (3*R*,5*R*)-7-[5-(2-benzyloxyethyl)-2-(4-fluorophenyl)-3-phenyl-4-phenylcarbamoylpyrrol-1-yl]-3,5-dihydroxyheptanoate.

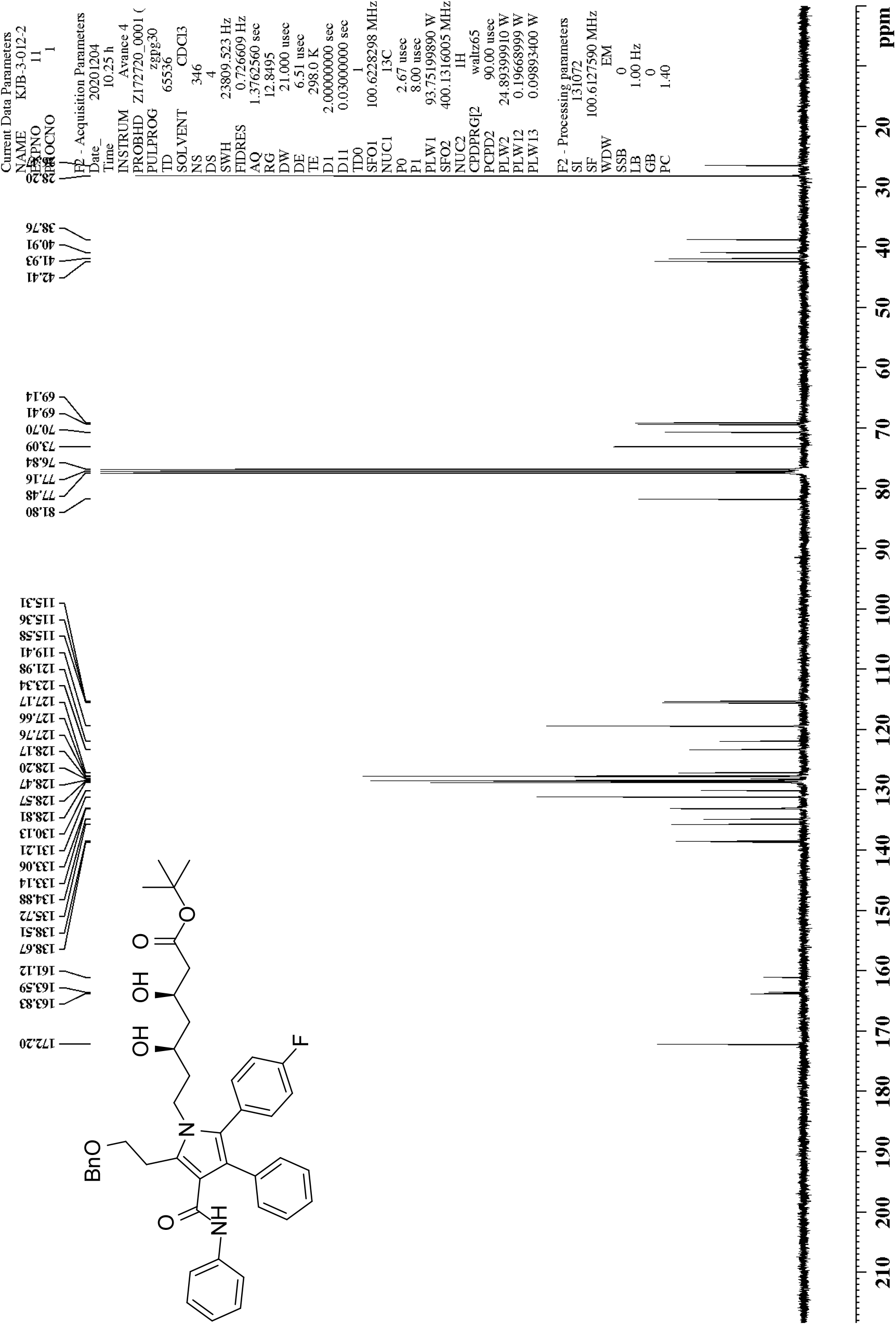
^13^C NMR of 1,1-dimethylethyl (3*R*,5*R*)-7-[5-(2-benzyloxyethyl)-2-(4-fluorophenyl)-3-phenyl-4-phenylcarbamoylpyrrol-1-yl]-3,5-dihydroxyheptanoate.

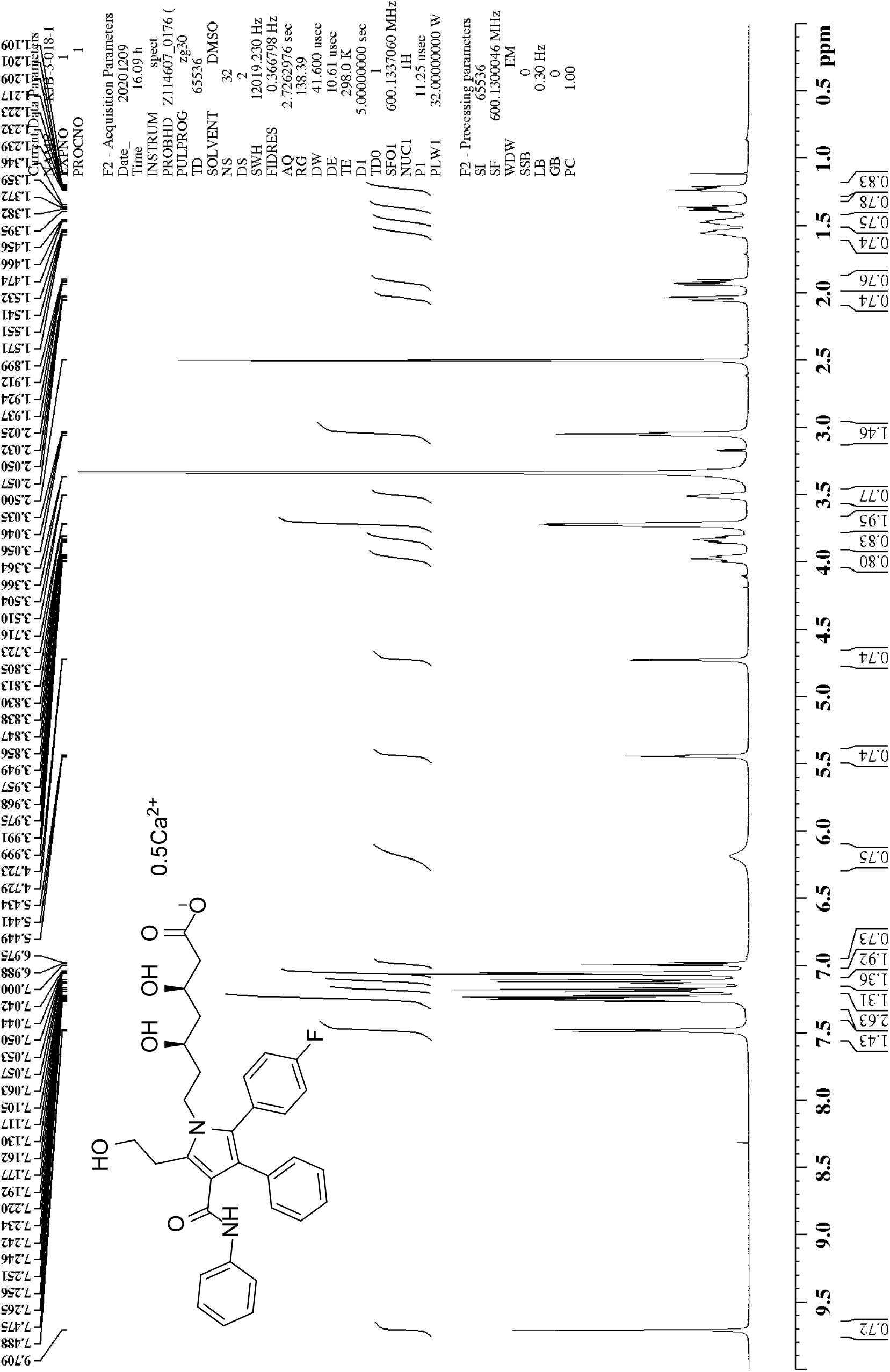
^1^H NMR of (β*R*,δ*R*)-2-(4-fluorophenyl)-β,δ-dihydroxy-5-(2-hydroxyethyl)-3-phenyl-4-[(phenylamino)carbonyl]-1*H*-pyrrole-1-heptanoic acid hemicalcium salt (4)

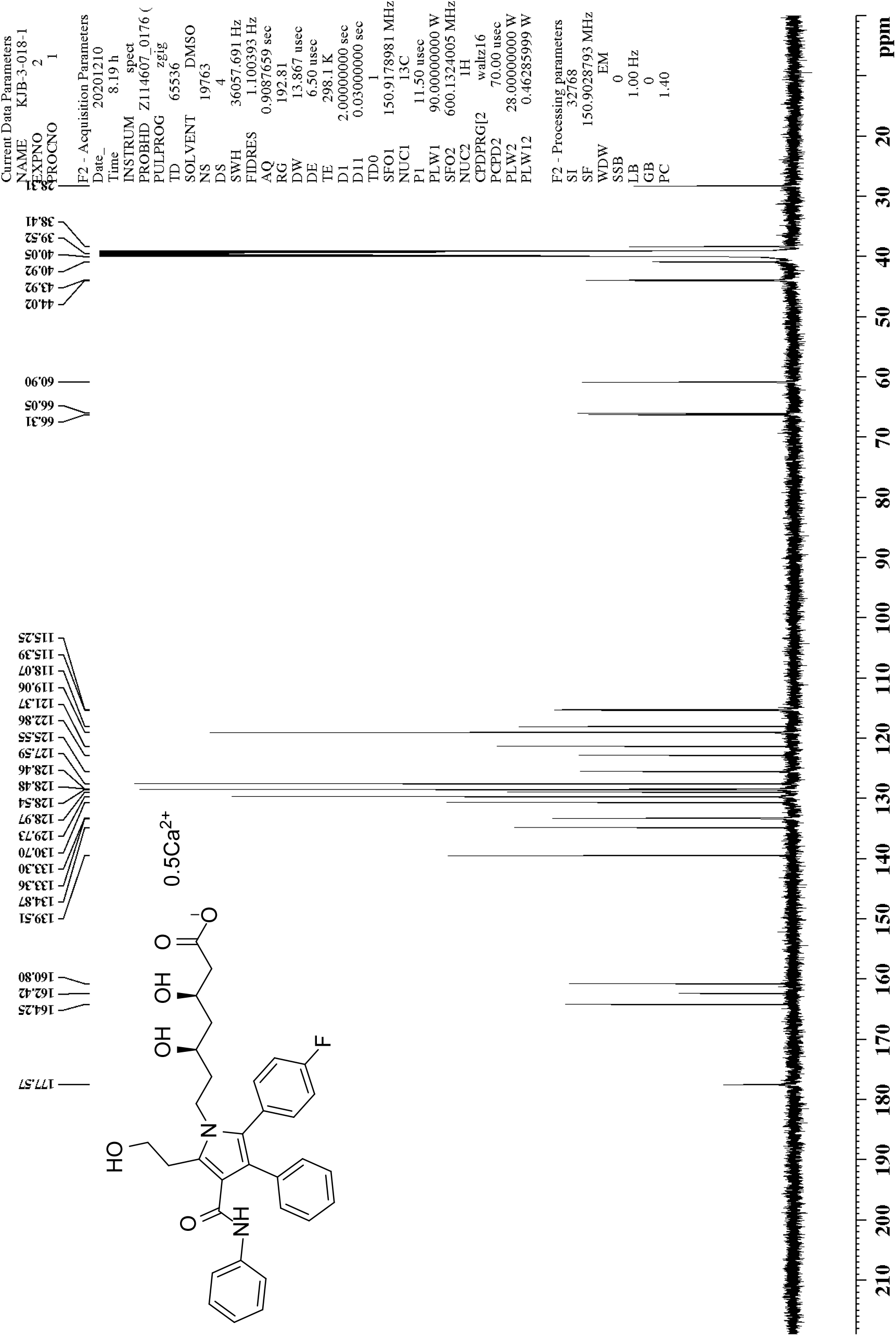
^13^C NMR of (β*R*,δ*R*)-2-(4-fluorophenyl)-β,δ-dihydroxy-5-(2-hydroxyethyl)-3-phenyl-4-[(phenylamino)carbonyl]-1*H*-pyrrole-1-heptanoic acid hemicalcium salt (4)

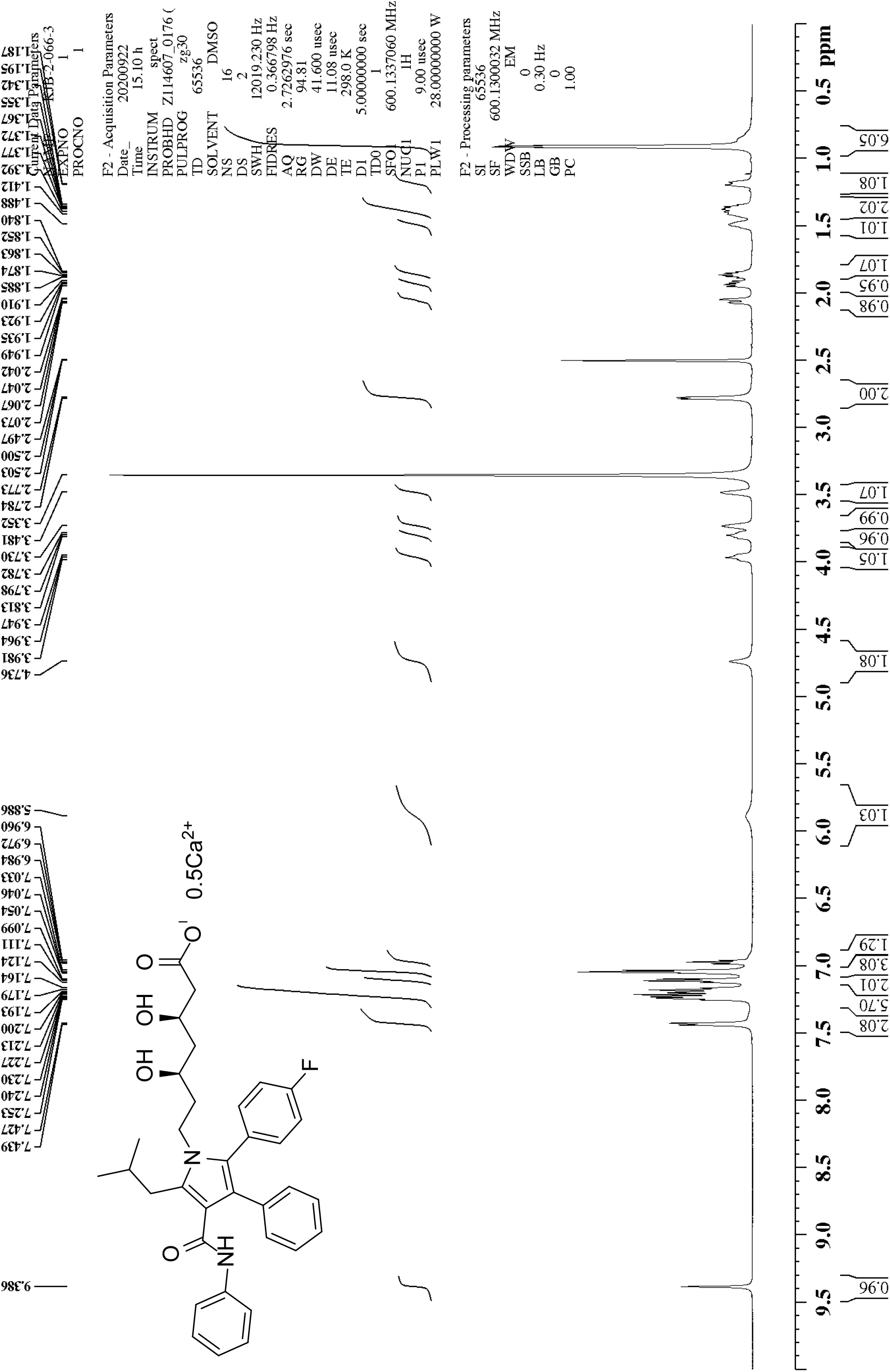
^1^H NMR of (β*R*,δ*R*)-2-(4-fluorophenyl)-β,δ-dihydroxy-5-(2-methylpropyl)-3-phenyl-4-[(phenylamino)carbonyl]-1*H*-pyrrole-1-heptanoic acid hemicalcium salt (5)

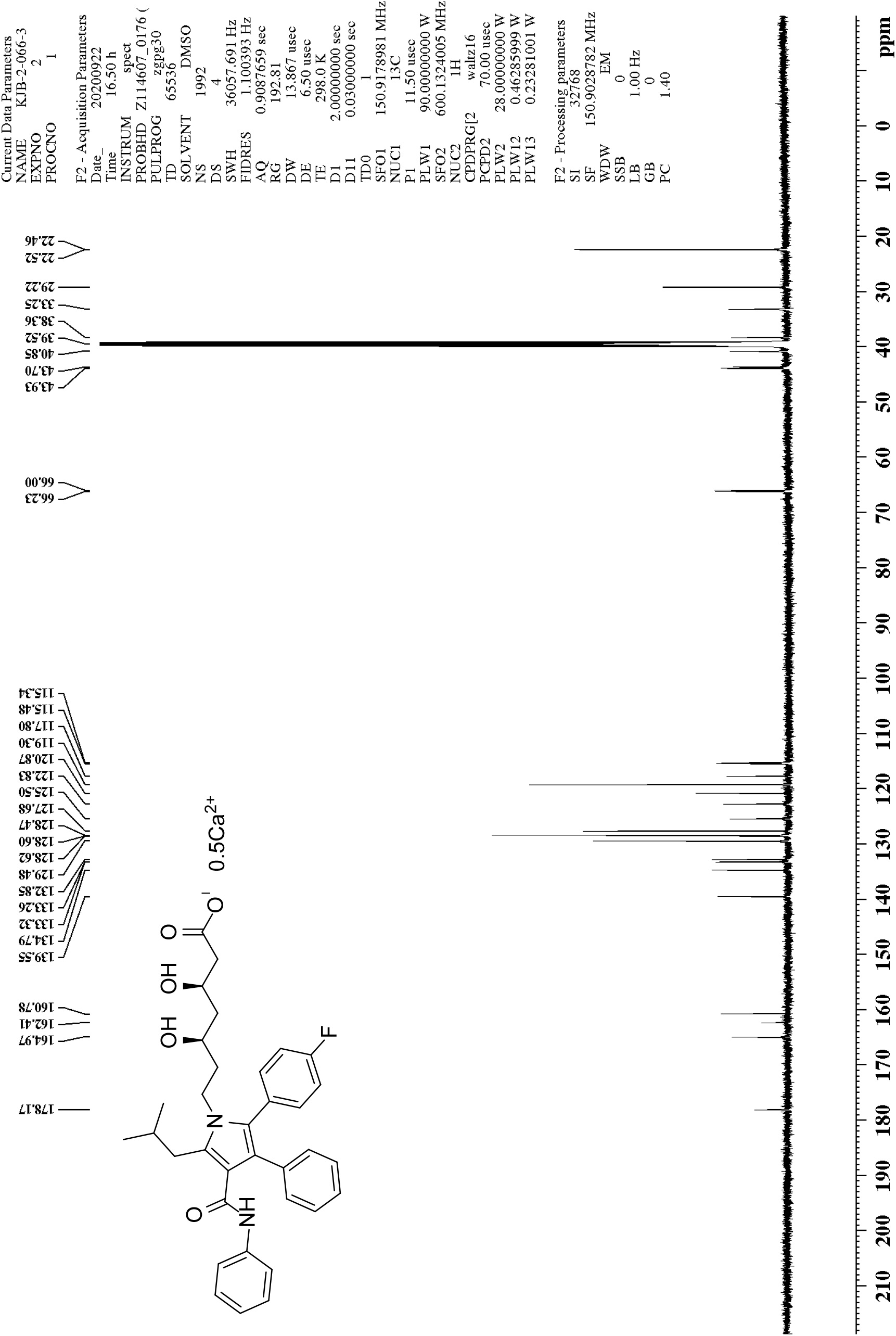
^13^C NMR of (β*R*,δ*R*)-2-(4-fluorophenyl)-β,δ-dihydroxy-5-(2-methylpropyl)-3-phenyl-4-[(phenylamino)carbonyl]-1*H*-pyrrole-1-heptanoic acid hemicalcium salt (5)

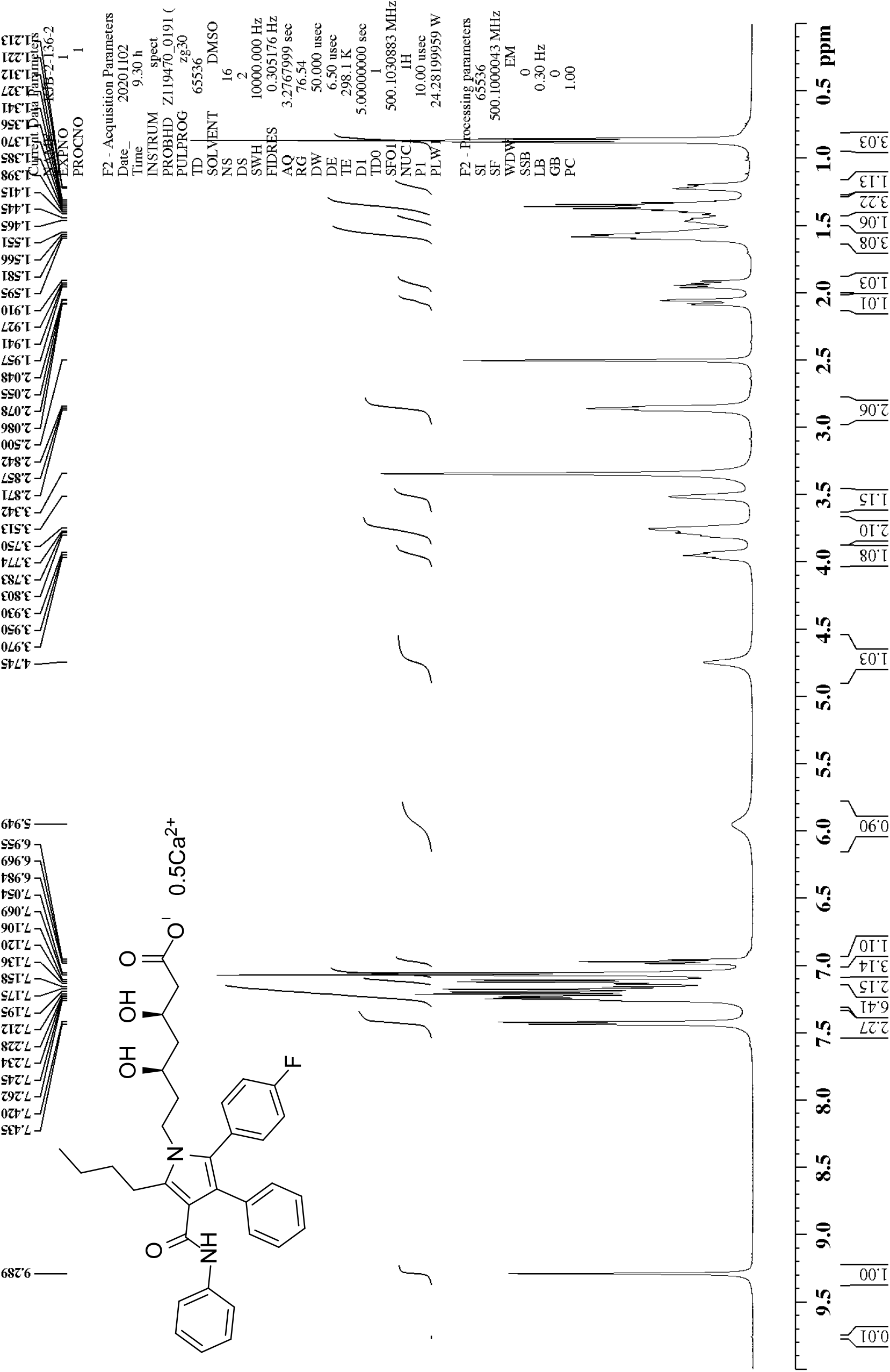
^1^H NMR of (β*R*,δ*R*)-5-butyl-2-(4-fluorophenyl)-β,δ-dihydroxy-3-phenyl-4-[(phenylamino)carbonyl]-1*H*-pyrrole-1-heptanoic acid hemicalcium salt (6)

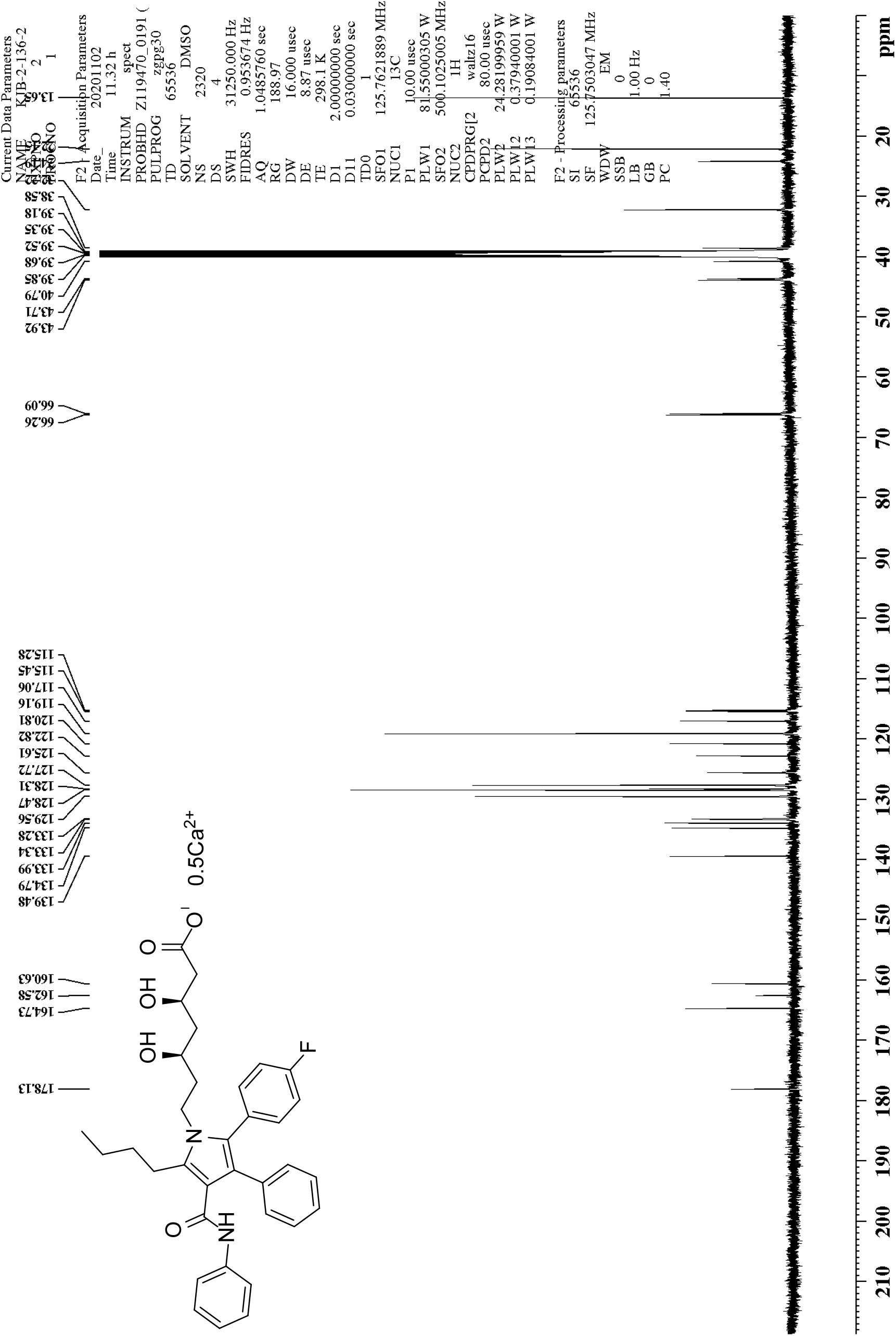
^13^C NMR of (β*R*,δ*R*)-5-butyl-2-(4-fluorophenyl)-β,δ-dihydroxy-3-phenyl-4-[(phenylamino)carbonyl]-1*H*-pyrrole-1-heptanoic acid hemicalcium salt (6)

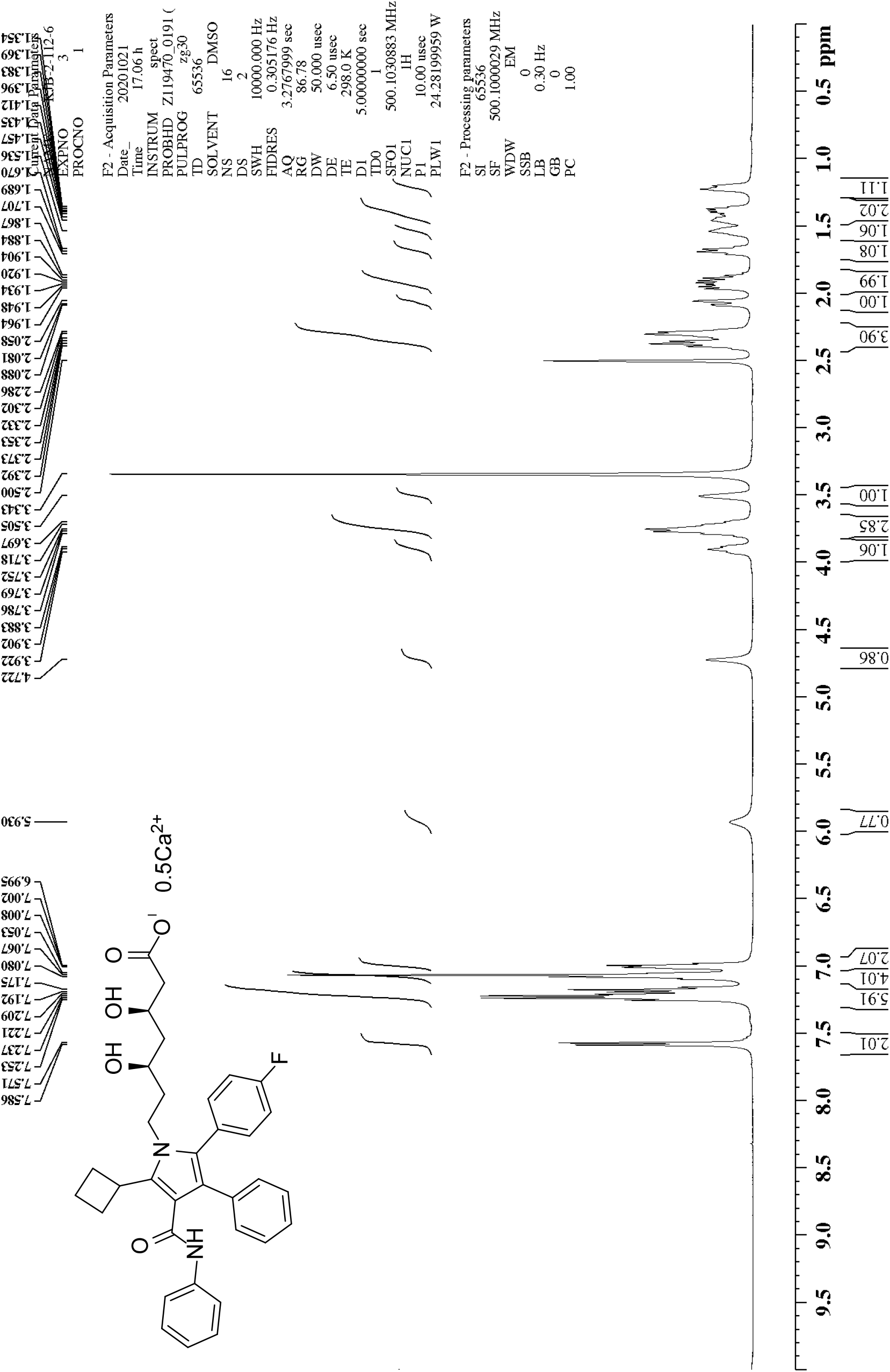
^1^H NMR of (β*R*,δ*R*)-5-cyclobutyl-2-(4-fluorophenyl)-β,δ-dihydroxy-5-cyclobutyl-3-phenyl-4-[(phenylamino)carbonyl]-1*H*-pyrrole-1-heptanoic acid hemicalcium salt (7)

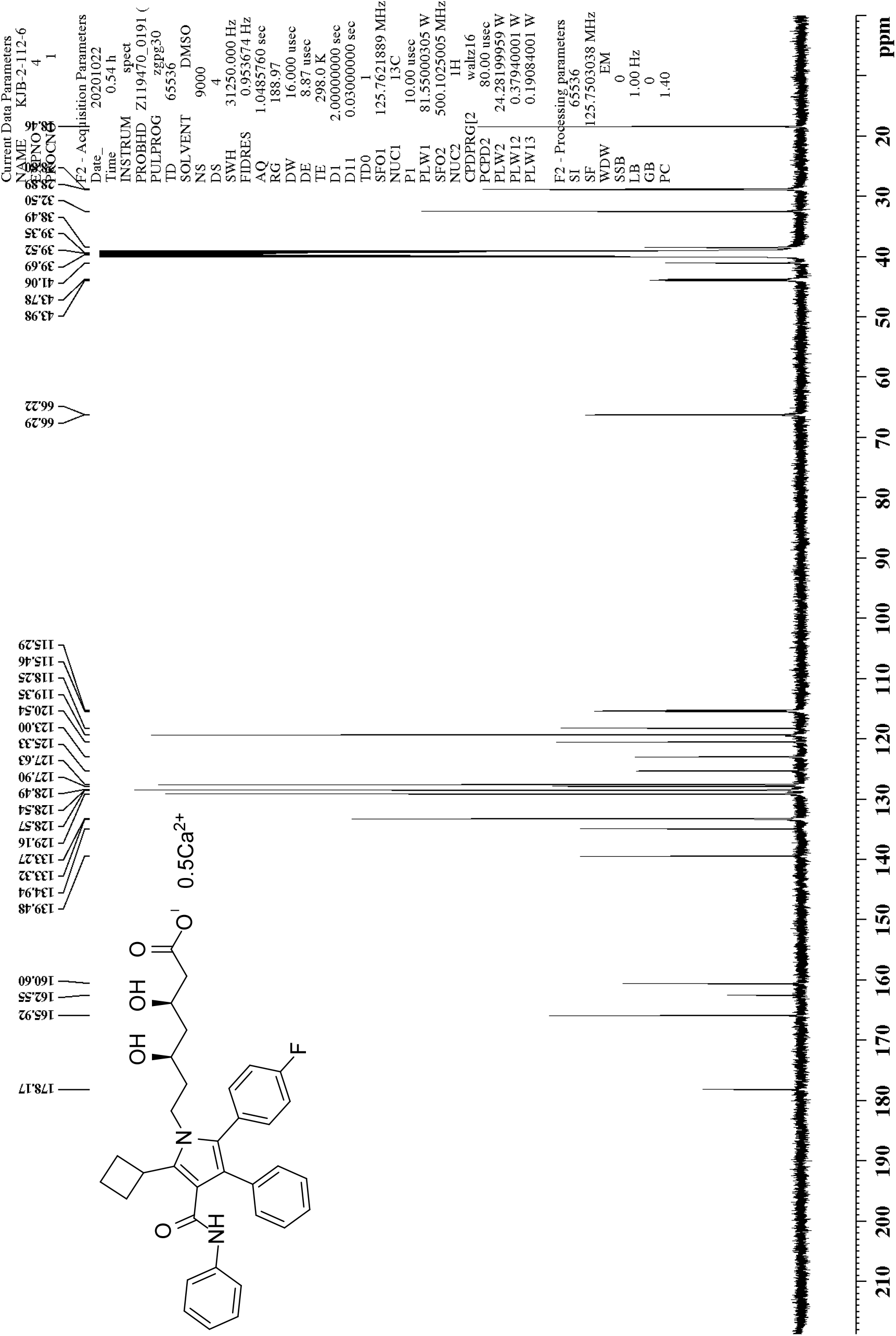
^13^C NMR of (β*R*,δ*R*)-5-cyclobutyl-2-(4-fluorophenyl)-β,δ-dihydroxy-5-cyclobutyl-3-phenyl-4-[(phenylamino)carbonyl]-1*H*-pyrrole-1-heptanoic acid hemicalcium salt (7)

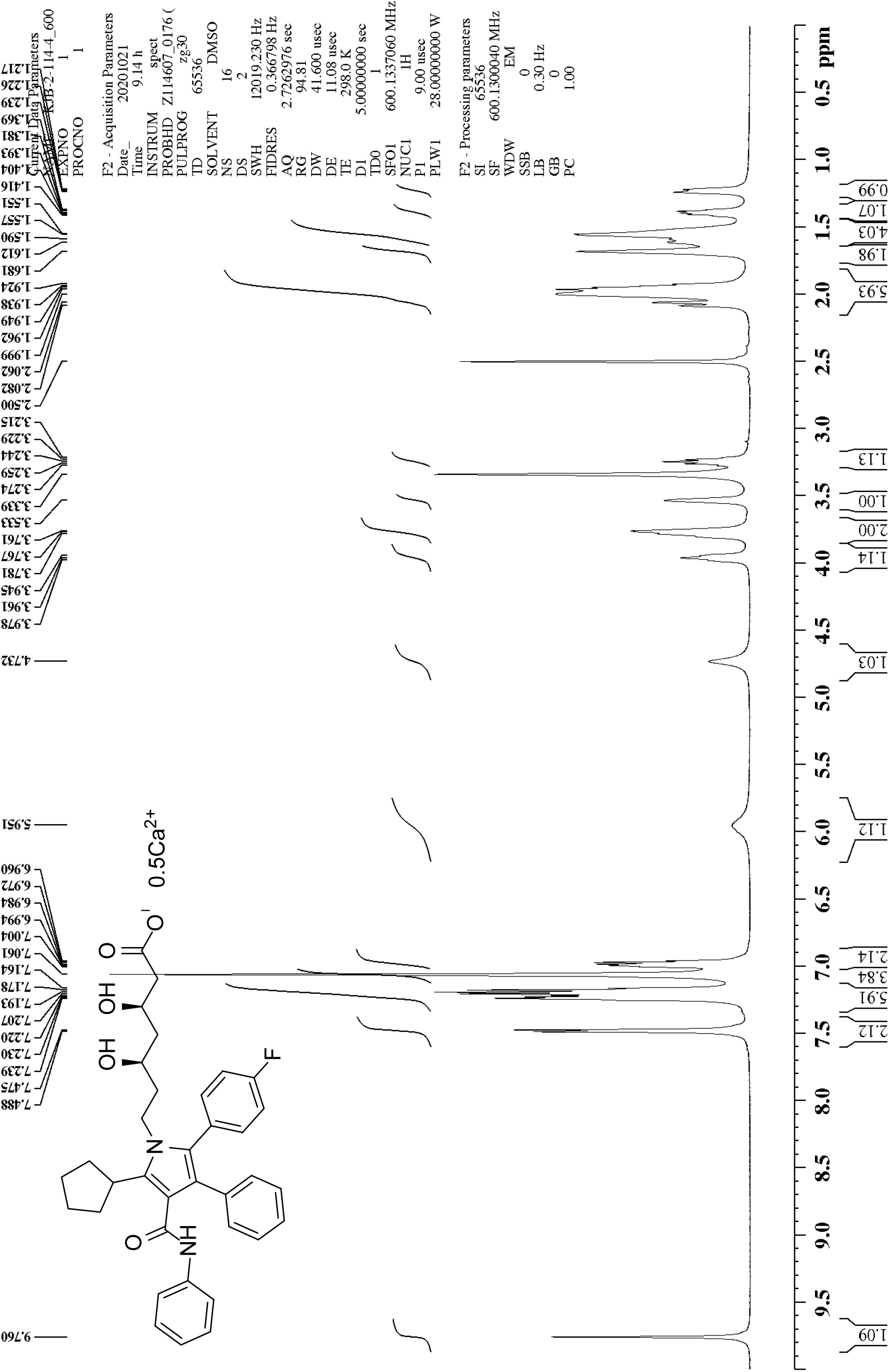
^1^H NMR of (β*R*,δ*R*)-5-cyclopentyl-2-(4-fluorophenyl)-β,δ-dihydroxy-3-phenyl-4-[(phenylamino)carbonyl]-1*H*-pyrrole-1-heptanoic acid hemicalcium salt (8)

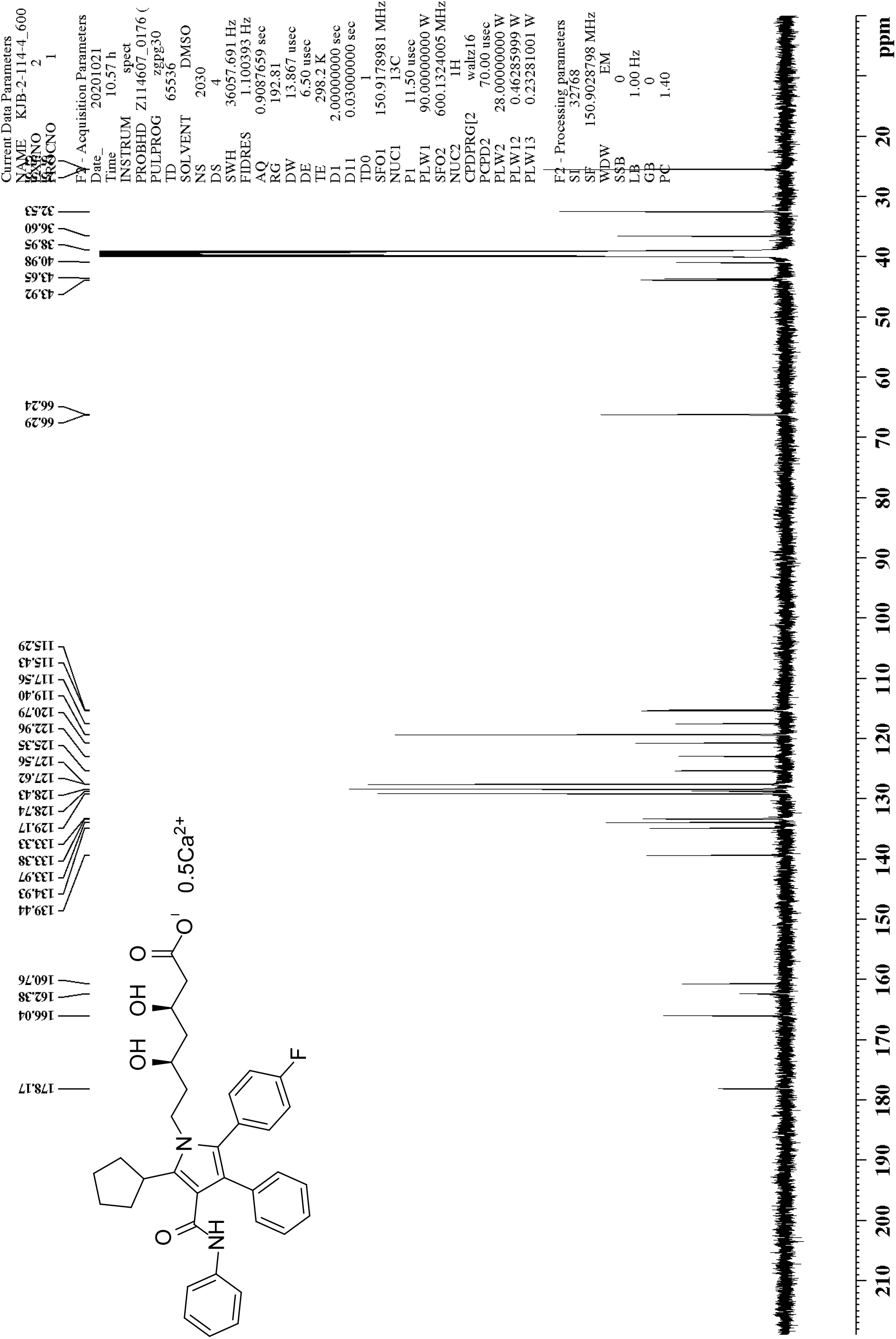
^13^C NMR of (β*R*,δ*R*)-5-cyclopentyl-2-(4-fluorophenyl)-β,δ-dihydroxy-3-phenyl-4-[(phenylamino)carbonyl]-1*H*-pyrrole-1-heptanoic acid hemicalcium salt (8)

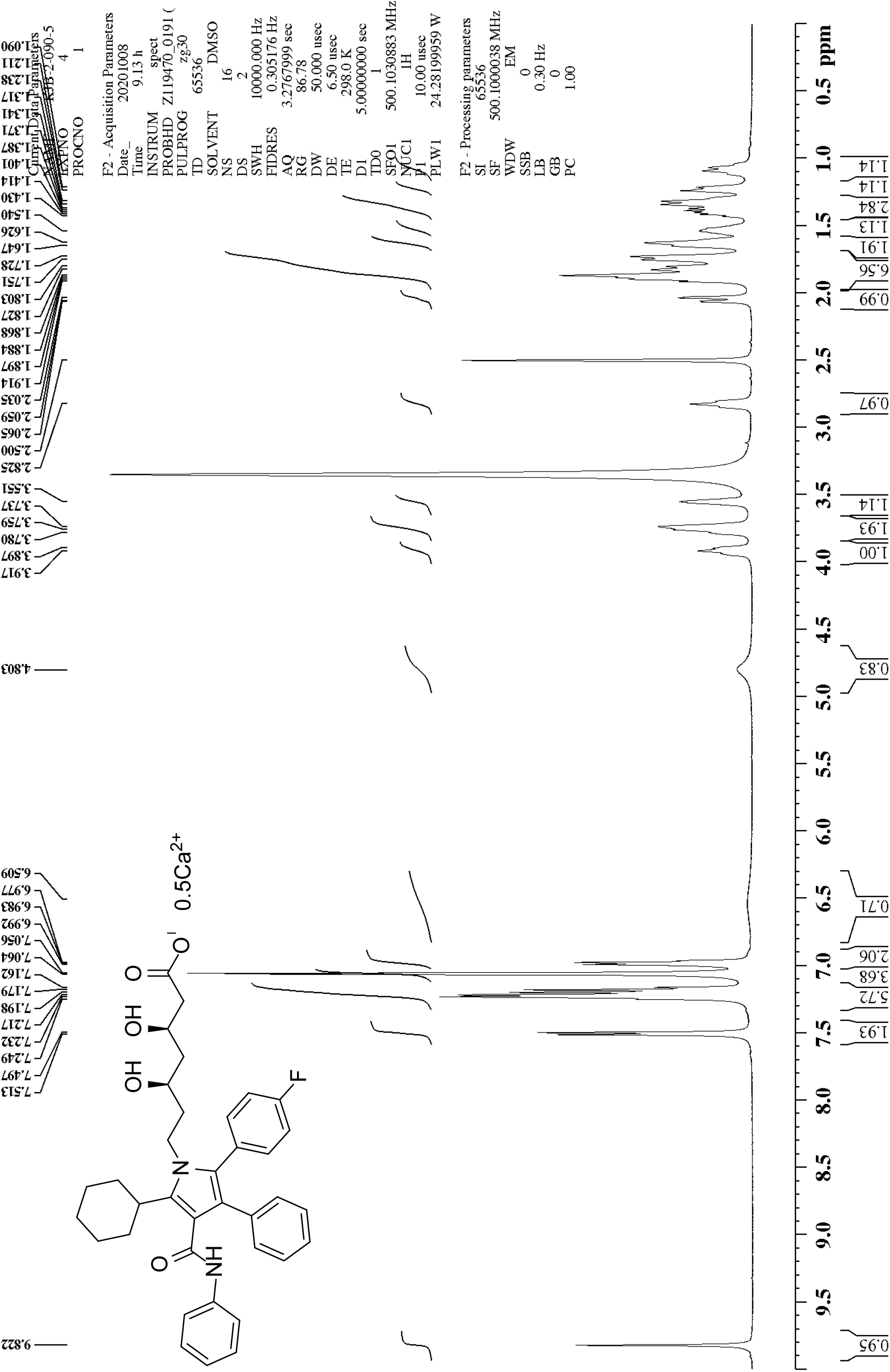
^1^H NMR of (β*R*,δ*R*)-5-cyclohexyl-2-(4-fluorophenyl)-β,δ-dihydroxy-3-phenyl-4-[(phenylamino)carbonyl]-1*H*-pyrrole-1-heptanoic acid hemicalcium salt (9)

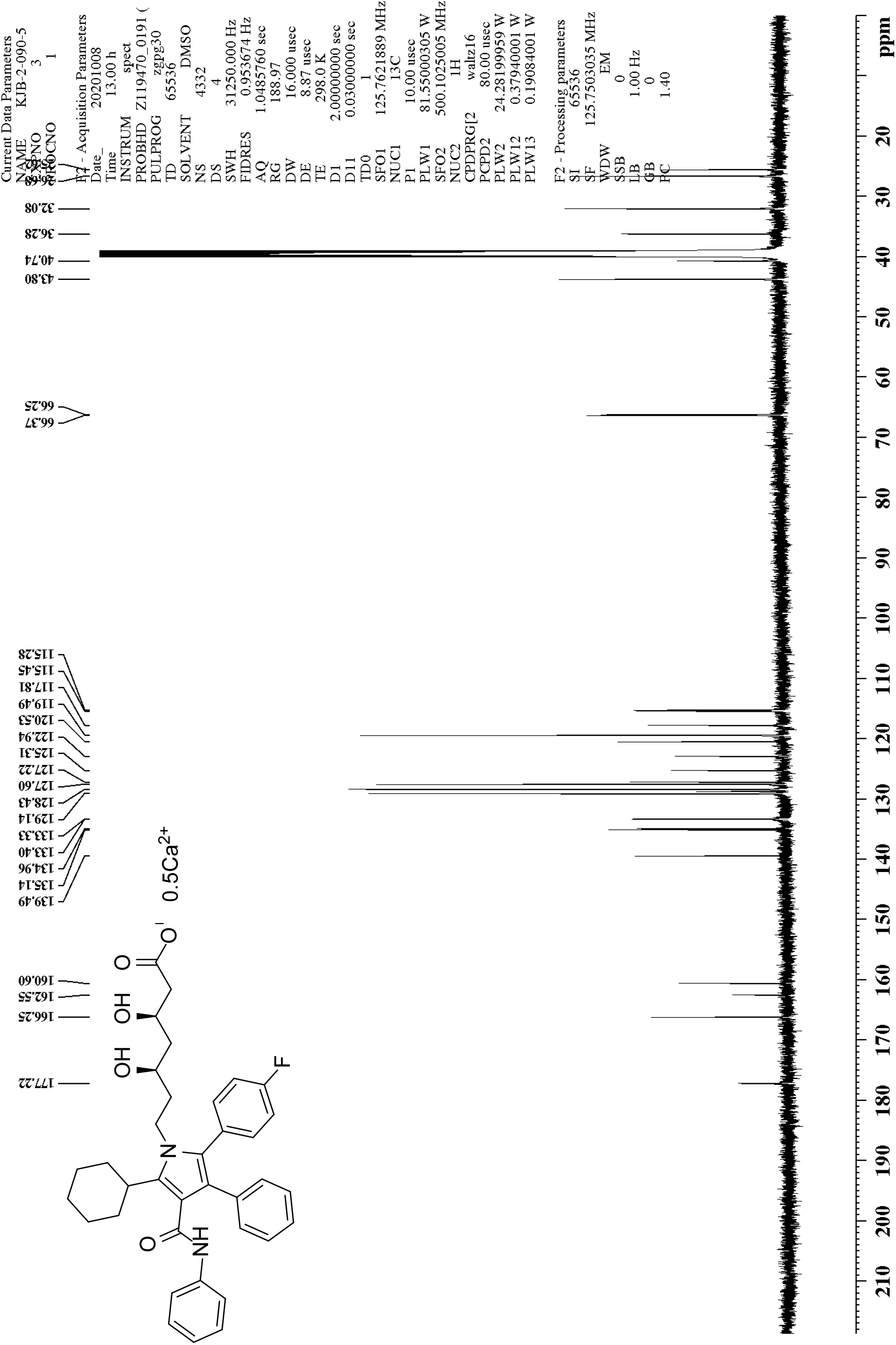
^13^C NMR of (β*R*,δ*R*)-5-cyclohexyl-2-(4-fluorophenyl)-β,δ-dihydroxy-3-phenyl-4-[(phenylamino)carbonyl]-1*H*-pyrrole-1-heptanoic acid hemicalcium salt (9)

## Footnotes

1 Xing, Y. et al. Efficient Synthesis of the Nucleus of Atorvastatin Calcium. *Synthetic Communications* **45**, 2832-2840 (2015).

2 Kawade, R.K. et al. Copper-Catalyzed Aerobic Oxidations of 3-N-Hydoxyaminoprop-1-ynes to Form 3-Substituted 3-Amino-2-en-1-ones: Oxidative Mannich Reactions with a Skeletal Rearrangement. *Chemistry - A European Journal* **20**, 13927-13931 (2014).

3 Boyle, R.G. et al. CHK-1 Inhibitors. PCT Int. Appl. WO 2005028474 A2, 2005.

4 Yuan, Y. et al. One-Pot Synthesis of 3-Hydroxyquinolin-2(1*H*)-ones from *N*-Phenylacetoacetamide via PhI(OCOCF_3_)_2_-Mediated α-Hydroxylation and H_2_SO_4_-Promoted Intramolecular Cyclization. *Journal of Organic Chemistry* **78**, 5385-5392 (2013).

5 Xing, Y. et al. Efficient Synthesis of the Nucleus of Atorvastatin Calcium. *Synthetic Communications* **45**, 2832-2840 (2015).

6 Naidu, A.A. and Sharma, G.V.R. Synthesis of novel impurities in 2-(2-(4-fluorophenyl)-2-oxo-1-phenylethyl)-4-methyl-3-oxo-N-phenylpentanamide; an atorvastatin intermediate. *Organic Communications* **10**, 314-322 (2017).

7 Sattigeri, J.A. et al. Process for preparation of (3R, 5R)-7-[2-(4-fluorophenyl)-5-isopropyl-3-phenyl-4-[(4-hydroxy methyl phenyl amino) carbonyl]-pyrrol-1-yl]-3,5-dihydroxy-heptanoic acid hemi calcium salt. PCT Int. Appl. WO 2007054790 A1, 2007.

8 Estévez, V. et al. Concise synthesis of atorvastatin lactone under high-speed vibration milling conditions. *Organic Chemistry Frontiers* **1**, 458-463 (2014).

9 Sattigeri, J.A. et al. Process for preparation of (3R, 5R)-7-[2-(4-fluorophenyl)-5-isopropyl-3-phenyl-4-[(4-hydroxy methyl phenyl amino) carbonyl]-pyrrol-1-yl]-3,5-dihydroxy-heptanoic acid hemi calcium salt. PCT Int. Appl. WO 2007054790 A1, 2007.

